# Mendelian pathway analysis of laboratory traits reveals distinct roles for ciliary subcompartments in common disease pathogenesis

**DOI:** 10.1101/2020.08.31.275685

**Authors:** Theodore George Drivas, Anastasia Lucas, Xinyuan Zhang, Marylyn DeRiggi Ritchie

## Abstract

Rare monogenic disorders of the primary cilium, termed ciliopathies, are characterized by extreme presentations of otherwise-common diseases, such as diabetes, hepatic fibrosis, and kidney failure. However, despite a revolution in our understanding of the cilium’s role in rare disease pathogenesis, the organelle’s contribution to common disease remains largely unknown. We hypothesized that common genetic variants affecting Mendelian ciliopathy genes might also contribute to common complex diseases pathogenesis more generally. To address this question, we performed association studies of 16,875 common genetic variants across 122 well-characterized ciliary genes with 12 quantitative laboratory traits characteristic of ciliopathy syndromes in 378,213 European-ancestry individuals in the UK BioBank. We incorporated tissue-specific gene expression analysis, expression quantitative trait loci (eQTL) and Mendelian disease information into our analysis, and replicated findings in meta-analysis to increase our confidence in observed associations between ciliary genes and human phenotypes. 73 statistically-significant gene-trait associations were identified across 34 of the 122 ciliary genes that we examined (including 8 novel, replicating associations). With few exceptions, these ciliary genes were found to be widely expressed in human tissues relevant to the phenotypes being studied, and our eQTL analysis revealed strong evidence for correlation between ciliary gene expression levels and patient phenotypes. Perhaps most interestingly our analysis identified different ciliary subcompartments as being specifically associated with distinct sets of patient phenotypes, offering a number of testable hypotheses regarding the cilium’s role in common complex disease. Taken together, our data demonstrate the utility of a Mendelian pathway-based approach to genomic association studies, and challenge the widely-held belief that the cilium is an organelle important mainly in development and in rare syndromic disease pathogenesis. The continued application of techniques similar to those described here to other phenotypes/Mendelian diseases is likely to yield many additional fascinating associations that will begin to integrate the fields of common and rare disease genetics, and provide insight into the pathophysiology of human diseases of immense public health burden.

## INTRODUCTION

Found on nearly every human cell type, the primary cilium is a small, non-motile projection of the cell’s apical surface that acts as an antenna for the reception and integration of signals from the extracellular environment.^1^ Signaling at the cilium is mediated by cell surface receptors that are specifically trafficked to the cilium through a complex and highly regulated series of interactions.^2,3^ Receptors destined for the ciliary space must first interact with a protein subcomplex known as the BBSome, which allows receptors to dock at the base of the cilium (the basal body) and navigate through the diffusion barrier formed by the ciliary transition zone to enter the inner ciliary space.^4,5^ Once inside, cargo is specifically trafficked along the microtubule core of the cilium through a process known as intraflagellar transport (IFT), before being ultimately returned to the extraciliary space to complete signal transduction.^6^

We now know that the interruption of any one of these critical ciliary pathways can lead to the development of devastating syndromic Mendelian disorders, collectively known as ciliopathies.^7^ The list of confirmed ciliopathies is constantly expanding, and currently includes disorders such as Alström syndrome, Bardet-Biedl syndrome, Joubert syndrome, Meckel-Gruber syndrome, Nephronophthisis, Senior–Løken syndrome, Sensenbrenner syndrome, and others.^8–16^ Much of the pathology of the ciliopathy syndromes can be attributed to the adverse effects of ciliary dysfunction on cell signaling pathways critical to embryologic development.^7,17^ Furthermore, although each ciliopathy is defined by its own unique constellation of characteristic phenotypic findings, many ciliopathies share similar or overlapping features.

Phenotypes that are often observed in individuals with ciliary disease include renal failure, hepatic fibrosis, obesity, dyslipidemia, and diabetes.^8–10^ Interestingly, these very same phenotypes are characteristic of common diseases that affect large proportions of the population. Recent evidence suggests that the development of these phenotypes in ciliopathy patients may be driven by the perturbation of ciliary processes required for signaling through common disease-relevant pathways not previously known to be dependent on the cilium for signal transduction, such as insulin, IGF-1, PDGF, and TGFβ.^2,18–28^ Together, these observations raise the possibility that the same ciliary genes causative of rare Mendelian disease may also be involved in the pathogenesis of common complex diseases, such as kidney failure or dyslipidemia, in the general population.

To investigate this hypothesis, we set out to identify associations between common genetic variants in 122 well-characterized ciliary genes and 12 quantitative laboratory traits relevant to common disease in a large biobank cohort. We incorporated tissue-specific gene expression analysis, expression quantitative trait loci (eQTL), and Mendelian disease information into our analysis, and replicated findings in meta-analysis to increase our confidence in observed associations between ciliary genes and human phenotypes (Figure 1). The results of our analysis revealed 73 statistically significant associations between 34 ciliary genes and diverse laboratory traits (including 8 novel, replicating associations), and also identified distinct ciliary subcompartments associated with different disease processes and demonstrated pleiotropic effects for many ciliary genes across a number of phenotypes and organ systems. Our data challenge the widely-held belief that the cilium is an organelle important mainly in development and in rare syndromic disease pathogenesis, and establish a framework for the investigation of the common disease contributions of other Mendelian disease pathways.

**Figure 1.**
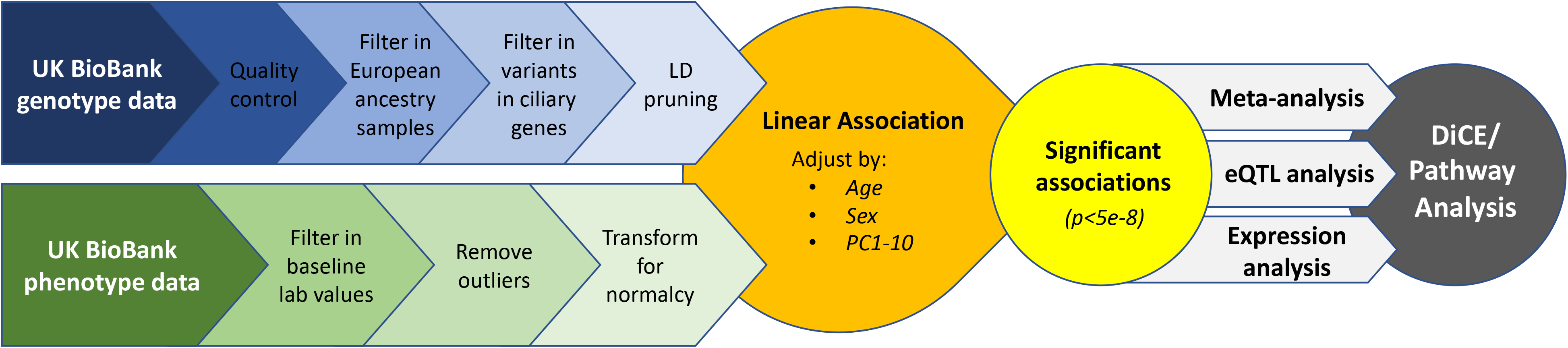
Schematic of analysis workflow for the data presented. Phenotype and genotype data from the UKBB release version 2 was extracted and processed as indicated. Linear association studies between ciliary gene variants and patient phenotypes was performed, with genome-wide significant associations (p <5e-8) being studied further by tissue-specific expression, eQTL, and replication meta-analysis. Using this data, a DiCE/Pathway analysis was performed to identify ciliary subcompartments associated with specific traits.

## RESULTS

### 34 ciliary gene loci are associated with diverse quantitative laboratory traits

We examined 16,875 genotyped or imputed variants within genomic loci defined by the boundaries of 122 well-defined ciliary genes in 378,213 European ancestry individuals in the UK BioBank. We tested each variant for association with each of 12 quantitative laboratory traits: Alanine aminotransferase (ALT), Alkaline Phosphatase (AlkPhos), Aspartate aminotransferase (AST), Total Cholesterol (Cholesterol), Serum Creatinine (Creatinine), Gamma glutamyltransferase (GGT), Serum Glucose (Glucose), Glycated haemoglobin (A1c), HDL cholesterol (HDL), LDL cholesterol (LDL), Triglycerides, and Urea using linear regression, adjusting our model for age, sex, and ancestry PCs 1-10. ALT, AlkPhos, AST, and GGT are all markers of liver disease, Creatinine and Urea are markers of kidney disease, perturbations in HDL, LDL, Total Cholesterol, or Triglyceride levels are the hallmark of dyslipidemia, while elevations in Glucose or A1c levels can be diagnostic of diabetes and impaired glucose homeostasis. The characteristics of the subjects and phenotypes included in the discovery analysis are summarized in Table S1. To replicate novel findings, large scale meta-analyses of the same 12 quantitative laboratory traits were carried out, as described in the methods section, for genetic variants contained within the same 122 ciliary genes as analyzed in our discovery set. Results of our discovery analysis and replication meta-analyses are displayed in Figures 2-5.

**Figure 2.**
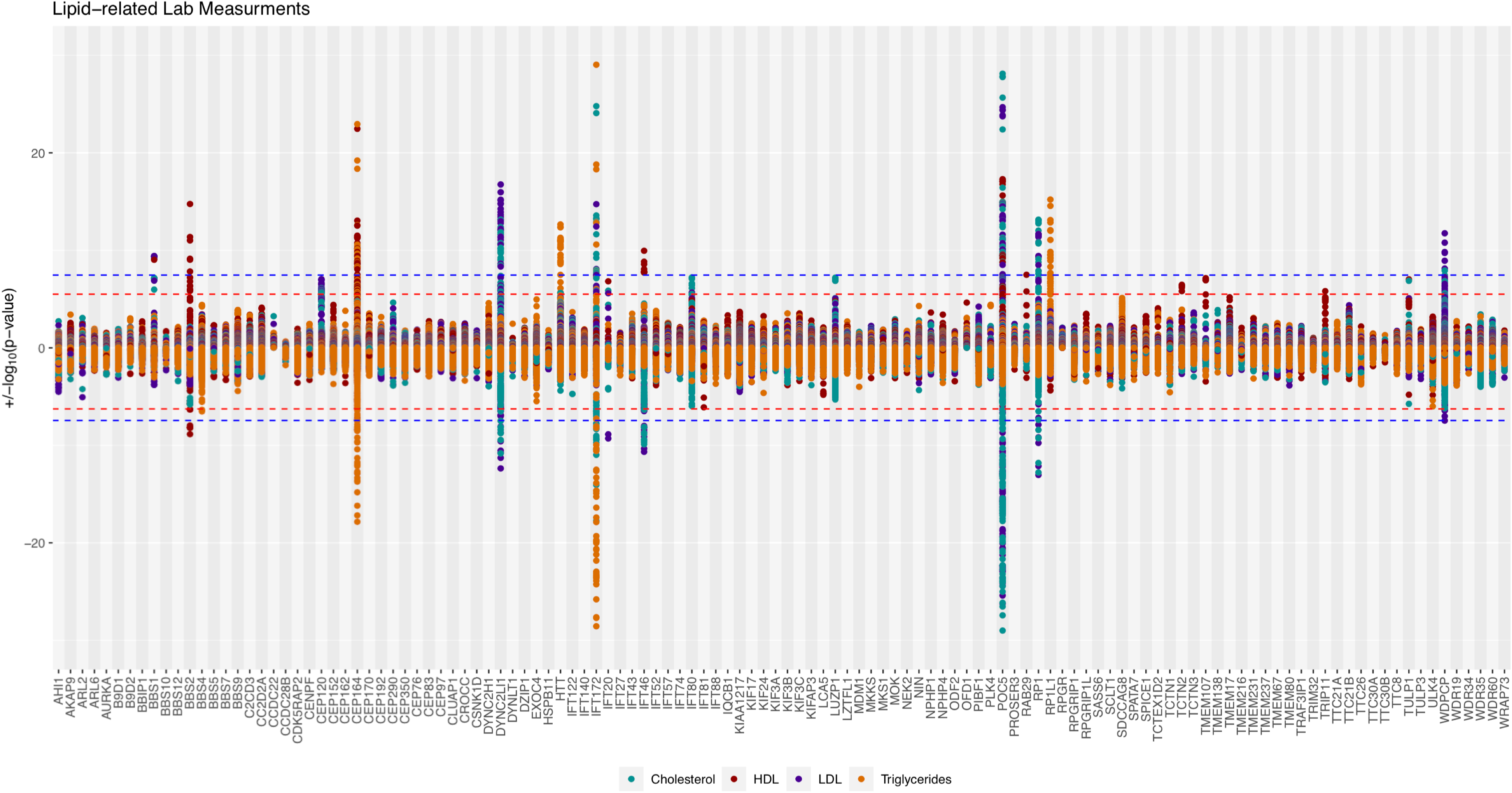
Association study and meta-analysis results for lipid-related traits. Hudson plot illustrating the results of the discovery association analysis (top) and replication meta-analysis (bottom) of common variants within 122 ciliary gene with lipid-related traits. Data for cholesterol is in teal, HDL in red, LDL in purple, and triglycerides in orange. The study-wide Bonferroni-adjusted significance threshold (p<3.4e-8) is shown as a blue dashed line, while the experiment-wide Bonferroni-adjusted significance threshold (p<3.0e-6 for the discovery analysis, p<3.3e-6 for the meta-analysis) is shown as a red dashed line. Each analyzed gene is given equal space along the horizontal axis, with all tested genetic variants for a given gene plotted at the midline of the gene block.

Our discovery analysis identified 549 variants within 34 different ciliary genes reaching genome-wide significance (p < 5e-8) for association with at least one of the 12 phenotypes (Figure 2-5, Table S2-13). Many ciliary gene loci were significantly associated with multiple traits, a phenomenon known as pleiotropy (with one locus, containing the gene *IFT172*, significantly associated with nine different traits). 73 significant gene-trait associations were identified altogether, and of these 27 (37%) represent replication of previously-known associations from the literature, whereas 46 (63%) represent previously-unreported associations (novel findings). 8 of these 46 (17%) unreported associations were found to replicate in our meta-analyses. These results are summarized in Table S14. Of the 8 replicating novel associations, five were for kidney-related phenotypes, which is likely reflective of the fact that these phenotypes had the largest samples sizes in our meta-analysis (the meta-analysis of kidney related traits had a sample size of ~800,000 individuals, whereas no other meta-analyzed trait had a sample size greater than 370,000). All replicating novel associations and interesting known associations are discussed in more detail below.

### Tissue-specific analysis demonstrates widespread expression of significant ciliary genes

For all genes with significant associations in our discovery analysis, we examined tissue-specific expression in the GTEx version 8 database database^29^ to determine if each gene was expressed in tissues relevant to the detected phenotype association (Figure 6, Table S15). As the cilium is a nearly ubiquitous organelle,^30^ we were not surprised to find that the majority of ciliary genes examined (20 of 34, 59%) were broadly expressed in all tissues (Figure 6). Four genes, however, stood out as being poorly expressed in all or nearly all examined tissues – *CENPF*, *RP1*, *RP1L1*, and *WDPCP*. Why these four specific ciliary genes were the only ones found to be poorly expressed is not entirely clear.

### Trait-significant variants are enriched for and correlate with eQTLs of ciliary genes

The majority of the 549 ciliary gene variants statistically significantly associated with laboratory traits lay within untranslated regions of the genome. To explore the possibility that these variants might be exerting their effect on laboratory traits by modulating ciliary gene expression, we performed an eQTL analysis of each of the 73 significant gene-trait pairs (Figures S1-S34). Utilizing the GTEx version 8 database of cis-eQTL variants,^29^ we plotted the association peaks for each significant gene-trait pair in chromosomal space, overlaying the relevant eQTL information for each ciliary candidate gene using eQTpLot.^31,32^ We simultaneously generated plots illustrating enrichment of ciliary candidate gene eQTLs among trait-significant variants, and generated p-p plots (plotting, for each variant, the p-value of association with candidate gene expression (p_eQTL_) against the p-value of association with laboratory trait level(p_trait_)) to illustrate and identify significant correlations between variants’ effects on candidate gene expression and their effects on laboratory traits, as determined by linear regression and calculation of the Pearson correlation coefficient and p-value of correlation.

Interestingly, we found that for 70 of the 73 (96%) significant gene-trait pairs, the detected association peaks were significantly enriched for eQTLs of the candidate ciliary gene (p< 7e-4). Furthermore, for 48 of the 70 (69%) significant gene-trait pairs that demonstrated eQTL enrichment, there was a significant correlation between p_trait_ and p_eQTL_ (Pearson correlation coefficient >0.25, p-value of correlation <3.5e-4). These data indicate that the majority of association signals are being driven by genetic variants that are similarly associated with significant changes in expression of candidate ciliary genes. Together, these findings are consistent with the hypothesis that the detected laboratory trait associations are driven by changes in ciliary gene expression.

### Novel Associations

Our discovery analysis identified 46 previously-unreported genome-wide significant associations, 8 of which were found to replicate in our meta-analyses at either the variant or gene level:

#### BBS2

We found a significant association between variants within the *BBS2* gene and HDL cholesterol levels (most significant association with rs118024138, p-value 5.73e-37). This same locus was found to be significantly associated with HDL in our meta-analysis (Figure 2). *BBS2* encodes a BBSome-associated protein critical in the regulation of cargo transport through the ciliary space.^33^ Biallelic pathogenic variants in the gene have been associated with Bardet-Biedl syndrome, characterized by retinal degeneration, kidney failure, obesity, impaired glucose handling, and, interestingly, dyslipidemia.^8,34,35^ Furthermore, *BBS2* was found to be significantly expressed in both liver and adipose tissue (Figure 6), with a majority of variants significantly associated with HDL levels also found to be significantly associated with *BBS2* expression in the GTEx database (Figure S3), lending a number of lines of supporting evidence to this novel association.

#### CLUAP1

A significant association was detected between variants within the *CLUAP1* gene and serum creatinine levels (most significant association for variant rs9790, p-value 2.56e-12), with variant-level replication of this finding in our meta-analysis (Figure 3). Although *CLUAP1* exists in a gene-dense region of the genome, the peak of association was confined specifically to within the *CLUAP1* gene boundaries (Figure S8). We found *CLUAP1* to be significantly expressed in kidney tissues (Figure 6), with a majority of the variants significantly associated with creatinine also significantly associated with *CLUAP1* expression levels in the GTEx eQTL database (Figure S8). *CLUAP1* encodes the intraflagellar transport (IFT)-associated protein CLUAP1/IFT38, a component of the ciliary IFT-B complex important for anterograde transport of cargo along the ciliary axoneme. Biallelic pathogenic variants in the gene have been associated with an overlap syndrome similar to Joubert syndrome and Orofaciodigital syndrome in one individual.^36^ While this individual was not found to have evidence of renal disease, renal disease is a common feature of Joubert syndrome patients more generally.^9^ Interestingly, this individual was also noted to be obese (a common feature of ciliopathy syndromes); the *CLUAP1* locus has been published as associated with BMI and type 2 diabetes in previous GWAS studies.^37–40^

**Figure 3.**
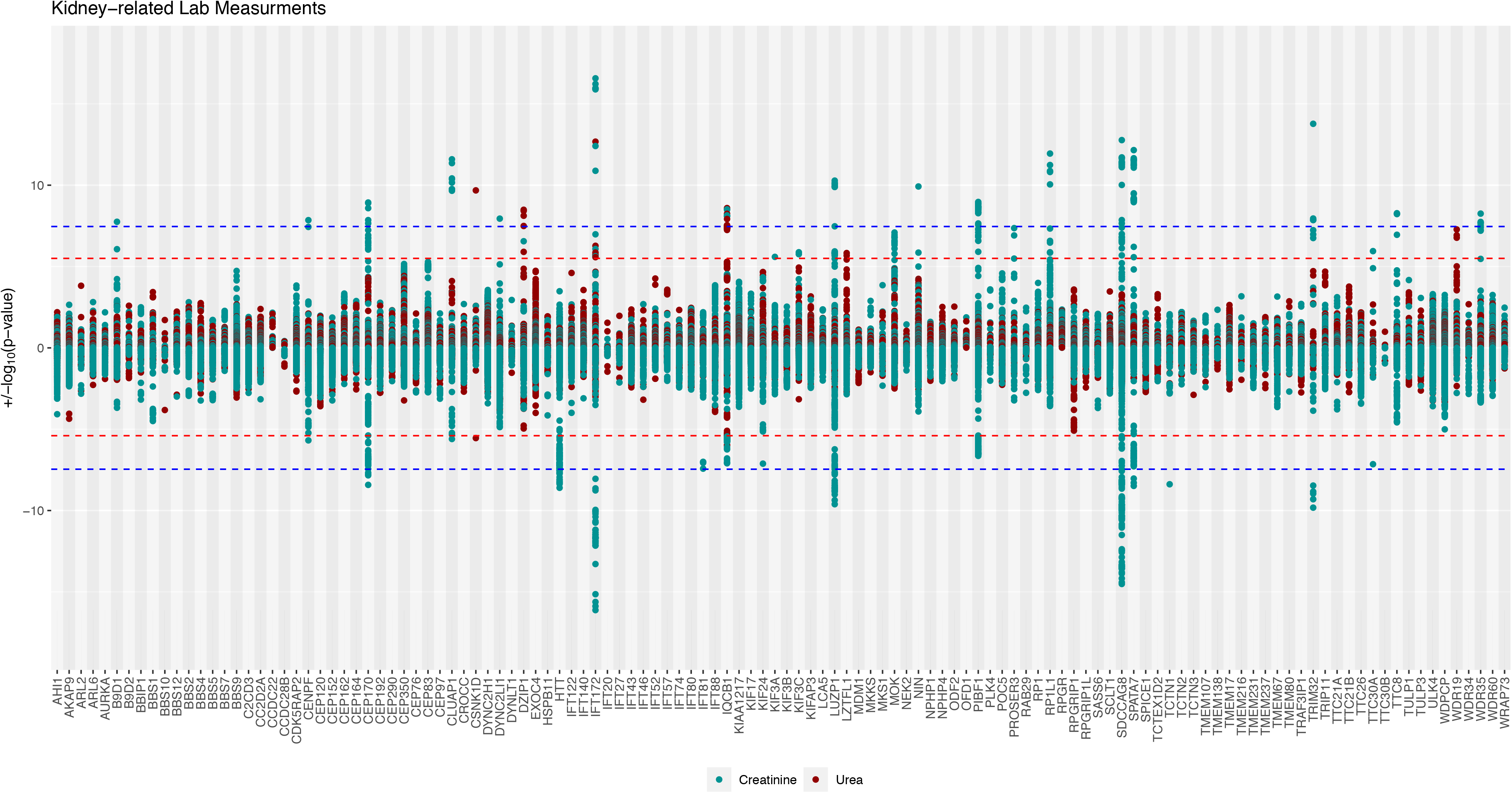
Association study and meta-analysis results for kidney-related traits. Hudson plot illustrating the results of the discovery association analysis (top) and replication meta-analysis (bottom) of common variants within 122 ciliary gene with kidney-related traits. Data for creatinine is in teal, and data for urea is in red. The study-wide Bonferroni-adjusted significance threshold (p<3.4e-8) is shown as a blue dashed line, while the experiment-wide Bonferroni-adjusted significance threshold (p<3.0e-6 for the discovery analysis, p<3.8e-6 for the meta-analysis) is shown as a red dashed line. Each analyzed gene is given equal space along the horizontal axis, with all tested genetic variants for a given gene plotted at the midline of the gene block.

#### CSNK1D

A single variant within the *CSNK1D* gene, rs11653735, was found to be significantly associated with urea levels with a p-value of 2.08e-10, with this same variant found to replicate in our meta-analysis (Figure 3). *CSNK1D* encodes the protein kinase CK1δ that has been found to play a critical role in regulating ciliogenesis.^41^ Variants in *CSNK1D* have not been implicated in any classic ciliopathy syndrome. The rs11653735 variant is a deep intronic variant, and if/how it affects CK1δ function is unclear. Intriguingly, the genomic region surrounding *CSNK1D* contains the gene *CCDC57*, a poorly characterized gene associated with the centriole and cilium that was not included in our study.^42^ The peak of association for urea levels spans the entirety of the *CCDC57* gene (Figure S9), with a strong correlation between variants associated with urea and those associated with *CCDC57* expression (Figure S35), raising the possibility that this locus is associated with urea levels through perturbation of either or both of these ciliary genes, *CCDC57* and *CSNK1D*.

#### IQCB1

We found a number of variants within and surrounding the *IQCB1* locus to be significantly associated with both serum creatinine (a known association, most significant p-value of association in our discovery analysis 3.05e-09 for SNP rs9823335) and urea levels (a novel association, most significant p-value of association in our discovery analysis 2.56e-09 for rs75382826). Both of these associations were found to replicate at the variant and gene level in our meta-analysis (Figure 3). *IQCB1* is significantly expressed in the kidney (Figure 6), and nearly every variant in the *IQCB1* locus that was found to be significantly associated with urea/creatinine levels was also found to be significantly associated with *IQCB1* expression, with strong evidence for correlation between levels of *IQCB1* expression and creatinine/urea levels (Figure S16). *IQCB1* encodes a well-characterized protein of the same name that is a critical component of the ciliary transition zone. Rare pathogenic biallelic variants in *IQCB1* are a major cause of Senior-Løken syndrome, a disorder characterized by retinal degeneration and progressive kidney failure.^14^ The novel association of the *IQCB1* locus with blood urea levels corroborates the known association with creatinine/glomerular filtration rate, and adds further evidence supporting the role of *IQCB1* in kidney function more generally.

#### KIF17

We identified a very strong novel association between variants within the *KIF17* locus with AlkPhos levels (most significant p-value of association 4.09e-47 for rs61778523), which replicated at the gene level in our meta-analysis (Figure 4). Interestingly, we did not find *KIF17* to be significantly expressed in hepatic tissue (Figure 6), and there did not appear to be a significant correlation between variants associated with *KIF17* expression levels and AlkPhos levels (Figure S17). *KIF17* encodes a motor protein important in driving IFT-B-mediated anterograde ciliary transport, but no Mendelian disease has yet been associated with dysfunction of this gene.^43^ Thus altogether there is minimal corroborating evidence to support *KIF17* as the causative gene of this novel association.

**Figure 4.**
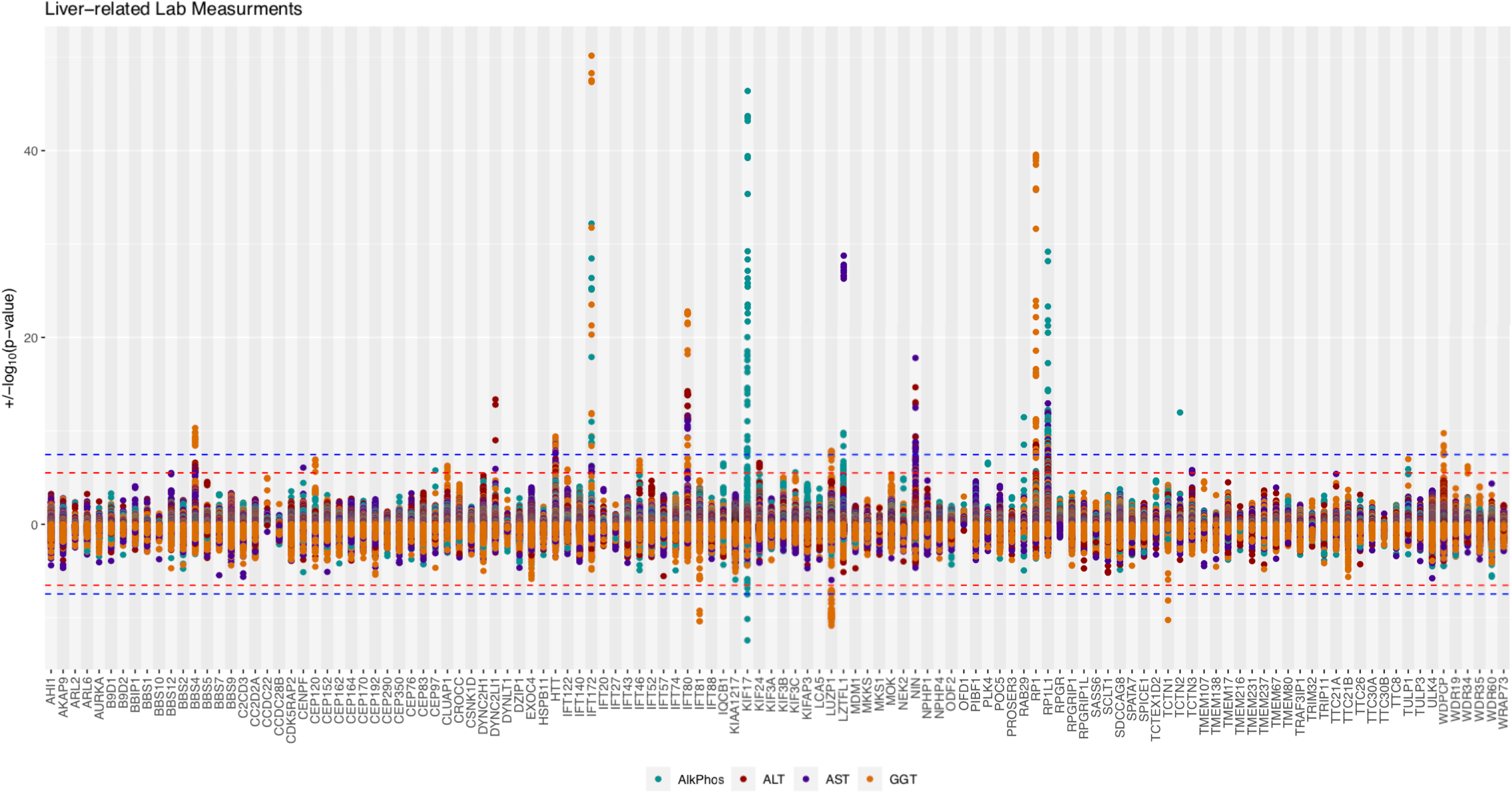
Association study and meta-analysis results for liver-related traits. Hudson plot illustrating the results of the discovery association analysis (top) and replication meta-analysis (bottom) of common variants within 122 ciliary gene with liver-related traits. Data for AlkPhos is in teal, ALT in red, AST in purple, and GGT in orange. The study-wide Bonferroni-adjusted significance threshold (p<3.4e-8) is shown as a blue dashed line, while the experiment-wide Bonferroni-adjusted significance threshold (p<3.0e-6 for the discovery analysis, p<3.3e-6 for the meta-analysis) is shown as a red dashed line. Each analyzed gene is given equal space along the horizontal axis, with all tested genetic variants for a given gene plotted at the midline of the gene block.

**Figure 5.**
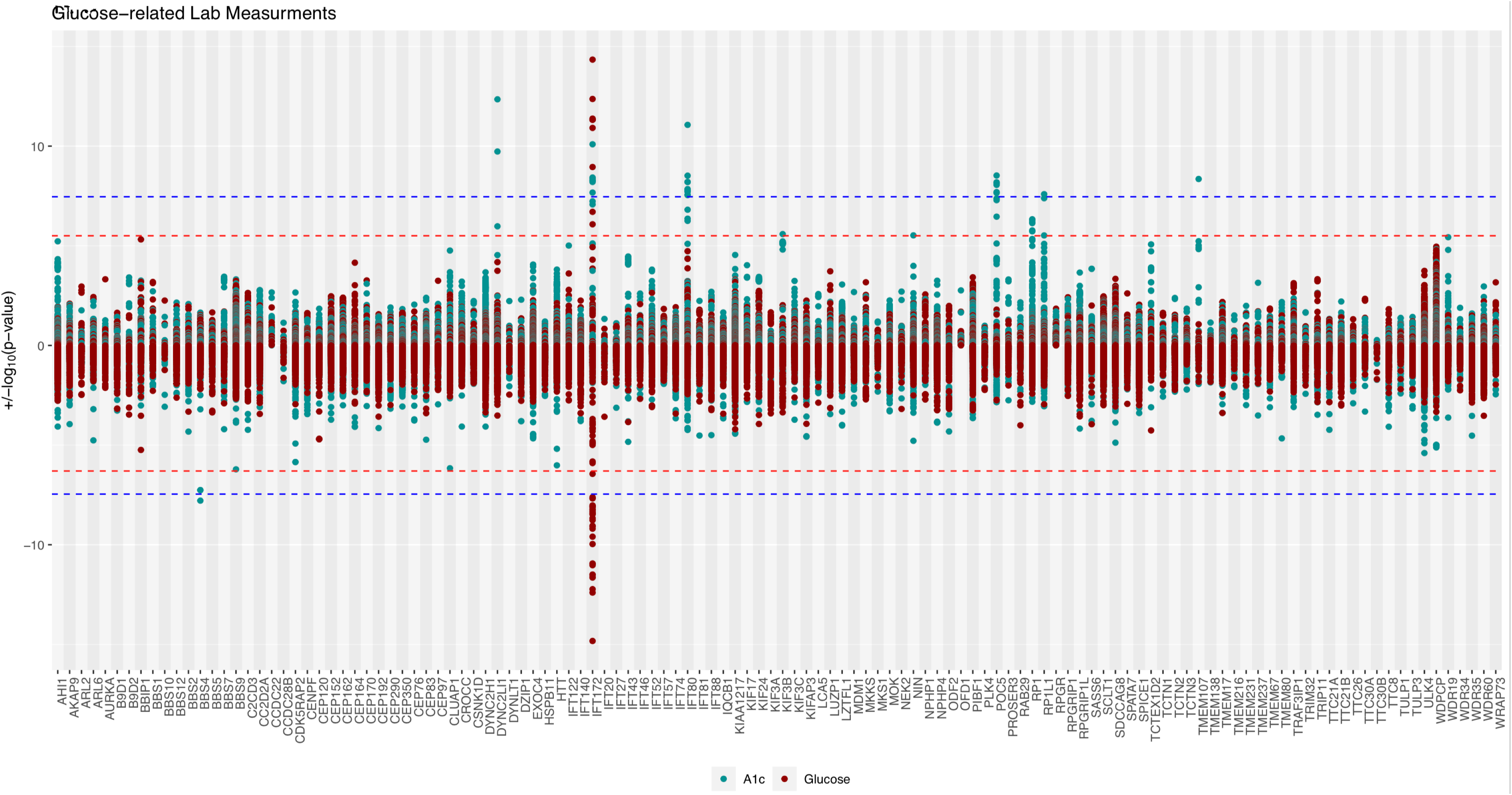
Association study and meta-analysis results for glucose-related traits. Hudson plot illustrating the results of the discovery association analysis (top) and replication meta-analysis (bottom) of common variants within 122 ciliary gene with glucose-related traits. Data for A1c is in teal, and data for glucose is in red. The study-wide Bonferroni-adjusted significance threshold (p<3.4e-8) is shown as a blue dashed line, while the experiment-wide Bonferroni-adjusted significance threshold (p<3.0e-6 for the discovery analysis, p<3.3e-6 for the meta-analysis) is shown as a red dashed line. Each analyzed gene is given equal space along the horizontal axis, with all tested genetic variants for a given gene plotted at the midline of the gene block.

**Figure 6.**
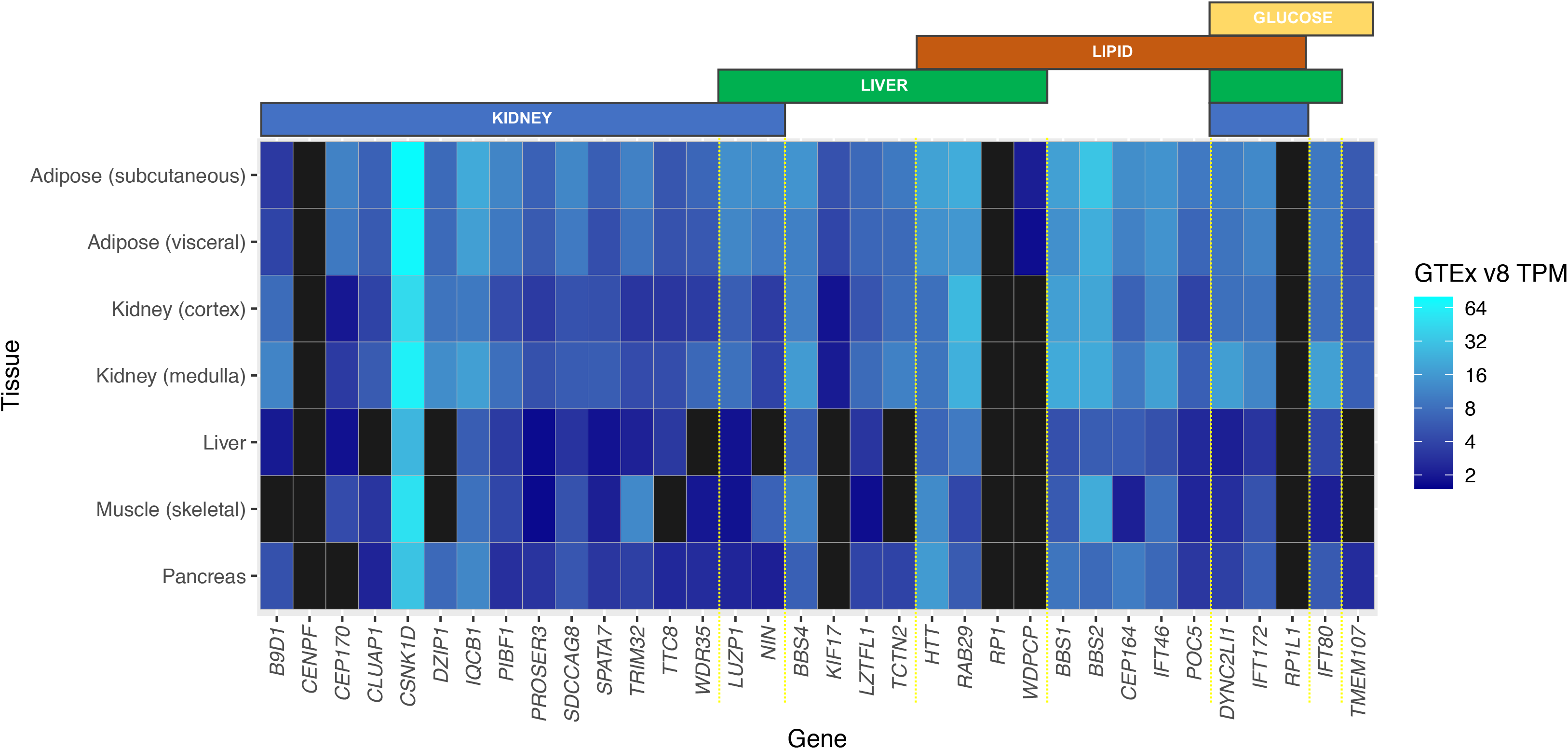
Results of tissue-specific expression analysis. Heatmap depicting the results of the tissue-specific expression analysis for each of the 34 genes with significant associations in our discovery analysis. The phenotype domain(s) significantly associated with each gene are indicated by the colored bars at the top of the graph.

#### LUZP1

We found variants within the *LUZP1* gene to be significantly associated with both GGT levels (most significant p-value 1.41e-08 for variant rs56087807) and serum creatinine levels (most significant p-value 5.23e-11 for variant rs1208930), with both of these findings replicating at both the variant and gene level in our meta-analysis (Figures 3-4). The association between common variants near the *LUZP1* locus and GGT levels has been reported,^44^ but the association with creatinine is novel. *LUZP1* was found to be significantly expressed in all tissues relevant to these phenotypes (Figure 6), and nearly every variant significantly associated with creatinine/GGT levels was also found to be significantly associated with *LUZP1* expression, with evidence for correlation between levels of *LUZP1* expression and creatinine/GGT levels (Figure S18). *LUZP1* encodes a ciliary basal body protein that has recently been shown to play important roles in regulating ciliogenesis.^45,46^ While there is no Mendelian disease yet known to result from disruption of the *LUZP1* gene, recent work has shown reduced *LUZP1* expression in fibroblasts derived from Townes-Brockes syndrome (characterized by renal disease, among other findings) patients,^47^ suggesting that *LUZP1* may play a role in the pathogenesis of this disorder.

#### PIBF1

A significant, novel association was detected between variants within the *PIBF1* gene and serum creatinine levels (most significant p-value 1.08e-09 for variant rs111440455), with this finding replicating at both the variant and gene level in our meta-analysis (Figure 3). We found *PIBF1* to be significantly expressed in the kidney (Figure 6), with nearly every variant significantly associated with creatinine levels also found to be significantly associated with *PIBF1* expression, with strong evidence for correlation between levels of *PIBF1* expression and creatinine levels (Figure S21). *PIBF1* encodes the protein PIBF1/CEP90, a component of the basal body essential for ciliogenesis and has been implicated as a cause of Joubert syndrome with kidney disease.^48–50^ Altogether there appear to be a number of lines of evidence supporting this novel association.

#### WDPCP

A number of variants within the *WDPCP* gene were found to be significantly associated with GGT (most significant p-value 1.79e-10 for variant rs7566031), total cholesterol (most significant p-value 1.21e-08 for variant rs7566031), and LDL cholesterol (most significant p-value 1.79e-12 for variant rs7566031) levels (Figures 2,4). None of these associations have been previously reported for any variant within 100kb of the *WDPCP* locus, and the association with LDL was specifically found to replicate, at both the variant and gene level, in our meta-analysis. WDPCP was only found to be significantly expressed in adipose tissue, of all the tissues investigated (Figure 6), and although nearly every variant significantly associated with LDL levels was also significantly associated with *WDPCP* expression levels (Figure S33), the nature of the association peak would suggest that the peak within the *WDPCP* locus may represent the tail of an association peak centered over the neighboring gene, *EHBP1*, which has been previously reported as associated with LDL and total cholesterol levels.^51,52^ That being said, the *EHBP1* gene has not been implicated in any Mendelian genetic disorders, whereas bi-allelic pathogenic variants in WDPCP have been shown to cause Bardet-Biedl syndrome, a disorder characterized, in part, by hepatic fibrosis and dyslipidemia, both phenotypes characterized by perturbed GGT and LDL cholesterol levels.^8,35^

### Replication of known associations

Of the 27 significant gene-trait associations we identified that replicate previously-known associations from the literature, 11 have been reported as mapping to the ciliary candidate genes that we set out to study: *CENPF* with Creatinine,^53,54^ *CEP164* with HDL,^44,55^ *POC5* with Cholesterol LDL and HDL,^51,52,56,57^ *PROSER3* with HDL,^52^ *RP1* with Cholesterol and LDL,^51,58,59^ *RP1L1* with Triglycerides,^51,52^ *SDCCAG8* with Creatinine,^53,55,60^ and *SPATA7* with Creatinine^54^ (Figures 2-3, S5-6, S22-23, S25-28). Our study adds further evidence to the hypothesis that these ciliary genes play in important role in affecting these laboratory phenotypes in the general population.

The remaining 16 significant, previously-reported associations mapped to 5 ciliary genes in our study (*CEP170*, *DYNC2LI1*, *IFT172*, *TRIM32*, *TTC8*), all of which exist within gene-dense regions of the genome. In these cases, the previously-reported mapped gene for the associated locus was different than the ciliary gene being investigated. In all cases, we found that the peak of association spanned multiple genes, making the identification of a single causative gene difficult. Four of these genes are discussed in more detail below.

#### *CEP170* and *TTC8*

Interestingly, in the case of *CEP170* and *TTC8*, the previously-mapped gene for each association peak was a different ciliary gene, and in both cases were ciliary genes that we also identified as significantly associated with the same trait. We found variants within the *CEP170* gene to be significantly associated with creatinine (Figure 3), with the association at this locus previously reported as mapping to the neighboring *SDCCAG8* gene (also found in our study); we found variants within the *TTC8* gene to also be significantly associated with creatinine (Figure 3), with the association at this locus previously reported as mapping to the neighboring *SPATA7* gene (also found in our study). All four of these ciliary genes are significantly expressed in the kidney (Figure 6). The association peak at the *SPATA7*/*TTC8* locus spans five genes altogether, with our eQTL analysis more supportive of *SPATA7*, rather than TTC8, as the potentially causative gene at this locus (Figures S28, S32). The association peak at the *SDCCAG8*/*CEP170* locus, on the other hand, spans only these two ciliary genes, with our eQTL analysis showing strong evidence for correlation between expression levels of both of these genes and creatinine levels (Figures S7, S27). In fact, our eQTL analysis shows that each variant associated with changes in expression of *SDCCAG8* has the same direction of effect on *CEP170* expression.

#### *DYNC2LI1* and *IFT172*

Two additional ciliary genes with significant, previously-reported associations bear mention: *DYNC2LI1* and *IFT172*. Both of these genes, critical components of the ciliary IFT machinery,^61,62^ are located in gene-dense regions of the genome that have previously been associated with a number of patient phenotypes. The *DYNC2LI1* gene neighbors the genes *ABCG5* and *ABCG8*, both known to be important in cholesterol metabolism,^51,52,59,63^ while the *IFT172* gene abuts the *GCKR* gene, which has been extensively associated with blood glucose, triglyceride, LDL, GGT, AlkPhos, and creatinine levels.^44,51–54,64–69^

We found the *DYNC2LI1* locus to be significantly associated with LDL and total cholesterol, as has been previously published, but also found significant associations with A1c (most significant p-value 4.50e-13 for rs116520905), Creatinine (most significant p-value 1.14e-08 for rs116520905), and ALT levels (most significant p-value 4.29e-14 for rs56266464) – all of them previously unreported, and none of which replicated in our meta-analysis (Figures 2-5). The *DYNC2LI1* gene is known to be causative of two Mendelian ciliopathy syndromes, Ellis-van Creveld syndrome and short rib polydactyly syndrome, both syndromes sometimes characterized by kidney and liver disease.^62,70^ eQTL analysis was not particularly supportive of the *DYNC2LI1* gene being the causative gene in the locus for any of these associations (Figure S10), and altogether it seems that these associations may be driven by the significant variants’ effects on neighboring genes such as *ABCG5* and *ABCG8.*

The *IFT172* locus, on the other hand, was found to be significantly associated with nine separate phenotypes (A1c, AlkPhos, Cholesterol, Creatinine, GGT, Glucose, LDL, Triglycerides, Urea)(Figures 2-5), all of which had been previously reported and mapped to the neighboring *GCKR* gene. *IFT172* is a critically important gene known to be causative of at least five severe ciliopathy syndromes characterized by renal disease, obesity, impaired glucose handling, dyslipidemia, and hepatic fibrosis.^61,71,72^ We found *IFT172* to be ubiquitously expressed in all tissues we examined (Figure 6), and our eQTL analysis revealed significant correlations between p_trait_ and p_eQTL_ for *IFT172* and a number of hepatic, liver, and lipid phenotypes (Figure S13), providing further evidence linking *IFT172* to the associated phenotypes. Thus altogether it remains difficult to confidently assign causality at this locus to either *GCKR* or *IFT172* alone.

### DiCE/pathway analysis reveals divergent phenotype associations for different ciliary compartments

To integrate the multiple lines of evidence we had gathered linking each ciliary gene to each significantly associated phenotype, we employed a Diverse Convergent Evidence^73^ (DiCE) analysis to estimate the strength of available corroborating data supporting a given gene-trait association. Each gene-trait pair with a significant association in our discovery analysis was given a DiCE score based on the reproducibility of the association, tissue expression analysis, eQTL analysis, and associated Mendelian disease phenotypes. The smallest score received was a 3 (given to *KIF17*-AlkPhos and *NIN*-Creatinine), whereas four genes received the maximum possible score of 11 (*IFT172*-Glucose, Triglycerides; *IQCB1*-Creatinine; *SDCCAG8*-Creatinine; *TRIM32*-Creatinine). These DiCE scores were used to generate a schematic (Figure 7) illustrating the strength of evidence supporting each ciliary gene’s association with a given phenotype domain.

**Figure 7.**
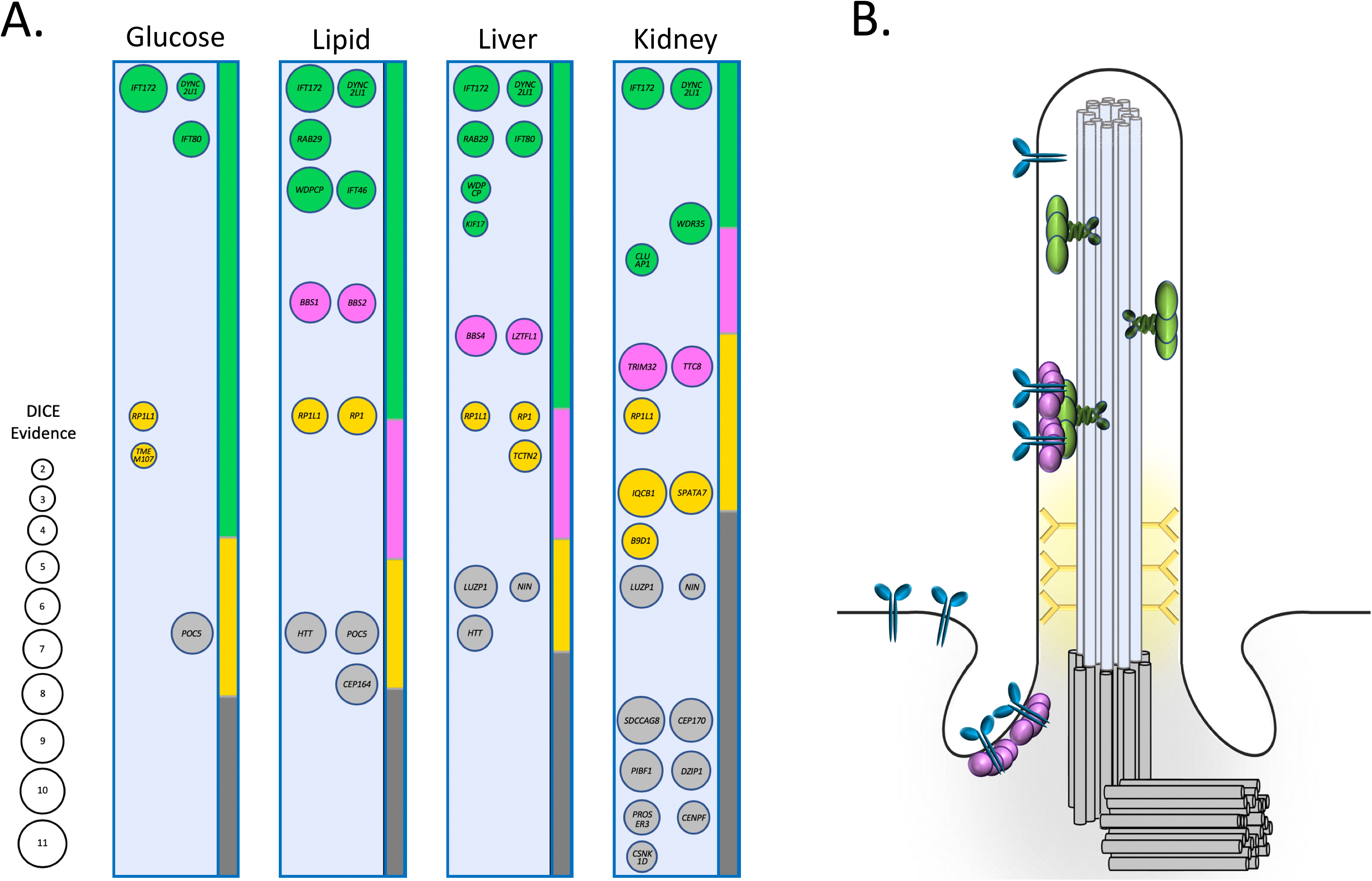
Schematics illustrating the results of DiCE/Pathway analysis, and a depiction of the primary cilium. (A) To integrate the multiple lines of evidence supporting each ciliary gene’s association with a given phenotype domain, we employed a Diverse Convergent Evidence (DiCE) analysis approach estimating the strength of available corroborating data (detailed in Table S14). The DiCE scores for each gene, illustrating the strength of evidence supporting each ciliary gene’s association with a given phenotype domain, are displayed here, with the magnitude of the score indicated by the size of each gene bubble. For each phenotype domain, gene bubbles are displayed at the same location for ease of comparison. Genes are colored by ciliary subcompartment – green for IFT-related genes, pink for the BBSome, orange for the transition zone, and grey for the basal body. A stacked bar graph is also displayed for each phenotype domain, illustrating the summed DiCE scores for all genes, by ciliary subcomparmtent, as a proportion of the total DiCE score per phenotype domain. (B) A schematic of the primary cilium. The basal body and centrioles are displayed in grey, the ciliary transition zone in orange, the IFT machinery in green, and the BBSome in pink. The core of 9 microtubule doublets and the overlying membrane of the organelle are shown, and receptors being trafficked into/along the cilium are shown in blue.

Using this approach, interesting patterns emerged in the data implicating different ciliary compartments in different disease processes. The DiCE score for genes affecting each phenotype domain were summed, by ciliary compartment, to generate a 4×4 contingency table (Table S16). A chi-square test performed on this data revealed significant difference across groups (p = 2.728e-05), primarily driven by the relative overabundance of basal body-associated genes and underabundance of IFT-associated genes within the kidney phenotype group (Figure S41). Of the 12 genes important to ciliary basal body function, 9 (75%) were found to associate with kidney phenotypes, whereas only 3 (25%) were found to associate with either liver or lipid traits, and only 1 (8%) was found to associate with glucose traits (Figure 7). At the same time, IFT-related genes accounted for 50% of the genes significantly associated with glucose traits and 42% of genes significantly associated with liver and lipid traits, but for only 21% of genes associated with kidney traits (Figure 7). Taken together, these data indicate that components of the ciliary basal body appear to play an important role in the normal function of the kidney, while ciliary genes involved in IFT appear more critical to liver-, lipid-, and glucose-related traits.

Yet another pattern that emerged in the data regarded differences in pleiotropy across different phenotypes. A majority of genes associated with either glucose (83%), lipid (67%), or liver-related (57%) traits demonstrated pleiotropic effects, being significantly associated with multiple traits across these three domains, whereas only 5 of 19 (26%) genes associated with kidney function demonstrated any pleiotropy at all (Figure 7). Interestingly, all five of the genes significantly associated with kidney phenotypes that were found to exhibit pleiotropy were specifically also associated with liver traits, with three (*DYNC2LI1*, *IFT172*, and *RP1L1*) exhibiting pleiotropic affects across all four phenotypic domains.

Taken together, the data yielded by our DiCE/Pathway analysis, seem to indicate that there exist divergent ciliary genetic pathways, one more specific to kidney function, and the other more specific to liver, lipid, and glucose physiology.

## DISCUSSION

Our study set out to investigate associations between common variants in genes important to primary cilium structure/function and 12 diverse quantitative laboratory traits associated with common complex diseases. Our analysis identified 73 statistically-significant gene-trait associations across 34 of the 122 ciliary genes that we examined. With few exceptions, these ciliary genes were found to be widely expressed in human tissues relevant to the phenotypes being studied. In the vast majority of cases, significant trait-associated variants were also found to act as eQTLs for the ciliary genes being investigated, with strong evidence for correlation between gene expression levels and patient phenotypes. Strikingly, in many cases, the phenotypes significantly associated with the ciliary gene being studied were features of the Mendelian ciliopathies caused by rare loss of function variants in the same gene. Together, our data provide a number of lines of evidence supporting the cilium’s role in the pathogenesis of common disease, and challenge the widely-held belief that the cilium is an organelle important mainly in development and in rare syndromic disease pathogenesis.

The finding that the primary cilium is involved in common disease pathogenesis is an important milestone in our understanding of this organelle’s role in human physiology. Just as the cilium had been overlooked for decades before being rediscovered as a major player in rare Mendelian disease, the discovery of the cilium’s role in common complex disease pathogenesis may represent a new rethinking of the importance of this tiny organelle. Perhaps even more exciting, however, is the possibility that common pathologic pathways might underlie the development of both rare syndromic and common complex disease. This possibility is further highlighted by perhaps the most interesting and unexpected finding of our study – that different ciliary subcompartments are specifically associated with distinct sets of patient phenotypes. This concept, that even within a single organelle, distinct subsets of genes/proteins may act together to affect patient phenotypes in divergent ways, is not new to Mendelian disease, but is an interesting and fascinating finding in common disease genetics. In fact, the recognition that particular ciliary pathways/complexes may be specifically involved in divergent common disease processes leaves the door open for novel targeted therapeutics and diagnostic approaches for common disease, informed by our extensive knowledge of ciliopathy pathogenesis and rare disease pathways. The further characterization of these previously unknown links between the cilium and common complex disease will establish a new field of study that has the potential to fundamentally transform our understanding of common disease pathogenesis.

An unexpected result of our analysis was the identification of four ciliary genes that were not significantly expressed in any tissues we examined in the GTEx database – *CENPF*, *WDPCP*, *RP1*, and *RP1L1* (Figure 5, Table S15). In the case of *CENPF* and *WDPCP*, this is a surprising finding, as rare variants in each of these genes are known to cause complex, multisystem ciliopathy disorders,^74–76^ suggesting broad expression across multiple tissue types. We cannot rule out the possibility that these genes are expressed broadly only during fetal development but not adulthood, that they are expressed only in a small but critical subset of cells, or that the corresponding mRNAs are unstable, making their detection in the GTEx database unlikely.

*RP1* and *RP1L1*, on the other hand, represent more interesting cases. These two genes are highly homologous to each other, with mutations in each being well-established causes of Mendelian disorders characterized by retinal degeneration.^77,78^ In both cases, the genes are thought to be expressed exclusively in the retina.^78^ In our discovery analysis, both gene loci were found to be significantly associated with numerous laboratory phenotypes spanning multiple organ systems (*RP1*: AST, GGT, LDL, Cholesterol; *RP1L1*: A1c, AlkPhos, ALT, AST, Cholesterol, Creatinine, and GGT), which is not consistent with retina-only expression of either gene. In both cases, the variants significantly associated with each phenotype were also found to significantly affect *RP1*/*RP1L1* expression levels based on our eQTL analysis, with evidence for correlation between gene expression levels and laboratory trait levels (Figures S25-26). *RP1* exists in a locus with only one nearby gene, *SOX17*; interestingly, there did not appear to be a significant correlation between variants associated with *SOX17* expression and any laboratory trait level (Figure S36), suggesting that the association between variants in this locus and AST, GGT, and LDL may, in fact, be mediated by changes in *RP1* expression, through currently unclear mechanisms. *RP1L1* exists in a locus with a number of neighboring genes – *C8orf74*, *PINX1*, *PRSS55*, *SOX7*– and although our eQTL analysis found that the variants significantly associated with each trait also associated with the expression of many of these genes, by far the strongest correlation was with *RP1L1* expression (figure S26, S37-40). Clearly, further work is needed to understand if and how genetic variants in these two apparently retina-specific genes are affecting non-retinal phenotypes.

Another interesting consequence of our analysis was the identification of gene-dense genomic loci, containing at least one ciliary gene, that were found to strongly associate with laboratory traits. It has traditionally been difficult to attribute causality to a single gene in loci such as this, which display high degrees of linkage disequilibrium across multiple genes. In two cases, the loci we identified contained two different ciliary genes (*CEP170* and *SDCCAG8*; *TTC8* and *SPATA7*) (Figures S7, S27, S28, S32). These examples suggest that a single genomic locus might be associated with a given phenotype through the perturbation of expression of multiple genes operating within the same pathway. In fact, the primary cilium may serve as an excellent model for observing such associations and testing this hypothesis.

In a different case, we identified variants within the critically important ciliary IFT gene *IFT172* as being significantly associated with nine separate phenotypes – A1c, AlkPhos, Cholesterol, Creatinine, GGT, Glucose, LDL, Triglycerides, and Urea (Figure S13). *IFT172* neighbors the gene *GCKR*, which has traditionally been considered the mostly likely causal candidate gene in this locus. *GCKR* encodes a well-studied regulatory protein important in controlling glucose flux through the glycolytic pathway, and has been shown *in vitro* to have significant effects on triglyceride and glucose metabolism.^79^ Previous fine mapping work of the *GCKR* locus has identified a relatively common missense variant in the gene as the putative causal variant responsible for the strong association of this locus with these multiple phenotypes.^80^ However, there is minimal evidence that the protein encoded by the *GCKR* gene actually plays an important role in disease pathogenesis; knockout mice completely deficient in *GCKR* are normoglycemic except under extreme dietary conditions, and express no other apparent health phenotypes,^81,82^ and domestic cats have been shown to be completely deficient in *GCKR* mRNA and protein product as a species.^83^ It seems unlikely that a gene that, in other species, can be completely abolished without obvious health effects could be solely responsible for so many robust human phenotype associations. *IFT172*, on the other hand, is a critically important gene known to be causative of at least five severe ciliopathy syndromes characterized by renal disease, obesity, impaired glucose handling, dyslipidemia, and hepatic fibrosis,^61,71,72^ and seems a much more likely candidate gene for the association peak based solely on it’s known pathogenic potential. Furthermore, unlike *GCKR,* which is expressed primarily in the liver,^84^ *IFT172* was found to be ubiquitously expressed in all tissues we examined, with our eQTL analysis revealing significant correlations between p_trait_ and p_eQTL_ for *IFT172* and a number of hepatic, liver, and lipid phenotypes. This situation is reminiscent of the ongoing discussion surrounding the *FTO* locus – countless GWAS and fine mapping studies have linked the locus surrounding the *FTO* gene to body mass index and body fat percentage, but recent evidence suggests that the neighboring ciliary gene *RPGRIP1L*, may at least be partially driving these associations.^85–87^ Clearly more work is needed to disentangle the contributions of both *GCKR* and *IFT172* to the many patient phenotypes associated with this genomic locus.

Altogether, our data demonstrate the utility of a Mendelian disease-based approach to common variant association studies, and show that Mendelian disease genes can play an important role in common disease pathogenesis. Our data implicate the primary cilium as an important player in common disease, challenging the widely-held belief that the cilium is an organelle important mainly in development and in rare syndromic disease pathogenesis. It is not altogether surprising that such a critically important organelle, causative of rare ciliopathies characterized by phenotypes ranging from kidney failure to hepatic fibrosis, has important roles in the function of these same organ systems more generally; it stands to reason that a pathway important in rare disease might also be important for common disease affecting the same organ systems. The continued application of techniques similar to those described here to other phenotypes/Mendelian diseases is likely to yield many additional fascinating associations that will begin to integrate the fields of common and rare disease genetics, and provide insight into the pathophysiology of human diseases of immense public health burden.

## ACKNOWLEDGMENTS

We would like to thank Michael P. Hart and Pinar S. Gurel for their careful reading of the manuscript and helpful comments.

## FUNDING

TGD is supported in part by the NIH T32 training grant 5T32GM008638-23. MDR is supported in part by NIH R01 AI077505.

## METHODS

### Selection of cilium genes for analysis

122 genes with well-characterized roles in ciliary biology^17^ were selected for analysis. In all cases, gene transcript coordinates were defined using reported transcriptional start and stop sites that yielded the maximum transcript length for each gene. A complete list of the genes included in this study, along with the genomic coordinates used, is included in Table S17. All genes were classified as being components of ciliary sub-compartments (basal body, transition zone, BBsome, or IFT-associated) based on literature review^17,88^. Mendelian disease associations for each gene were obtained from the Online Mendelian Inheritance in Man (OMIM)^89^ database and literature review.

### Preparation of UKBB Genotype data

The UKBB cohort release version 2 has deep genetic and phenotypic data on ~500,000 subjects across the United Kingdom. Subjects were genotyped on two similar genotype arrays across 106 batches and imputed to 96 million variants.^90^ Quality control of genotype data was performed largely as previously described.^90^ Subjects with poor quality genotype data were excluded from our analysis. Genetically related subjects (second-degree or closer with pi-hat larger than 0.25) were also excluded. European ancestry subjects were extracted using a combination of self-reported European ancestry (UKBB Data-Field: 21000) and genetic principal component analysis (UKBB Data-Field: 22006). We next filtered variants to retain only those with a minor allele frequency between 0.4 and 0.01, and which fell within the genomic regions defined by our set of 122 ciliary genes. This final set of variants was subjected to linkage disequilibrium (LD) pruning with an r^2^ threshold of 0.5 using a 50 variant window shifted by 5 variant steps. After quality control 378,213 subjects and 16,874 genetic variants were included in our discovery association analyses. Principle components of genomic data that were subsequently used as covariates for association studies were provided in the UKBB data release.

### Preparation of UKBB Phenotype data

For all 378,213 subjects passing genotype quality control, laboratory measurement information from the UKBB cohort release version 2 was extracted for the following phenotypes: Alanine aminotransferase (ALT; UKBB Data-Field: 30620), Alkaline Phosphatase (AlkPhos; UKBB Data-Field: 30610), Aspartate aminotransferase (AST; UKBB Data-Field: 30650), Cholesterol (UKBB Data-Field: 30690), Creatinine (UKBB Data-Field: 30700), Gamma glutamyltransferase (GGT)(UKBB Data-Field: 30730), Glucose (UKBB Data-Field: 30740), Glycated haemoglobin (A1c; UKBB Data-Field: 30750), HDL cholesterol (HDL; UKBB Data-Field: 30760), LDL cholesterol (LDL; UKBB Data-Field: 30780), Triglycerides (UKBB Data-Field: 30870), and Urea (UKBB Data-Field: 30670). For each subject, only baseline initial encounter laboratory measurements were included. For each phenotype, outlier subjects with laboratory measurements greater than three standard deviations from the mean were excluded. For each phenotype, values were subjected to BoxCox^91^ or natural log transformation to maximize the normalcy of distribution. Following these quality control steps, the following number of subjects, per phenotype, were included in association analyses: A1c (n= 354,174), AlkPhos (n= 357,629), ALT (n= 354,959), AST (n= 355,215), Cholesterol (n= 359,024), Creatinine (n= 358,650), GGT (n= 354,772), Glucose (n= 324,597), HDL (n= 327,335), LDL (n= 358,417), Triglycerides (n= 354,118), Urea (n= 357,364). Summary statistics for the laboratory data used for our analysis can be found in Table S1. The UKBB data was accessed using application #32133.

### Discovery Association Studies

Association studies were performed using the genotype and phenotype data prepared as described above using the PLatform for the Analysis, Translation, and Organization of large-scale data (PLATO)^92,93^, a standalone program developed by the Ritchie lab for the performance of phenome-wide linear/logistic regression of genetic variants against participant phenotypes. Linear regression, assuming an additive genetic model, was performed across all 16,874 ciliary genetic variants and the 12 phenotypes listed above. Regression models were adjusted by age, sex, and the first ten principle components of the corresponding genomic data for all 378,213 European ancestry subjects.

### Meta-analyses

Transethnic meta-analyses were carried out using the Stouffer method in the program METAL^94^ using publicly-available GWAS^44,95–97^ and meta-analysis^53,56,58,98–100^ summary statistics. Studies were analyzed together only if performed on the same trait, with the exception of Glucose (where studies looking at both fasting and non-fasting glucose levels were analyzed together) and Creatinine (where studies looking at both serum creatine and estimated glomerular filtration rate (eGFR, a metric derived from the serum creatinine, along with age, sex, and race) were analyzed together). In all cases, variants were filtered to retain only those which fell within the genomic regions defined by our set of 122 ciliary genes. Where necessary, variant nomenclature was standardized to ensure that identical variants were appropriately analyzed across studies. The studies included for meta-analysis contained no overlapping sample sets, with the exception of the meta-analyses of Creatinine/eGFR, and Urea. In both these cases, there was an overlap of 6,492 individuals between the large meta-analysis of eGFR and Urea traits (n = 567,460) conducted by Wuttke et al.^53^ (which contained samples from an earlier release of the BioVu dataset), and a GWAS performed on the current BioVU dataset^97^ (urea n = 33,322, creatinine n = 33,493). In both cases, we performed separate meta-analyses both including and excluding the current BioVu dataset (Figure S42) and found no differences in terms of the number of significant replicating loci identified. The meta-analyses including both of these data sets, with the small number of overlapping samples, are displayed in the main figures of the manuscript. The total transethnic sample size for each trait meta-analyzed was: A1c (n = 215,760), AlkPhos (n = 178,929), ALT (n = 208,562), AST (n = 208,165), Cholesterol (n = 369,671), Creatinine/eGFR (n = 831,567), GGT (n = 136,161), Glucose fasting/non-fasting (n = 229,080), HDL (n = 311,121), LDL (n = 312,410), Triglycerides (n = 346,901), and Urea (n = 798,670). A more detailed description of the studies included in our meta-analyses can be found in Table S18.

### Significance Thresholds and Novel Findings

For all 12 traits in our discovery analysis, analyzed across 16,874 ciliary variants, the Bonferroni-corrected p-value significance threshold was calculated at 2.5e-7, but we chose to only report as significant those associations reaching the more stringent, commonly-accepted genome-wide p-value significance threshold of 5e-8. Our meta-analyses included many more variants than those in our discovery analysis, and thus we defined significant replication in our meta-analyses in two ways; at the variant level, and at the gene level. At the variant level, analyzing only the union of variants present in both the discovery analysis and meta-analysis for a given trait, a variant-trait association was deemed to replicate if: (1) the variant was significantly associated with the trait in our discovery analysis with p<5e-8 and (2) the p-value of association for the same variant, with the same trait, in the meta-analysis was below the Bonferroni-corrected significance threshold for the number of tests performed, defined as the number of variants shared between the discovery analysis and replication meta-analysis for a given trait. Listed here, for each trait, is the total number of variants shared between the discovery analysis and replication meta-analysis, along with the corresponding Bonferroni-adjusted significance threshold used to determine significant variant-level replication: A1c (14,271; 3.5e-6), AlkPhos (15,197; 3.3e-6), ALT (15,197; 3.3e-6), AST (15,186; 3.3e-6), Cholesterol (15,143; 3.3e-6), Creatinine/eGFR (13,331; 3.8e-6), GGT (15,171; 3.3e-6), Glucose (15,132; 3.3e-6), HDL (15,143; 3.3e-6), LDL (15,143; 3.3e-6), Triglycerides (15,143; 3.3e-6), Urea (13,336; 3.8e-6). The experiment-wise error rate for all tests across all phenotypes/variants in this analysis was calculated at 2.8e-7. At the gene level, a gene-trait association was deemed to replicate if: (1) any variant within the gene was found to be significantly associated with the trait in our discovery set with p<5e-8 and (2) the p-value of association between the same trait and any variant within the same gene in the meta-analysis was below the Bonferroni-corrected significance threshold for tests performed, defined using the total number of variants analyzed in the meta-analysis for a given trait. Listed here, for each trait, is the total number of variants in the replication meta-analysis, along with the corresponding Bonferroni-adjusted significance threshold used to determine significant gene-level replication: A1c (104,706; 4.8e-7), AlkPhos (153,637; 3.3e-7), ALT (153,186; 3.3e-7), AST (149,729; 3.3e-7), Cholesterol (92,118; 5.4e-7), Creatinine/eGFR (42,925; 1.2e-6), GGT (146,818; 3.4e-7), Glucose (91,397; 5.5e-7), HDL (92,106; 5.4e-7), LDL (92,012; 5.4e-7), Triglycerides (92,107; 5.4e-7), Urea (42,701; 1.2e-6). The experiment-wise error rate for all tests across all phenotypes/variants in this analysis was calculated at 4e-8. We defined novel findings as those that (1) were significant in our discovery analysis with p<5e-8, (2) replicated at the gene and/or variant level (as defined above) in meta-analysis, and (3) were at least 100kb away from any other variant previously reported as significantly associated with the given phenotype. The Bonferroni-corrected significance threshold for all 1,456,062 independent association tests performed in our entire study was calculated to be 3.4e-8.

### Discovery/Replication Meta-analysis Result Visualization

Results were visualized using a modified version of the Hudson R package^101^ to easily compare findings between our discovery analysis in the UKBB and replication meta-analyses of the same phenotype. For each Hudson plot, our discovery analysis results are plotted on the top half of the plot, with our meta-analysis results plotted in the bottom half of the plot. Each analyzed gene is given equal space, with all tested genetic variants for a given gene plotted at the midline of the gene block (i.e., variants are not represented in chromosomal space, but compressed to a single horizontal position per gene). For all Hudson plots, the study-wise Bonferroni-adjusted significance threshold (3.4e-8) is represented by a blue dashed line, while the Bonferroni-adjusted significance threshold for the trait being analyzed is represented by a red dashed line.

### Tissue Specific Expression Studies

Tissue-specific expression data for each gene with significant associations in our discovery analysis was obtained from version 8 of the Genotype-Tissue Expression (GTEx) Project.^29^ The GTEx Project was supported by the Common Fund of the Office of the Director of the National Institutes of Health, and by NCI, NHGRI, NHLBI, NIDA, NIMH, and NINDS. The data used for the analyses described in this manuscript were obtained from the GTEx Portal on 5/20/2020. The tissues analyzed were limited to those deemed to be relevant to the studied phenotypes, and included data from kidney, liver, pancreas, adipose, and skeletal muscle. A gene was considered to be significantly expressed in a given tissue if the GTEx-derived transcripts per million (TPM) for the gene in that tissue was greater than 1.5. A heatmap of tissue-specific expression information for each gene was generated using the ggplot2 package for R.^102^

### eQTL Plots

The eQTpLot package was used to make all three eQTL plots; the R package is available on GitHub at https://github.com/RitchieLab/eQTpLot.^31,32^

#### eQTL Plot

For each candidate gene with a significant association in our discovery analysis, we obtained a list of all variants associated with the gene’s expression level, across all tissues, from the GTEx version 8 dataset, as above. For each variant with significant associations with a given candidate gene’s expression in multiple tissues, we selected data only from the tissue with the most significant association. We defined a variant as an expression quantitative trait locus (eQTL) for the candidate gene if its p-value of association with gene expression (p_eQTL_) was <0.05. We next defined a locus of interest (LOI) for each candidate gene to include the candidate gene’s coordinates, along with 200kb of flanking genomic material on either side. We repeated our association studies, as described in “Discovery Association Studies” above, for each significant candidate gene-trait pair, this time analyzing all variants in the LOI. To create the eQTL Plot, we plotted each variant in the LOI in chromosomal space on the x-axis, with the p-value of association with the phenotypic trait (p_trait_) on the y-axis. If a variant was determined to be an eQTL for the candidate gene, it was plotted as a colored triangle – in blue, for variants that had congruent directions of effect (e.g. the variant was associated with increased transcript levels of the candidate gene, and also with an increase in the quantitative phenotype being studied), or in red for variants that had incongruous directions of effect (e.g. the variant was associated with increased transcript levels of the candidate gene, but with a decrease in the quantitative phenotype being studied), with a color gradient corresponding to the magnitude of p_eQTL_. The size of each triangle was set to correspond to the eQTL normalized effect size for the variant, as obtained from GTEx, while the directionality of each triangle was set to correspond to the direction of effect of the variant for the phenotypic trait. Variants that were not found to be eQTLs (no eQTL data available, or p_eQTL_ >0.05) for the candidate gene were plotted as grey boxes. A depiction of the genomic positions of all genes within the LOI was added below the plot using the package Gviz for R^103^.

#### eQTL Enrichment Plots

For variants within the LOI with p_trait_ < 5e-8, the proportion that were also candidate gene eQTLs was calculated and plotted, and the same was done for variants with p_trait_ > 5e-8. Significant enrichment for eQTLs among the variants with significant association with the phenotypic trait was determined by Fisher’s exact test. Bonferroni-corrected significance thresholds for enrichment were set at p < 7e-4 (adjusting for testing for eQTL enrichment across the 73 gene-trait pairs identified as significant in our discovery analysis). Variants with congruent and incongruent directions of effect, as described above, were considered separately.

#### P-P Plots

Each variant within a given LOI, as defined above, was plotted with p_eQTL_ along the x-axis, and p_trait_ along the y-axis. Correlation between the two probabilities was visualized by plotting a best-fit linear regression over the points, with the line equation displayed on the plot. The Pearson correlation coefficient and p-value of correlation were computed and displayed on the plot as well. Separate plots were made and superimposed over each other for variants with congruent and incongruent directions of effect, as described above. Bonferroni-corrected significance thresholds for Pearson correlation were set at p < 3.5e-4 (adjusting for testing for correlation across the 73 gene-trait pairs identified as significant in our discovery analysis, once for congruent eQTLs and once for incongruent eQTLs).

### DiCE Analysis

Each significant gene-trait pair was given a DiCE score^73^ ranging from 1 to 11 summarizing the amount of evidence linking the candidate gene with the associated trait. One point was assigned for the initial discovery association. One point was assigned if the association replicated in our meta-analysis (at either the variant or gene level), and two points were assigned if the association had been previously reported (for any variant within 100kb of the cilium candidate gene). One point was given if a gene was found to be expressed in at least one tissue relevant to the trait of interest, while two points were given if a gene was found to be expressed in all relevant tissue types (relevant tissue types considered were: Kidney (medulla) and Kidney (cortex) for Creatinine/Urea; Liver, Pancreas, and Muscle (skeletal) for Glucose/A1c; Liver, Adipose (subcutaneous) and Adipose (visceral) for Cholesterol/LDL/HDL/Triglycerides; Liver for AST/ALT/AlkPhos/GGT). If the association peak for a candidate gene was significantly enriched for candidate gene eQTLs, one point was assigned. If the Pearson correlation between p_trait_ and p_eQTL_ was greater than 0.25 with a p-value < 5e-5, one point was assigned. If both these outcomes were true, an additional one point was assigned. Lastly, if the associated trait was relevant to the Mendelian disease associated with the candidate gene, two additional points were assigned. As a single gene could be associated with multiple traits within a single trait domain (e.g. AlkPhos, ALT, AST, or GGT for Liver-related traits), the highest DiCE score within a trait domain was selected for each gene with multiple associations.

## Notes

### Competing Interest Statement

The authors have declared no competing interest.

