## Supplemental Figures for "Mendelian pathway analysis of laboratory traits reveals distinct roles for ciliary subcompartments in common disease pathogenesis"

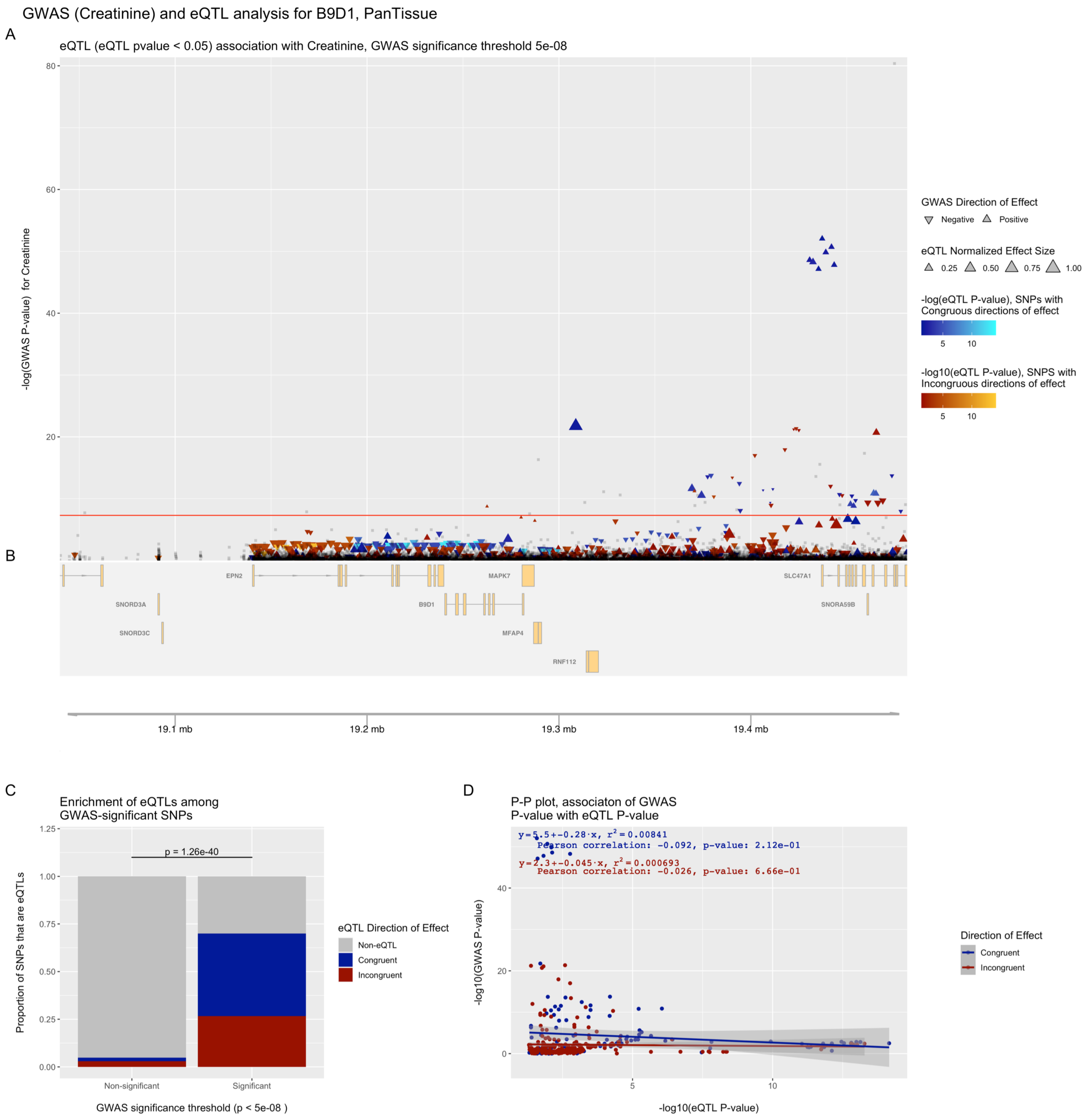


**Figure S1. eQTpLots analysis of the *B9D1* locus and all significant associated phenotypes**

For each phenotype: (A) Plot illustrating any potential colocalization between phenotype-significant variants and eQTLs for the given gene (B) A depiction of the genomic region surrounding the locus of interest (C) Enrichment of eQTLs for the given gene among trait-significant variants. The p-value of enrichment was determined by Fisher's exact test (D) P-P plot illustrating correlation between $p_{eQTL}$ and $p_{trait}$ for the gene of interest. Correlation between the two probabilities is visualized by plotting a best-fit linear regression over the points, with the line equation displayed on the plot. The Pearson correlation coefficient and p-value of correlation are displayed on the plot as well.

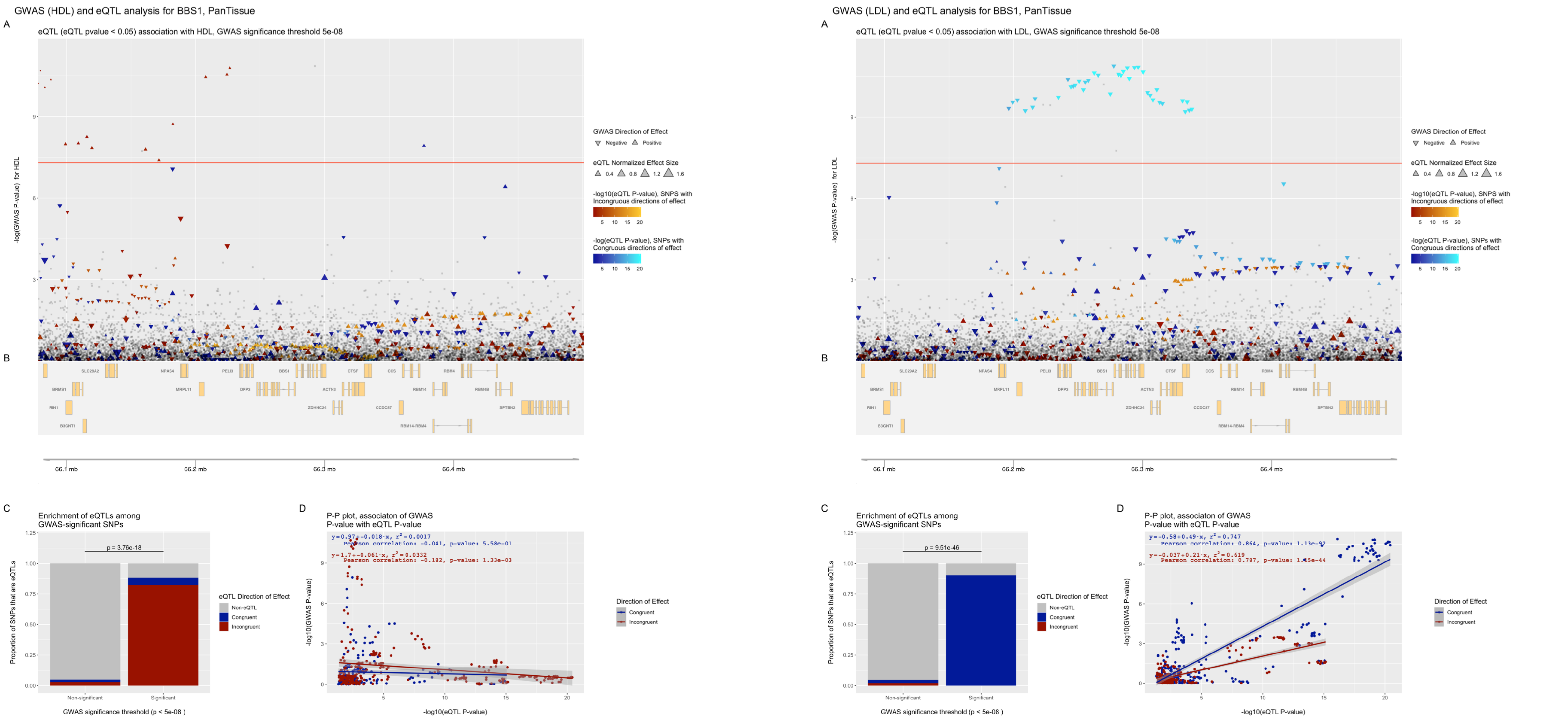


**Figure S2. eQTpLots analysis of the *BBS1* locus and all significant associated phenotypes**

For each phenotype: (A) Plot illustrating any potential colocalization between phenotype-significant variants and eQTLs for the given gene (B) A depiction of the genomic region surrounding the locus of interest (C) Enrichment of eQTLs for the given gene among trait-significant variants. The p-value of enrichment was determined by Fisher's exact test (D) P-P plot illustrating correlation between $p_{eQTL}$ and $p_{trait}$ for the gene of interest. Correlation between the two probabilities is visualized by plotting a best-fit linear regression over the points, with the line equation displayed on the plot. The Pearson correlation coefficient and p-value of correlation are displayed on the plot as well.

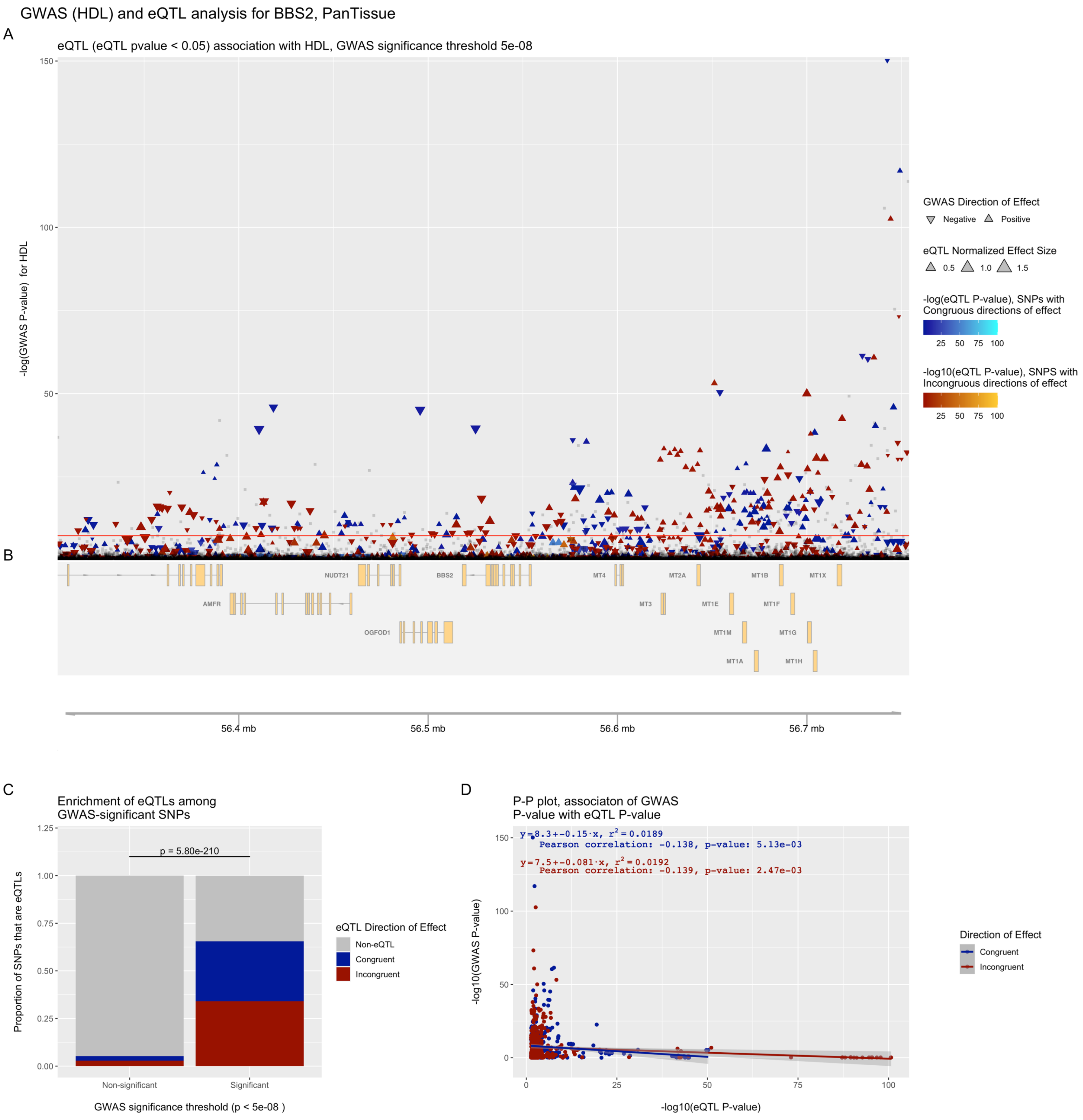


**Figure S3. eQTpLots analysis of the *BBS2* locus and all significant associated phenotypes**

For each phenotype: (A) Plot illustrating any potential colocalization between phenotype-significant variants and eQTLs for the given gene (B) A depiction of the genomic region surrounding the locus of interest (C) Enrichment of eQTLs for the given gene among trait-significant variants. The p-value of enrichment was determined by Fisher's exact test (D) P-P plot illustrating correlation between $p_{eQTL}$ and $p_{trait}$ for the gene of interest. Correlation between the two probabilities is visualized by plotting a best-fit linear regression over the points, with the line equation displayed on the plot. The Pearson correlation coefficient and p-value of correlation are displayed on the plot as well.

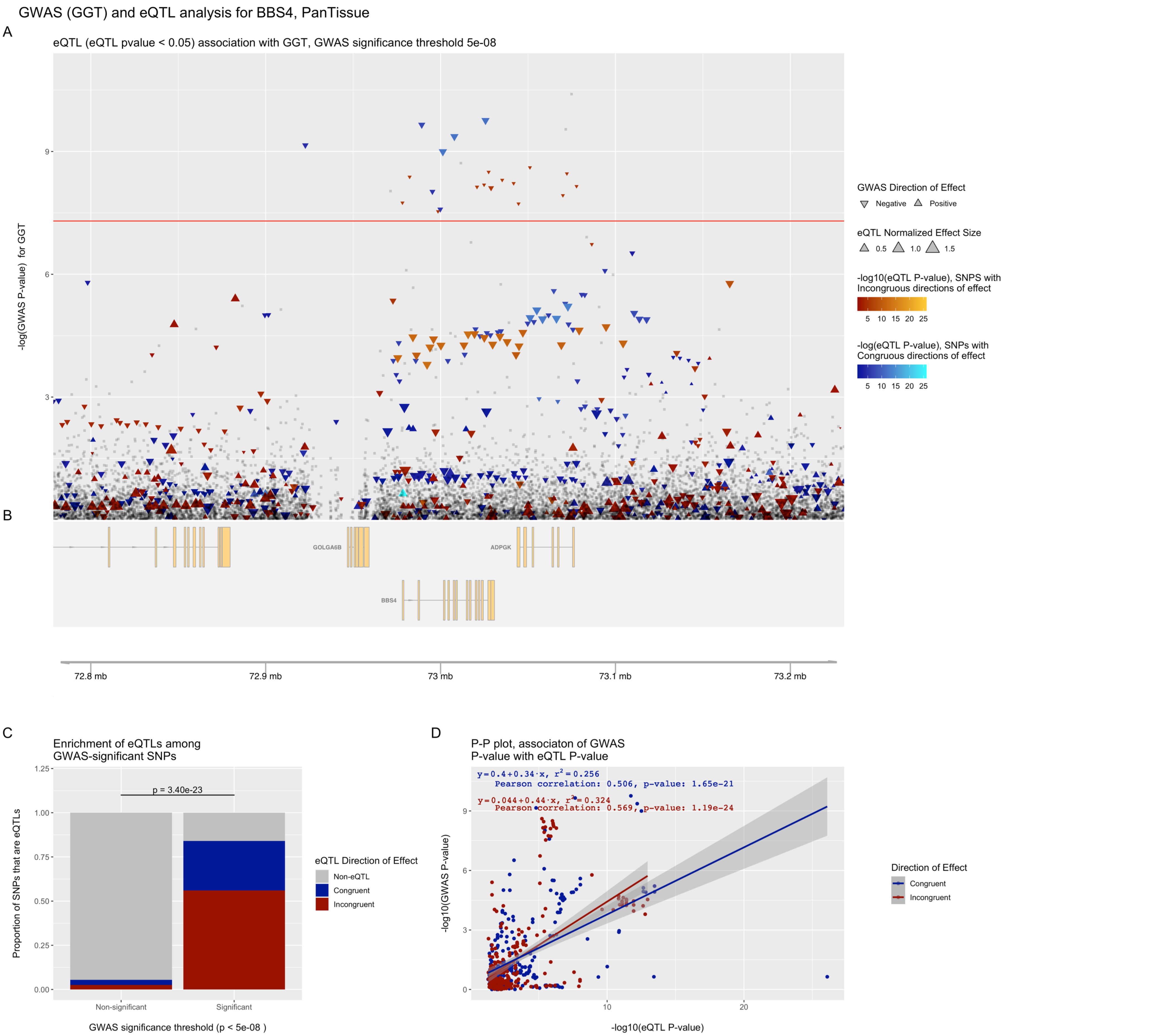


**Figure S4. eQTpLots analysis of the *BBS4* locus and all significant associated phenotypes**

For each phenotype: (A) Plot illustrating any potential colocalization between phenotype-significant variants and eQTLs for the given gene (B) A depiction of the genomic region surrounding the locus of interest (C) Enrichment of eQTLs for the given gene among trait-significant variants. The p-value of enrichment was determined by Fisher's exact test (D) P-P plot illustrating correlation between $p_{eQTL}$ and $p_{trait}$ for the gene of interest. Correlation between the two probabilities is visualized by plotting a best-fit linear regression over the points, with the line equation displayed on the plot. The Pearson correlation coefficient and p-value of correlation are displayed on the plot as well.

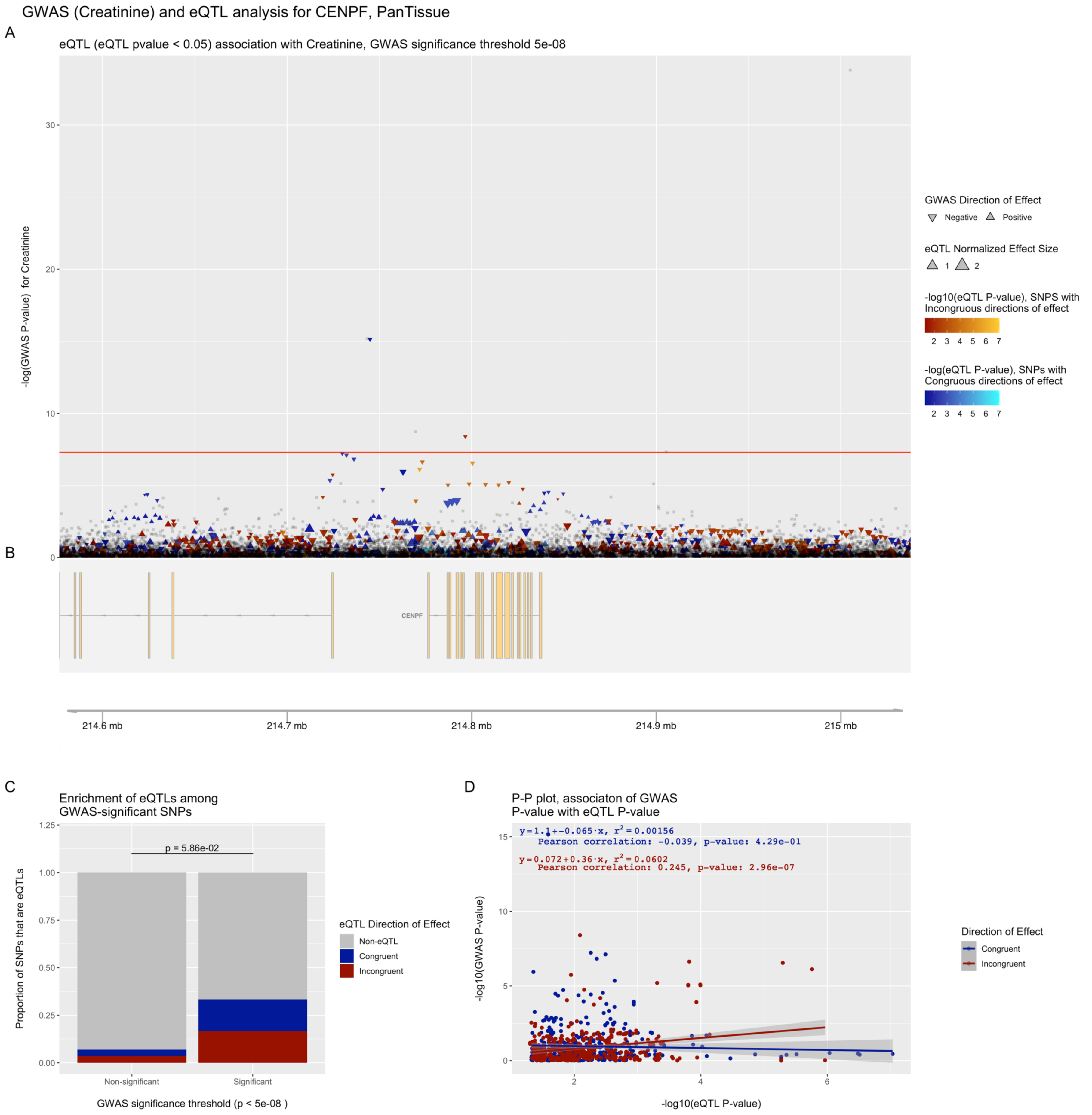


**Figure S5. eQTpLots analysis of the *CENPF* locus and all significant associated phenotypes**

For each phenotype: (A) Plot illustrating any potential colocalization between phenotype-significant variants and eQTLs for the given gene (B) A depiction of the genomic region surrounding the locus of interest (C) Enrichment of eQTLs for the given gene among trait-significant variants. The p-value of enrichment was determined by Fisher's exact test (D) P-P plot illustrating correlation between $p_{eQTL}$ and $p_{trait}$ for the gene of interest. Correlation between the two probabilities is visualized by plotting a best-fit linear regression over the points, with the line equation displayed on the plot. The Pearson correlation coefficient and p-value of correlation are displayed on the plot as well.

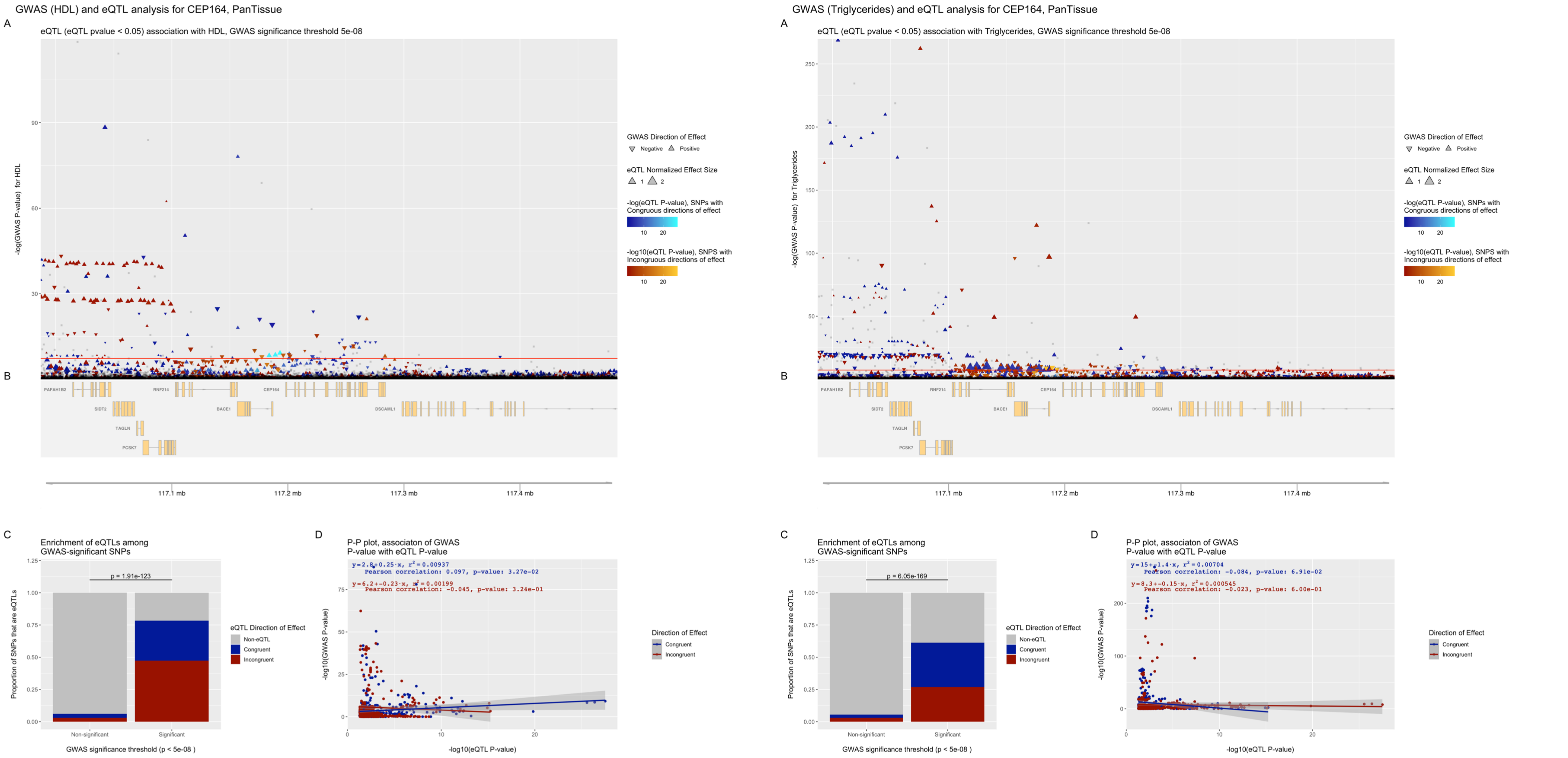


**Figure S6. eQTpLots analysis of the *CEP164* locus and all significant associated phenotypes**

For each phenotype: (A) Plot illustrating any potential colocalization between phenotype-significant variants and eQTLs for the given gene (B) A depiction of the genomic region surrounding the locus of interest (C) Enrichment of eQTLs for the given gene among trait-significant variants. The p-value of enrichment was determined by Fisher's exact test (D) P-P plot illustrating correlation between $p_{eQTL}$ and $p_{trait}$ for the gene of interest. Correlation between the two probabilities is visualized by plotting a best-fit linear regression over the points, with the line equation displayed on the plot. The Pearson correlation coefficient and p-value of correlation are displayed on the plot as well.

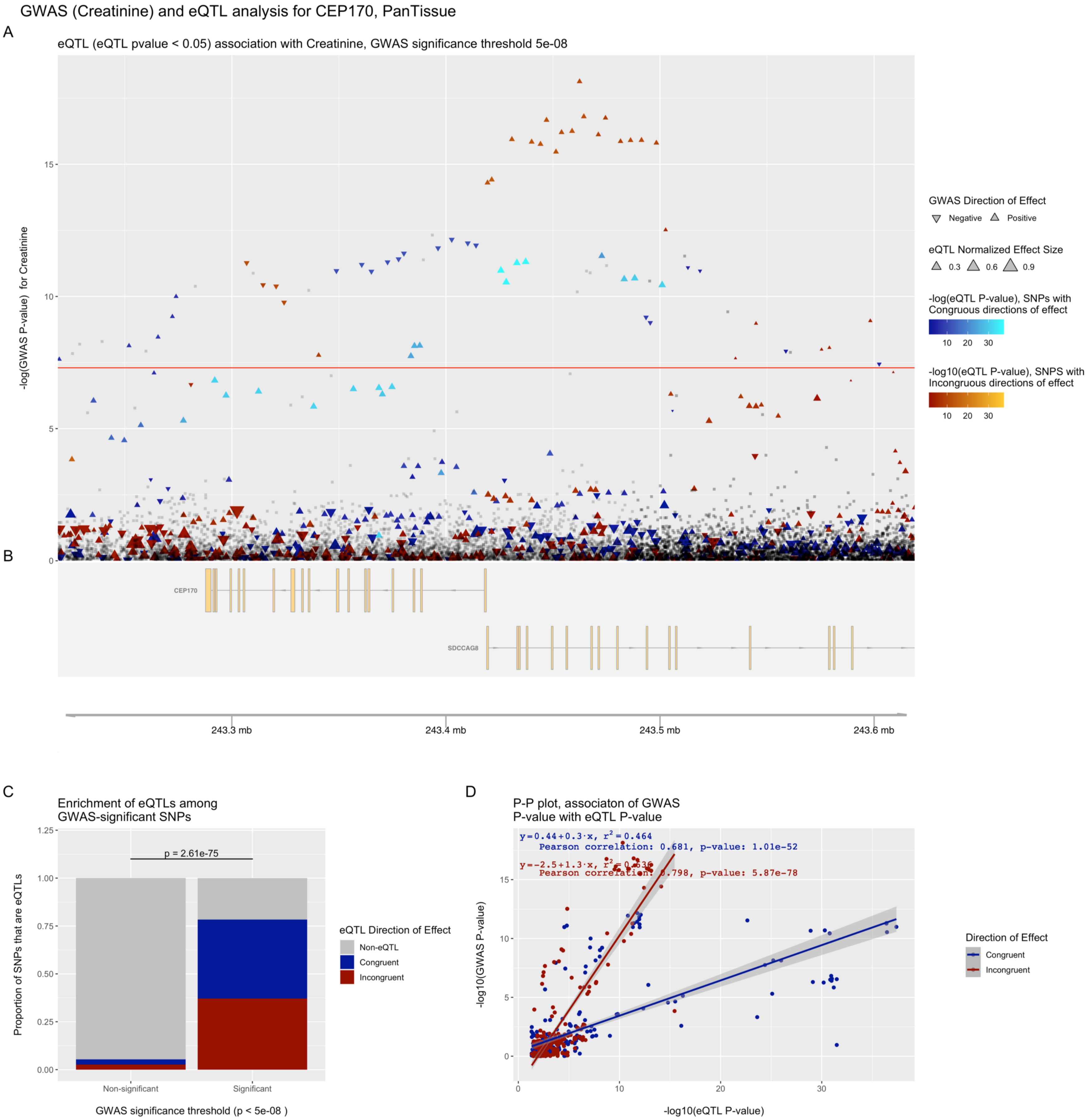


**Figure S7. eQTpLots analysis of the *CEP170* locus and all significant associated phenotypes**

For each phenotype: (A) Plot illustrating any potential colocalization between phenotype-significant variants and eQTLs for the given gene (B) A depiction of the genomic region surrounding the locus of interest (C) Enrichment of eQTLs for the given gene among trait-significant variants. The p-value of enrichment was determined by Fisher's exact test (D) P-P plot illustrating correlation between $p_{eQTL}$ and $p_{trait}$ for the gene of interest. Correlation between the two probabilities is visualized by plotting a best-fit linear regression over the points, with the line equation displayed on the plot. The Pearson correlation coefficient and p-value of correlation are displayed on the plot as well.

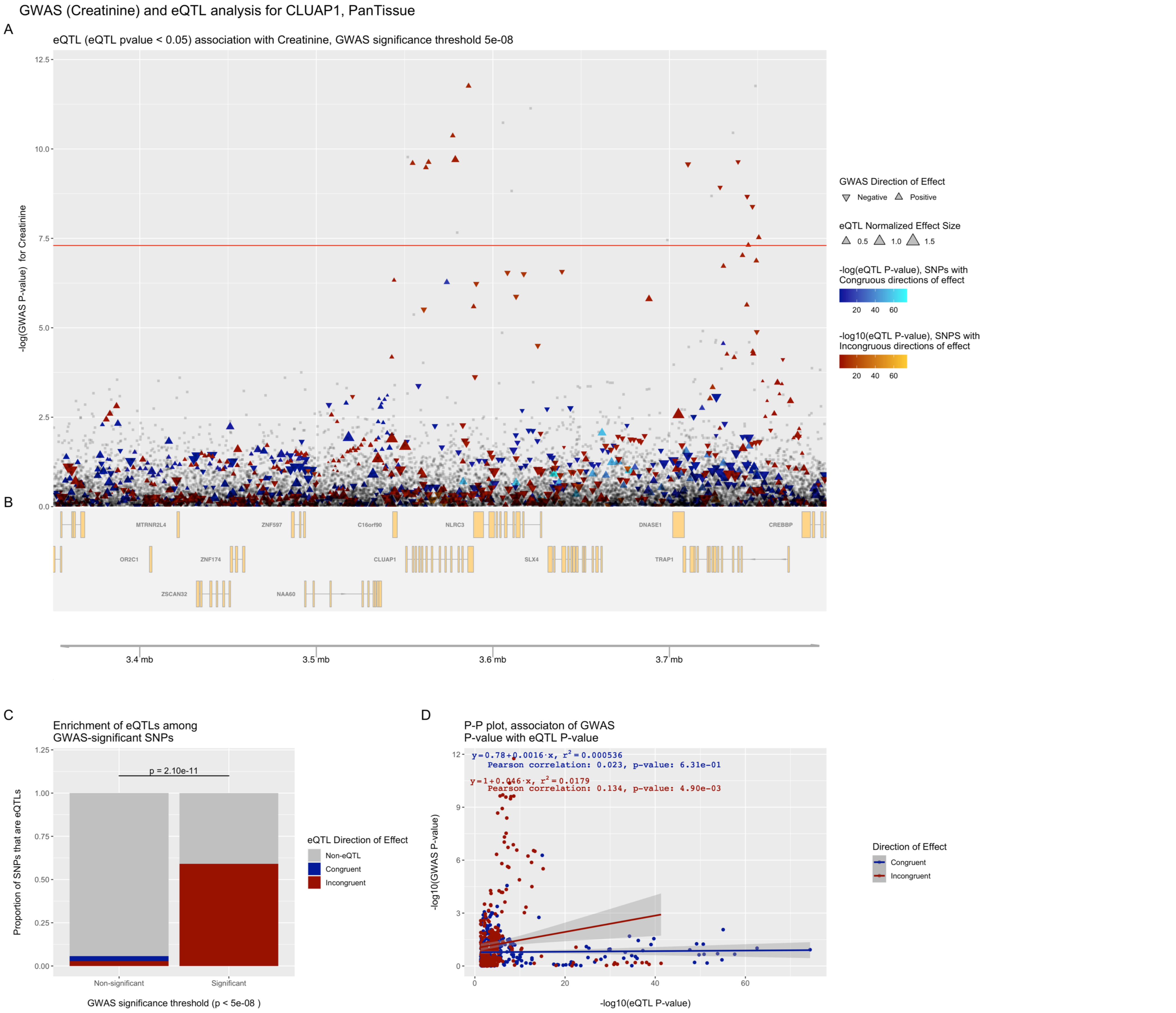


**Figure S8. eQTpLots analysis of the *CLUAP1* locus and all significant associated phenotypes**

For each phenotype: (A) Plot illustrating any potential colocalization between phenotype-significant variants and eQTLs for the given gene (B) A depiction of the genomic region surrounding the locus of interest (C) Enrichment of eQTLs for the given gene among trait-significant variants. The p-value of enrichment was determined by Fisher's exact test (D) P-P plot illustrating correlation between $p_{eQTL}$ and $p_{trait}$ for the gene of interest. Correlation between the two probabilities is visualized by plotting a best-fit linear regression over the points, with the line equation displayed on the plot. The Pearson correlation coefficient and p-value of correlation are displayed on the plot as well.

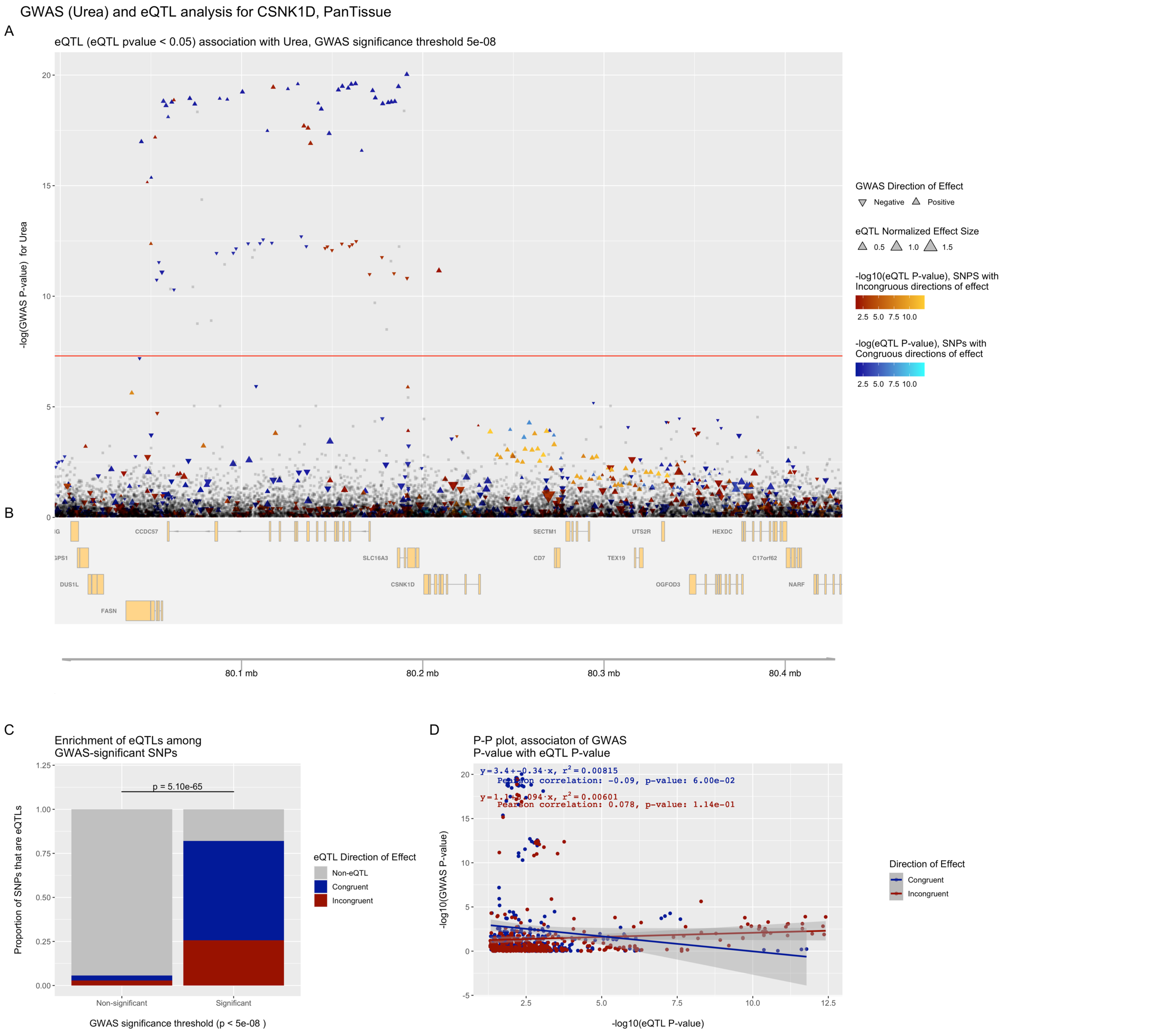


**Figure S9. eQTpLots analysis of the *CSNK1D* locus and all significant associated phenotypes**

For each phenotype: (A) Plot illustrating any potential colocalization between phenotype-significant variants and eQTLs for the given gene (B) A depiction of the genomic region surrounding the locus of interest (C) Enrichment of eQTLs for the given gene among trait-significant variants. The p-value of enrichment was determined by Fisher's exact test (D) P-P plot illustrating correlation between $p_{eQTL}$ and $p_{trait}$ for the gene of interest. Correlation between the two probabilities is visualized by plotting a best-fit linear regression over the points, with the line equation displayed on the plot. The Pearson correlation coefficient and p-value of correlation are displayed on the plot as well.

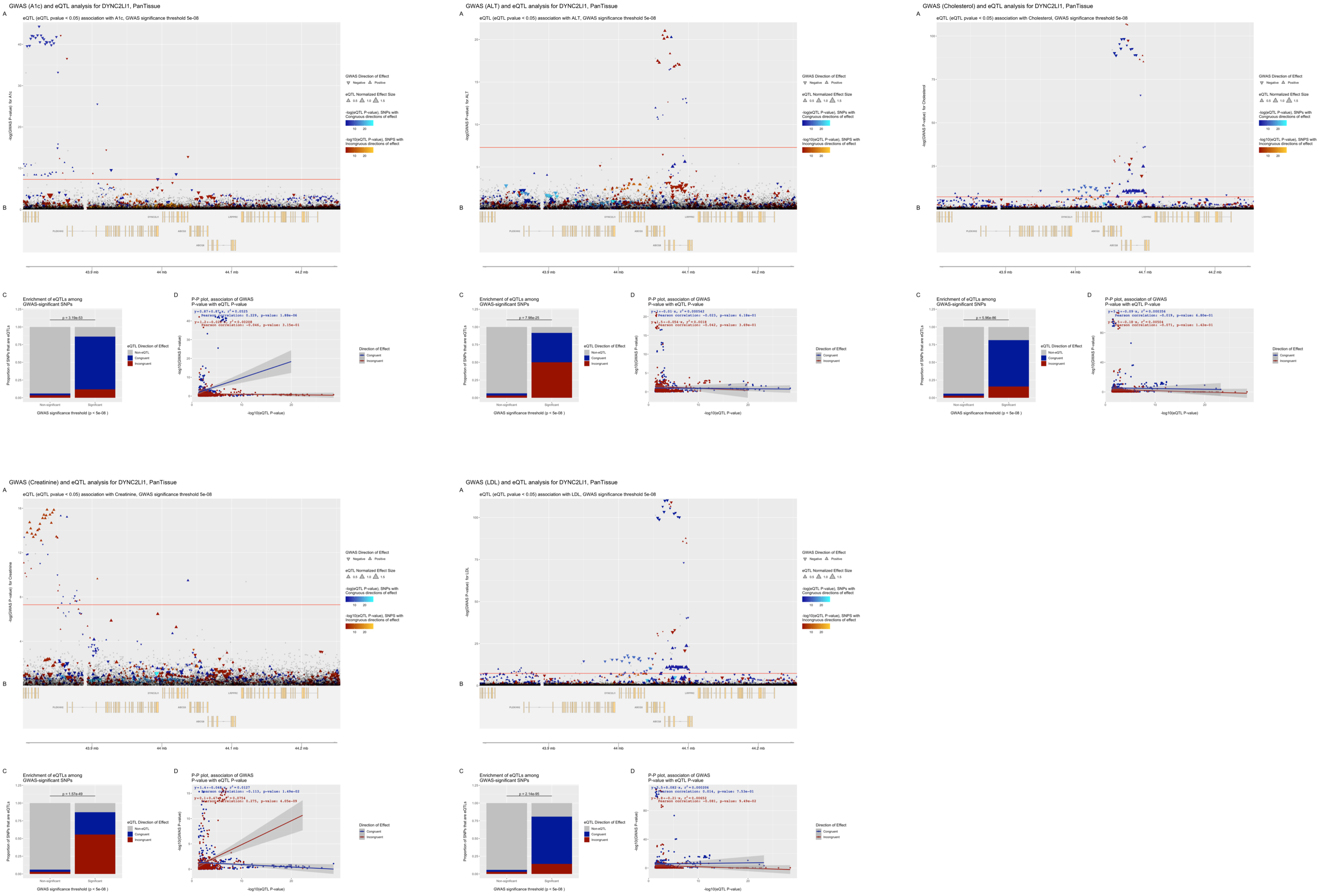


**Figure S10. eQTpLots analysis of the *DYNC2LI1* locus and all significant associated phenotypes**

For each phenotype: (A) Plot illustrating any potential colocalization between phenotype-significant variants and eQTLs for the given gene (B) A depiction of the genomic region surrounding the locus of interest (C) Enrichment of eQTLs for the given gene among trait-significant variants. The p-value of enrichment was determined by Fisher's exact test (D) P-P plot illustrating correlation between $p_{eQTL}$ and $p_{trait}$ for the gene of interest. Correlation between the two probabilities is visualized by plotting a best-fit linear regression over the points, with the line equation displayed on the plot. The Pearson correlation coefficient and p-value of correlation are displayed on the plot as well.

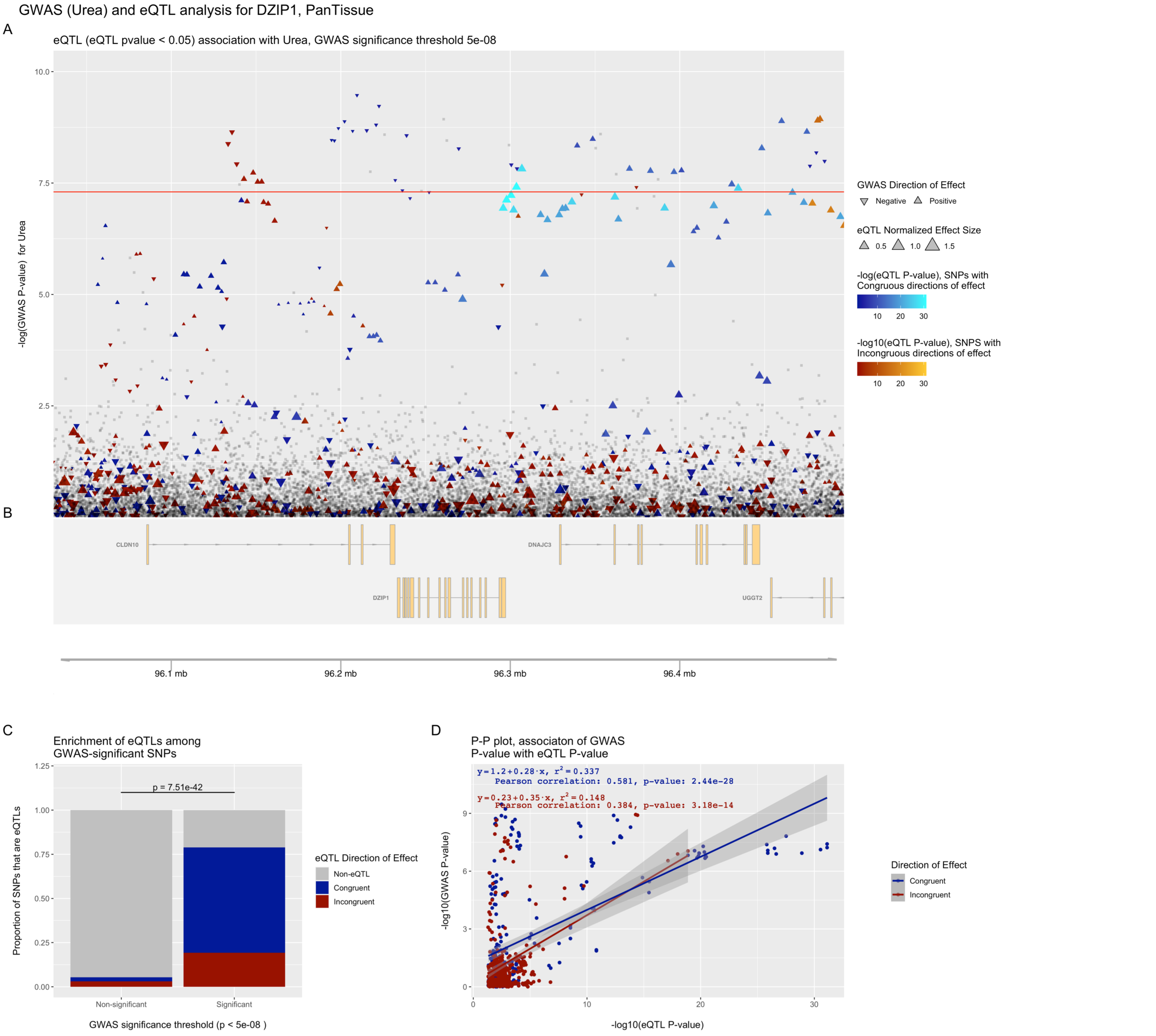


**Figure S11. eQTpLots analysis of the *DZIP1* locus and all significant associated phenotypes**

For each phenotype: (A) Plot illustrating any potential colocalization between phenotype-significant variants and eQTLs for the given gene (B) A depiction of the genomic region surrounding the locus of interest (C) Enrichment of eQTLs for the given gene among trait-significant variants. The p-value of enrichment was determined by Fisher's exact test (D) P-P plot illustrating correlation between $p_{eQTL}$ and $p_{trait}$ for the gene of interest. Correlation between the two probabilities is visualized by plotting a best-fit linear regression over the points, with the line equation displayed on the plot. The Pearson correlation coefficient and p-value of correlation are displayed on the plot as well.

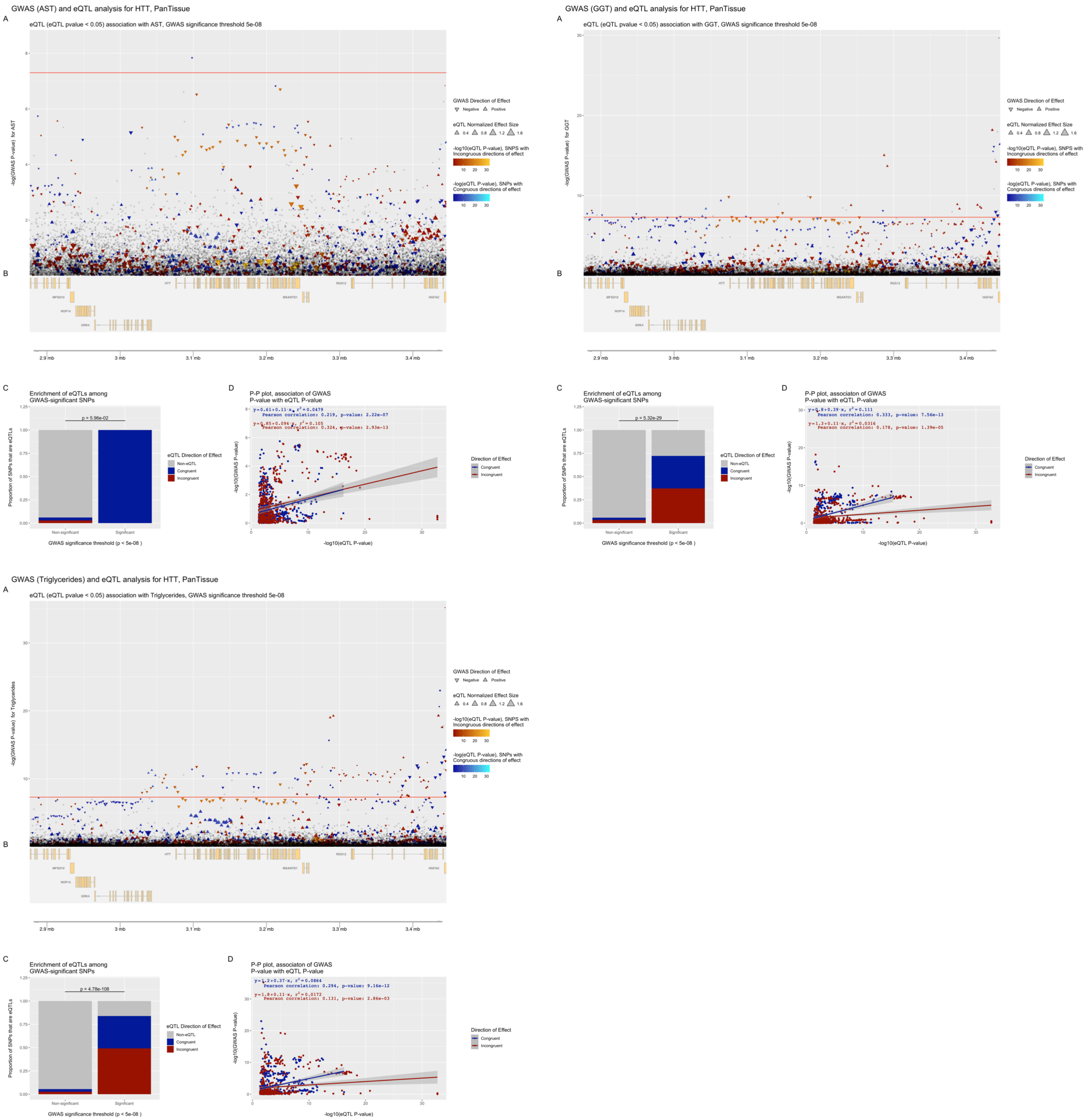


**Figure S12. eQTpLots analysis of the *HHT* locus and all significant associated phenotypes**

For each phenotype: (A) Plot illustrating any potential colocalization between phenotype-significant variants and eQTLs for the given gene (B) A depiction of the genomic region surrounding the locus of interest (C) Enrichment of eQTLs for the given gene among trait-significant variants. The p-value of enrichment was determined by Fisher's exact test (D) P-P plot illustrating correlation between $p_{eQTL}$ and $p_{trait}$ for the gene of interest. Correlation between the two probabilities is visualized by plotting a best-fit linear regression over the points, with the line equation displayed on the plot. The Pearson correlation coefficient and p-value of correlation are displayed on the plot as well.

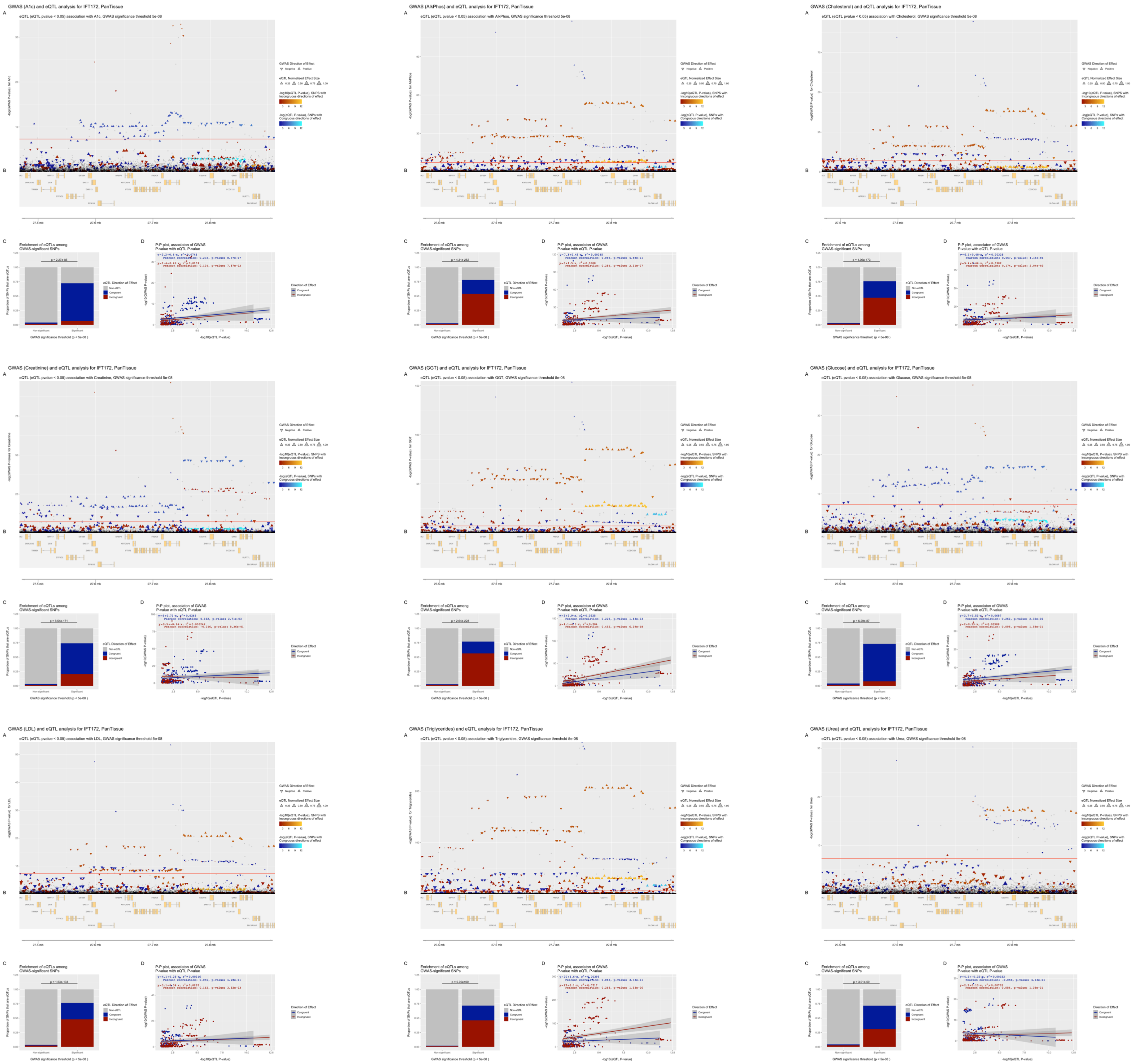


**Figure S13. eQTpLots analysis of the *IFT172* locus and all significant associated phenotypes**

For each phenotype: (A) Plot illustrating any potential colocalization between phenotype-significant variants and eQTLs for the given gene (B) A depiction of the genomic region surrounding the locus of interest (C) Enrichment of eQTLs for the given gene among trait-significant variants. The p-value of enrichment was determined by Fisher's exact test (D) P-P plot illustrating correlation between $p_{eQTL}$ and $p_{trait}$ for the gene of interest. Correlation between the two probabilities is visualized by plotting a best-fit linear regression over the points, with the line equation displayed on the plot. The Pearson correlation coefficient and p-value of correlation are displayed on the plot as well.

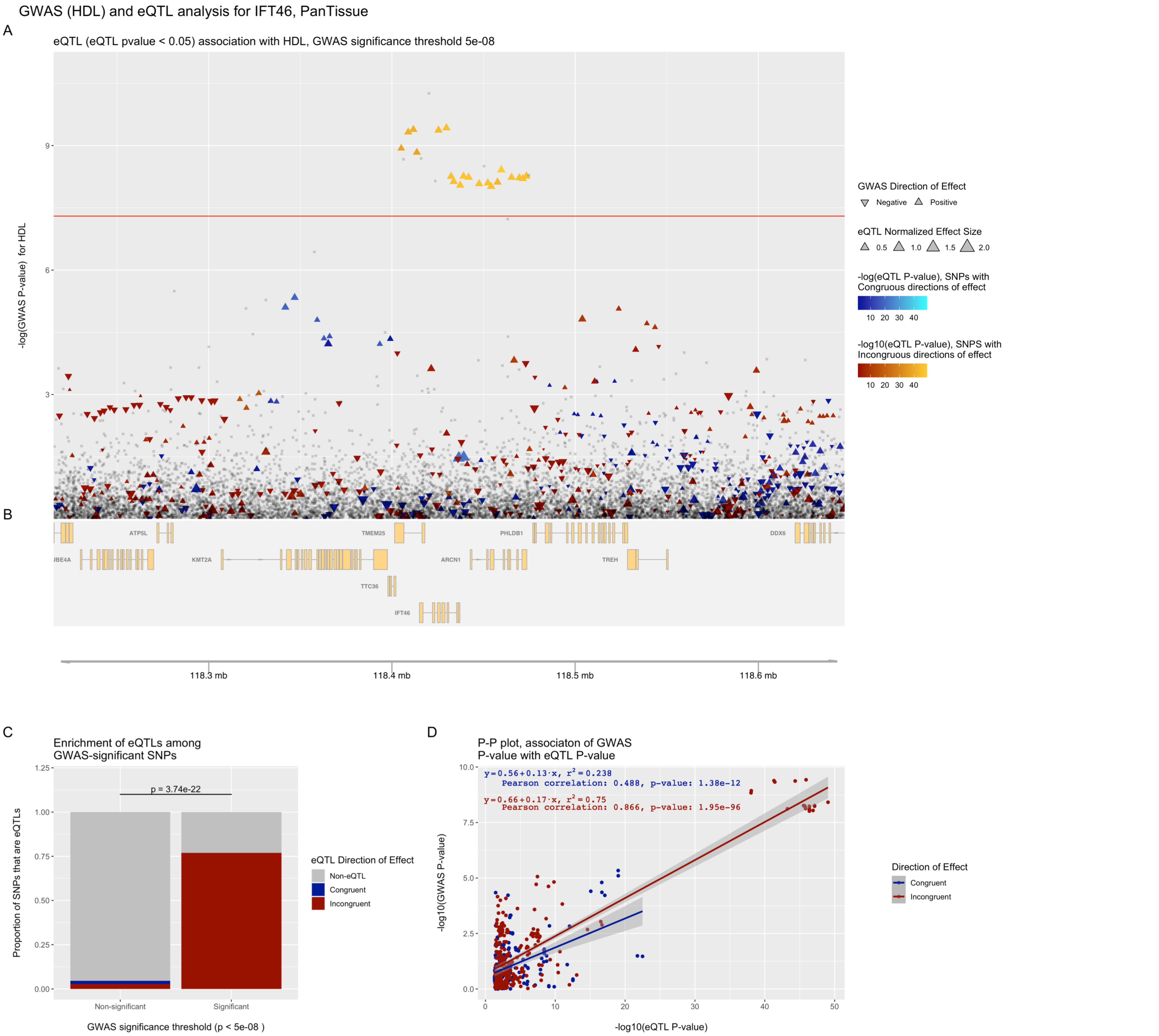


**Figure S14. eQTpLots analysis of the *IFT46* locus and all significant associated phenotypes**

For each phenotype: (A) Plot illustrating any potential colocalization between phenotype-significant variants and eQTLs for the given gene (B) A depiction of the genomic region surrounding the locus of interest (C) Enrichment of eQTLs for the given gene among trait-significant variants. The p-value of enrichment was determined by Fisher's exact test (D) P-P plot illustrating correlation between $p_{eQTL}$ and $p_{trait}$ for the gene of interest. Correlation between the two probabilities is visualized by plotting a best-fit linear regression over the points, with the line equation displayed on the plot. The Pearson correlation coefficient and p-value of correlation are displayed on the plot as well.

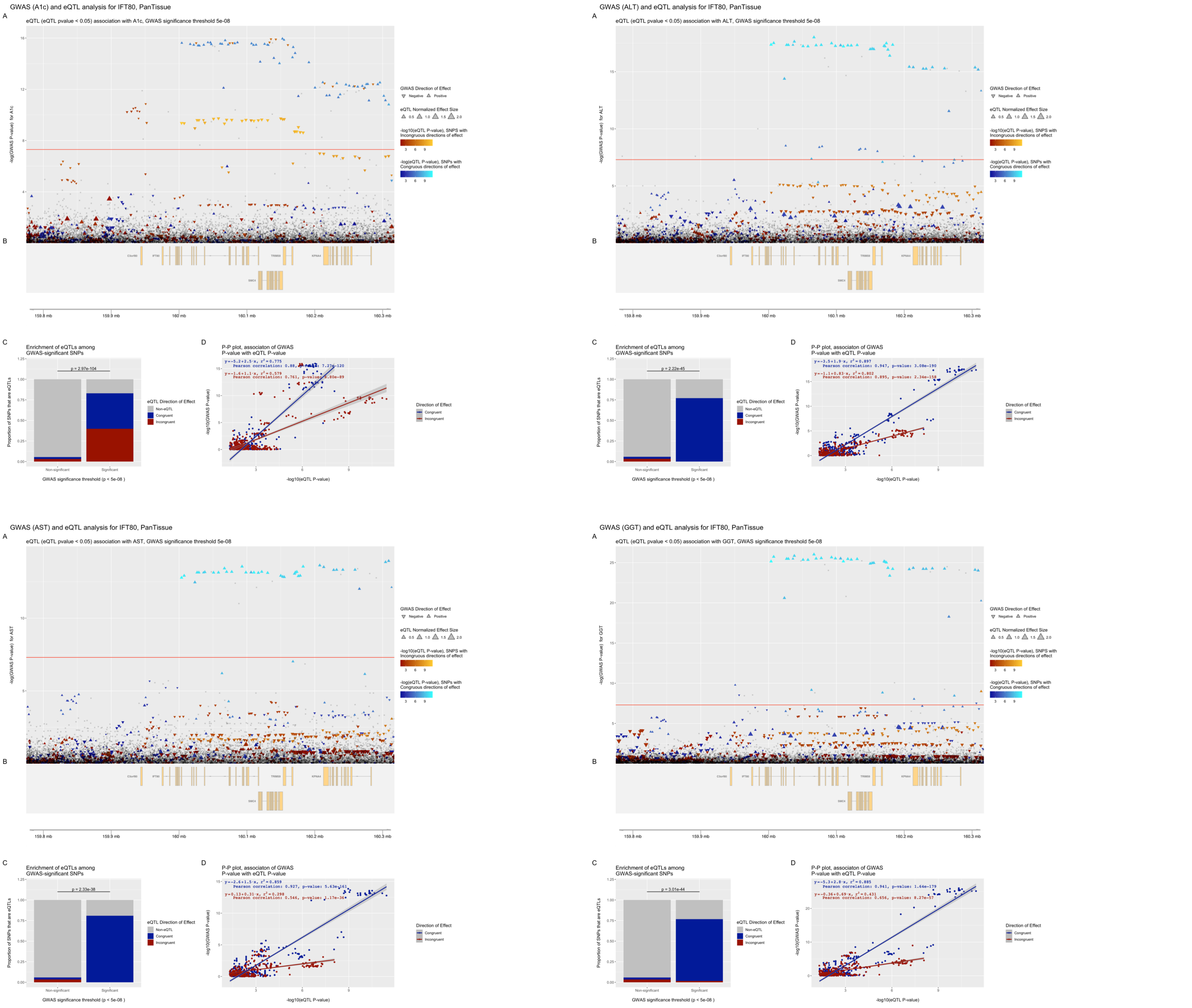


**Figure S15. eQTpLots analysis of the *IFT80* locus and all significant associated phenotypes**

For each phenotype: (A) Plot illustrating any potential colocalization between phenotype-significant variants and eQTLs for the given gene (B) A depiction of the genomic region surrounding the locus of interest (C) Enrichment of eQTLs for the given gene among trait-significant variants. The p-value of enrichment was determined by Fisher's exact test (D) P-P plot illustrating correlation between $p_{eQTL}$ and $p_{trait}$ for the gene of interest. Correlation between the two probabilities is visualized by plotting a best-fit linear regression over the points, with the line equation displayed on the plot. The Pearson correlation coefficient and p-value of correlation are displayed on the plot as well.

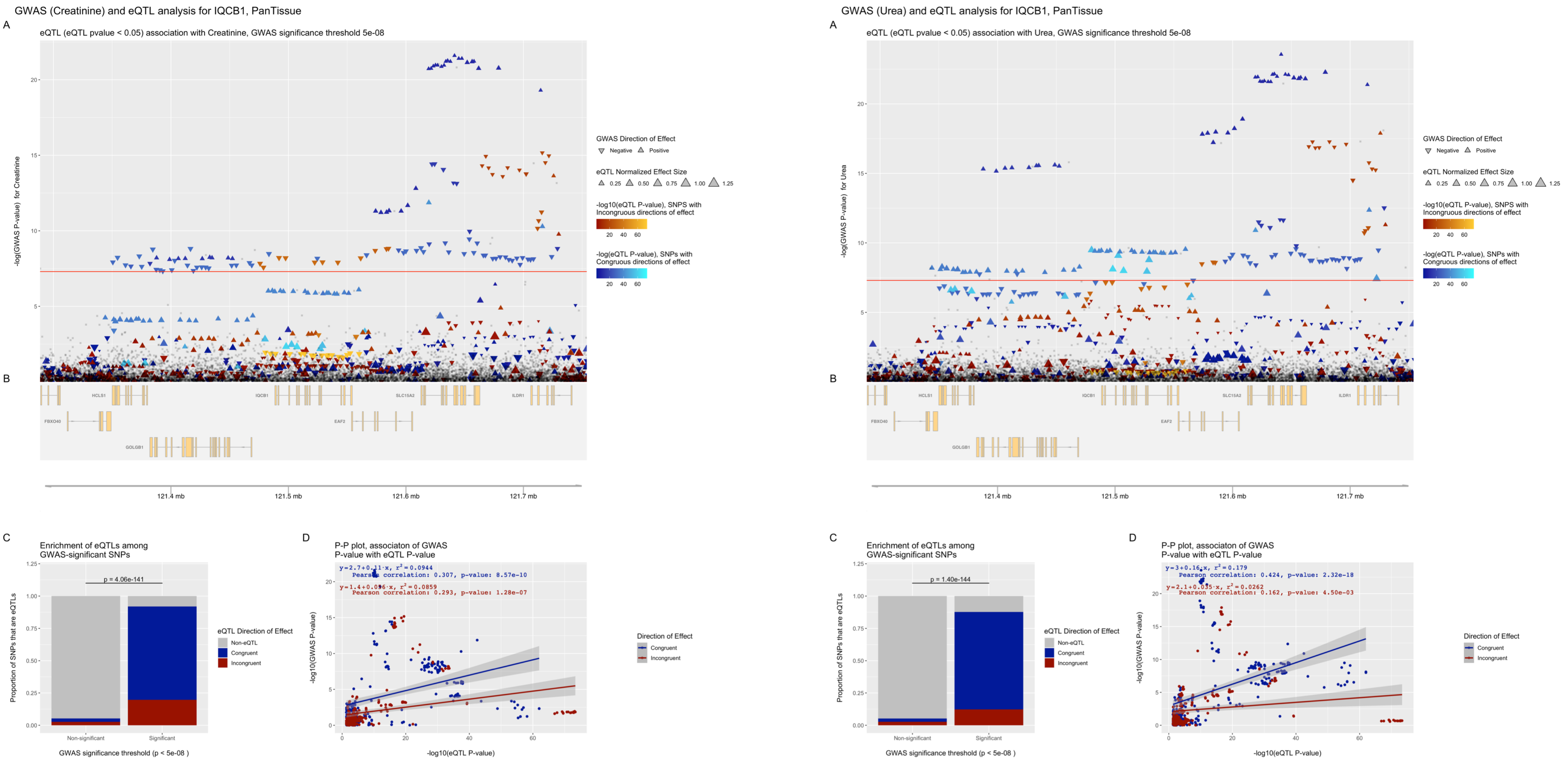


**Figure S16. eQTpLots analysis of the *IQCB1* locus and all significant associated phenotypes**
For each phenotype: (A) Plot illustrating any potential colocalization between phenotype-significant variants and eQTLs for the given gene (B) A depiction of the genomic region surrounding the locus of interest (C) Enrichment of eQTLs for the given gene among trait-significant variants. The p-value of enrichment was determined by Fisher’s exact test (D) P-P plot illustrating correlation between $p_{eQTL}$ and $p_{trait}$ for the gene of interest. Correlation between the two probabilities is visualized by plotting a best-fit linear regression over the points, with the line equation displayed on the plot. The Pearson correlation coefficient and p-value of correlation are displayed on the plot as well.

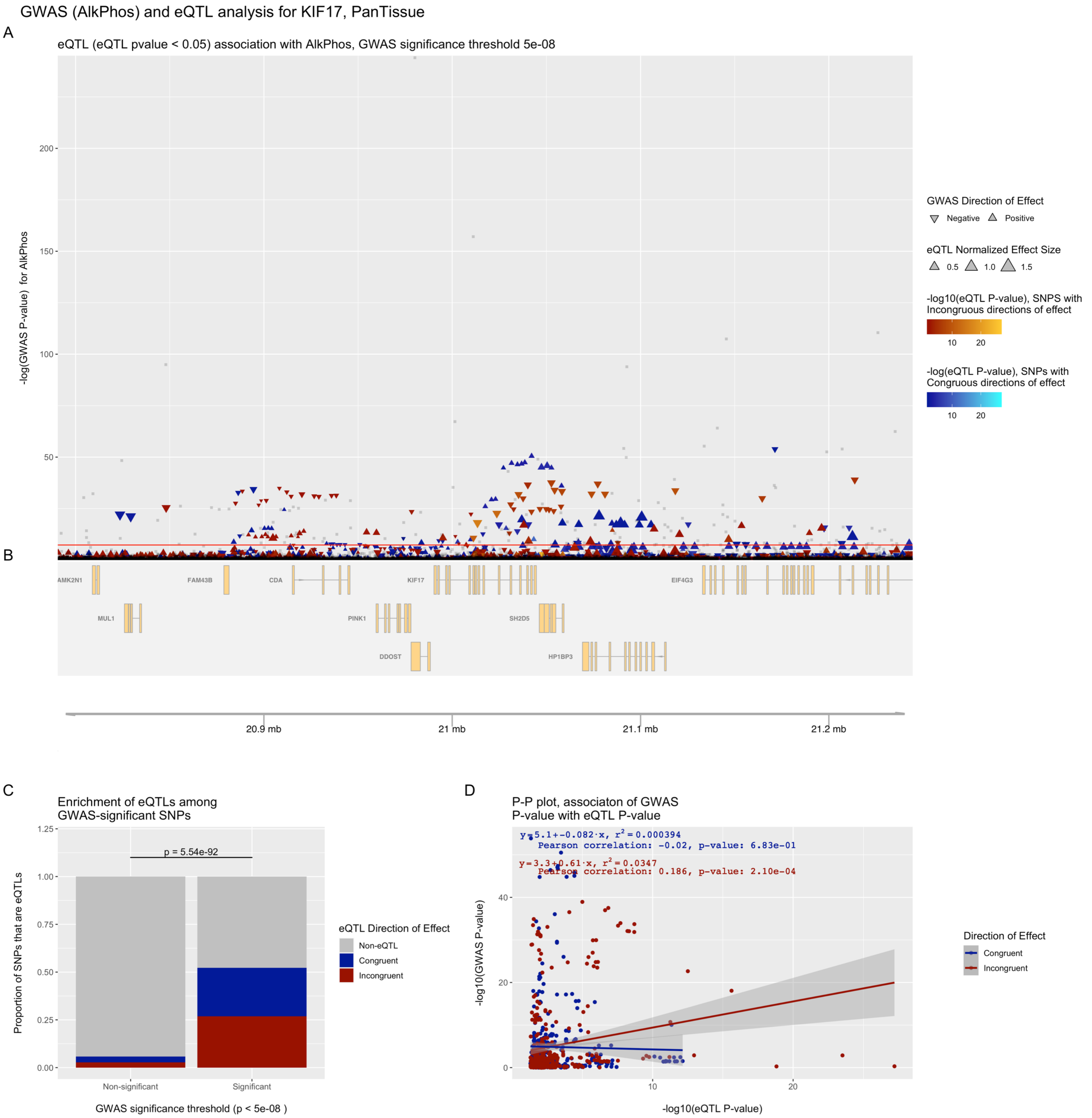


**Figure S17. eQTpLots analysis of the *KIF17* locus and all significant associated phenotypes**

For each phenotype: (A) Plot illustrating any potential colocalization between phenotype-significant variants and eQTLs for the given gene (B) A depiction of the genomic region surrounding the locus of interest (C) Enrichment of eQTLs for the given gene among trait-significant variants. The p-value of enrichment was determined by Fisher's exact test (D) P-P plot illustrating correlation between $p_{eQTL}$ and $p_{trait}$ for the gene of interest. Correlation between the two probabilities is visualized by plotting a best-fit linear regression over the points, with the line equation displayed on the plot. The Pearson correlation coefficient and p-value of correlation are displayed on the plot as well.

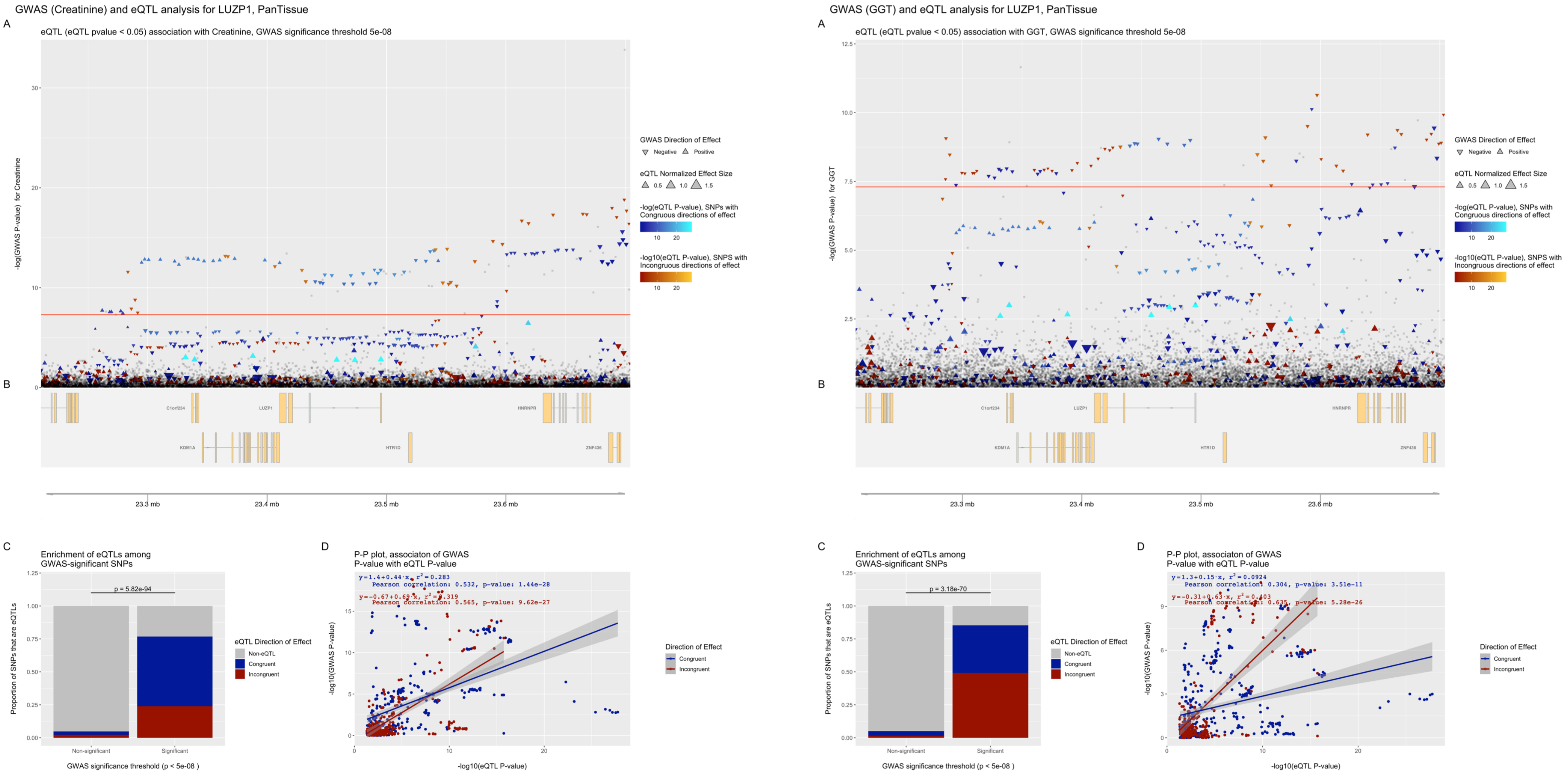


**Figure S18. eQTpLots analysis of the *LUZP1* locus and all significant associated phenotypes**

For each phenotype: (A) Plot illustrating any potential colocalization between phenotype-significant variants and eQTLs for the given gene (B) A depiction of the genomic region surrounding the locus of interest (C) Enrichment of eQTLs for the given gene among trait-significant variants. The p-value of enrichment was determined by Fisher's exact test (D) P-P plot illustrating correlation between $p_{eQTL}$ and $p_{trait}$ for the gene of interest. Correlation between the two probabilities is visualized by plotting a best-fit linear regression over the points, with the line equation displayed on the plot. The Pearson correlation coefficient and p-value of correlation are displayed on the plot as well.

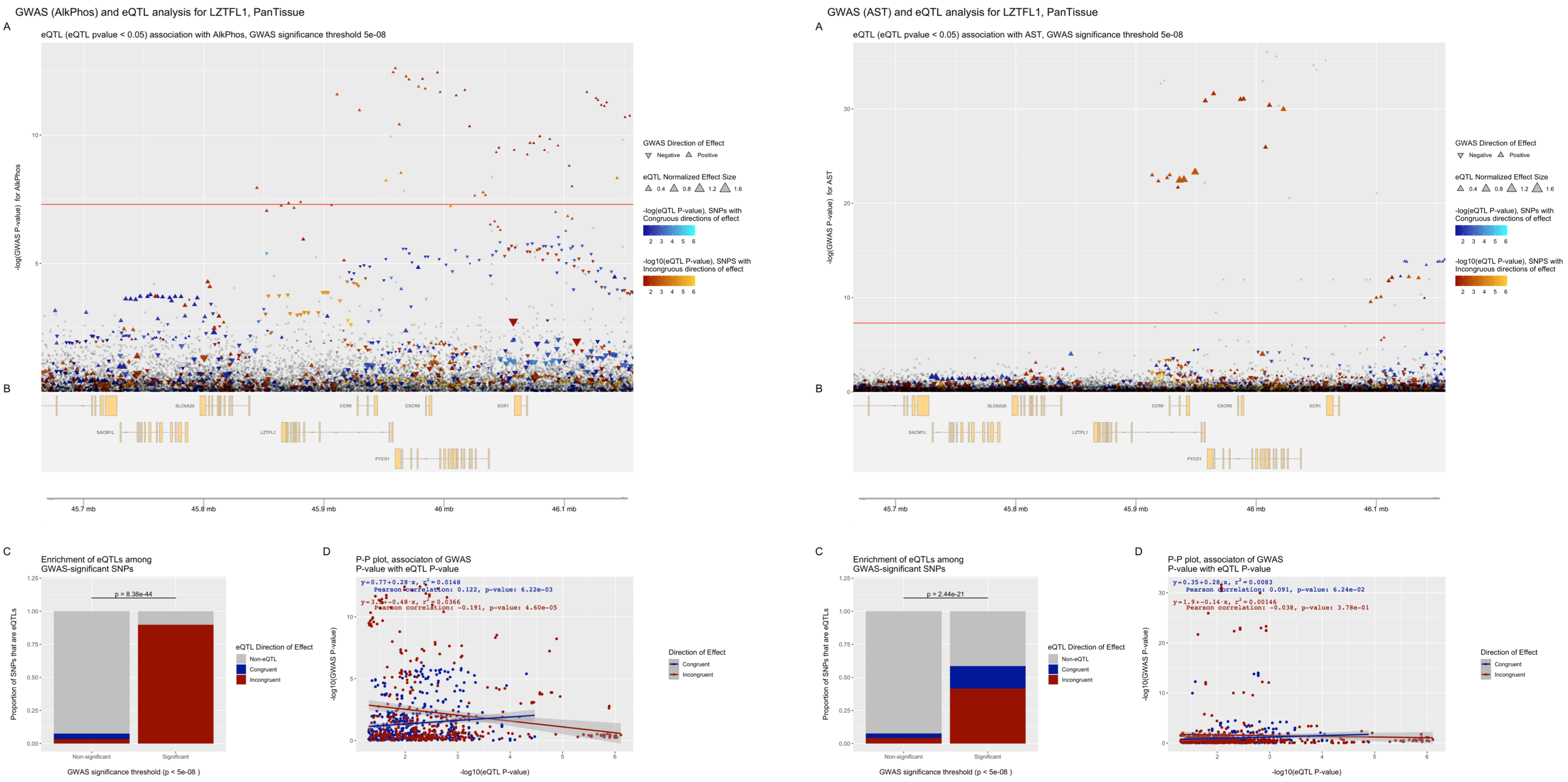


**Figure S19. eQTpLots analysis of the *LZTFL1* locus and all significant associated phenotypes**

For each phenotype: (A) Plot illustrating any potential colocalization between phenotype-significant variants and eQTLs for the given gene (B) A depiction of the genomic region surrounding the locus of interest (C) Enrichment of eQTLs for the given gene among trait-significant variants. The p-value of enrichment was determined by Fisher's exact test (D) P-P plot illustrating correlation between $p_{eQTL}$ and $p_{trait}$ for the gene of interest. Correlation between the two probabilities is visualized by plotting a best-fit linear regression over the points, with the line equation displayed on the plot. The Pearson correlation coefficient and p-value of correlation are displayed on the plot as well.

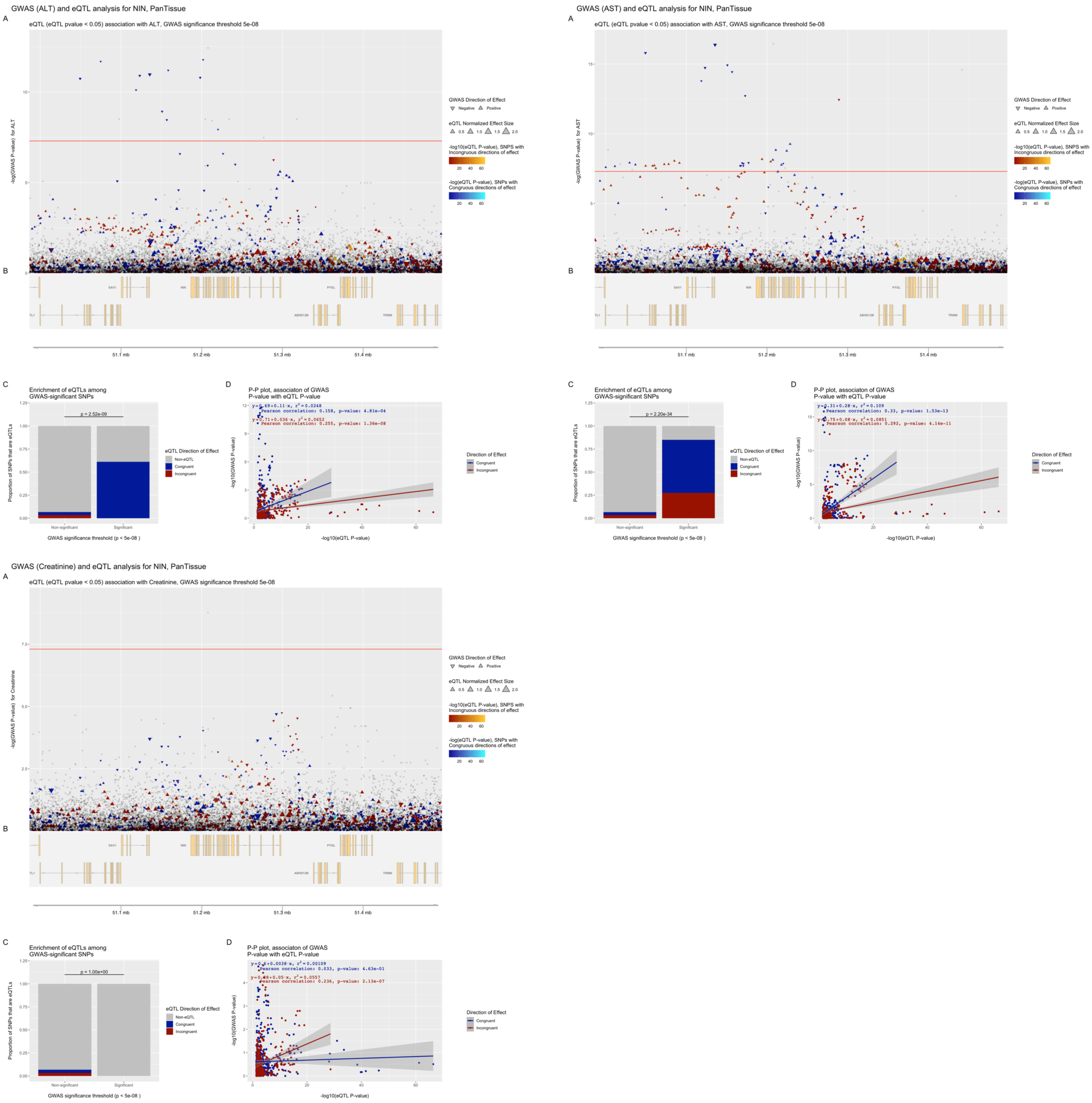


**Figure S20. eQTpLots analysis of the *NIN* locus and all significant associated phenotypes**

For each phenotype: (A) Plot illustrating any potential colocalization between phenotype-significant variants and eQTLs for the given gene (B) A depiction of the genomic region surrounding the locus of interest (C) Enrichment of eQTLs for the given gene among trait-significant variants. The p-value of enrichment was determined by Fisher's exact test (D) P-P plot illustrating correlation between $p_{eQTL}$ and $p_{trait}$ for the gene of interest. Correlation between the two probabilities is visualized by plotting a best-fit linear regression over the points, with the line equation displayed on the plot. The Pearson correlation coefficient and p-value of correlation are displayed on the plot as well.

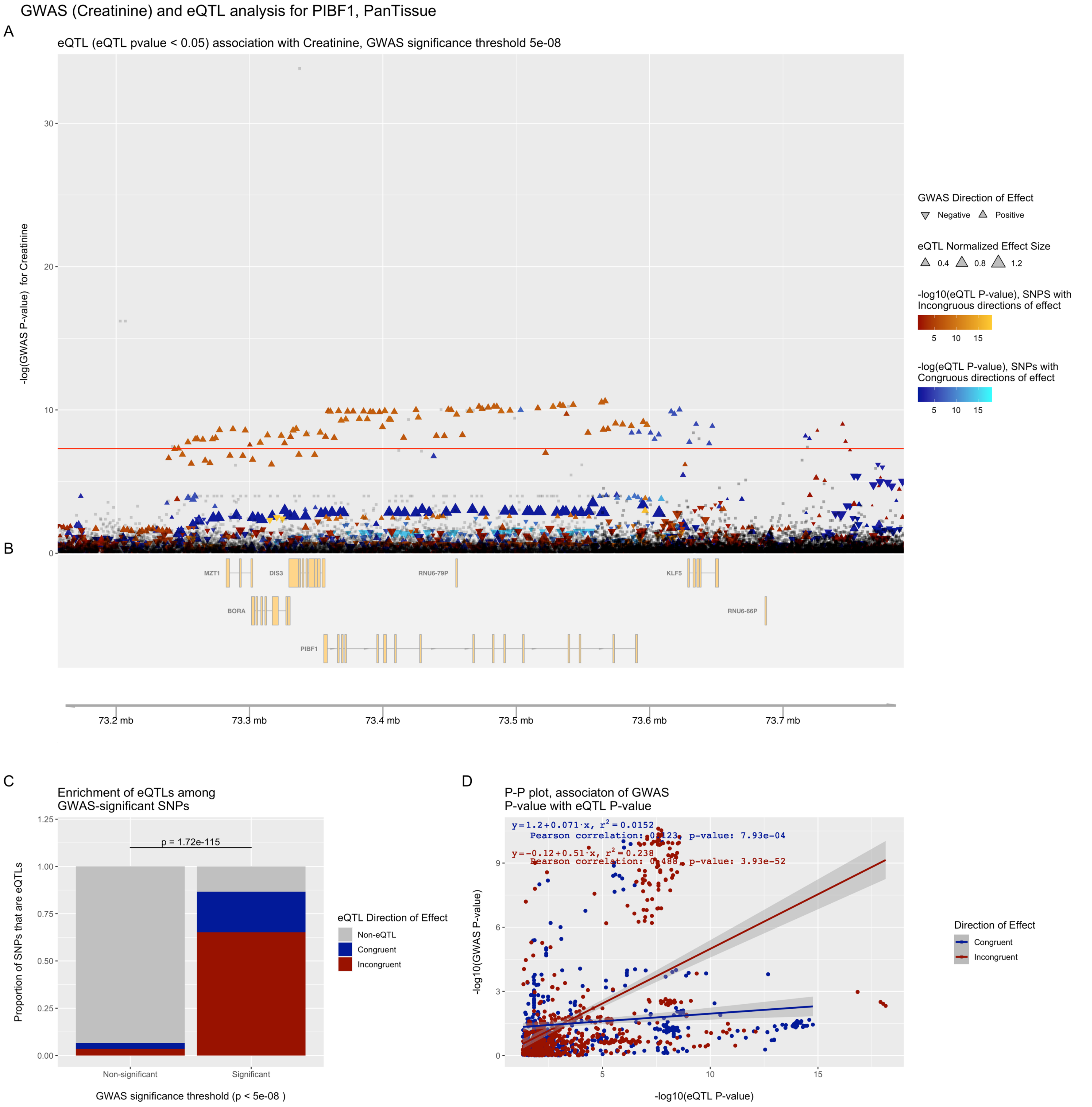


**Figure S21. eQTpLots analysis of the *PIBF1* locus and all significant associated phenotypes**

For each phenotype: (A) Plot illustrating any potential colocalization between phenotype-significant variants and eQTLs for the given gene (B) A depiction of the genomic region surrounding the locus of interest (C) Enrichment of eQTLs for the given gene among trait-significant variants. The p-value of enrichment was determined by Fisher's exact test (D) P-P plot illustrating correlation between $p_{eQTL}$ and $p_{trait}$ for the gene of interest. Correlation between the two probabilities is visualized by plotting a best-fit linear regression over the points, with the line equation displayed on the plot. The Pearson correlation coefficient and p-value of correlation are displayed on the plot as well.

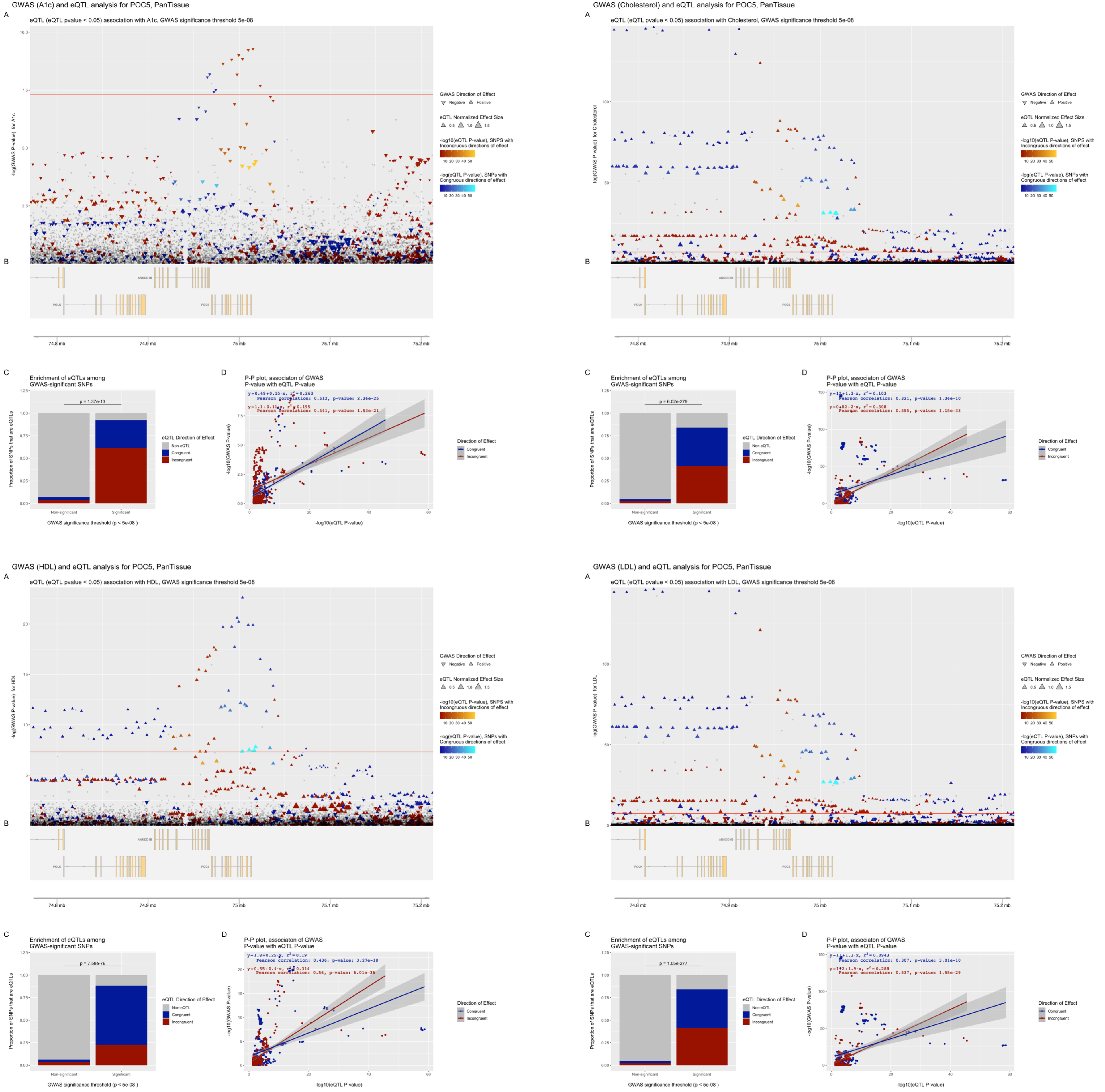


**Figure S22. eQTpLots analysis of the *POC5* locus and all significant associated phenotypes**

For each phenotype: (A) Plot illustrating any potential colocalization between phenotype-significant variants and eQTLs for the given gene (B) A depiction of the genomic region surrounding the locus of interest (C) Enrichment of eQTLs for the given gene among trait-significant variants. The p-value of enrichment was determined by Fisher's exact test (D) P-P plot illustrating correlation between $p_{eQTL}$ and $p_{trait}$ for the gene of interest. Correlation between the two probabilities is visualized by plotting a best-fit linear regression over the points, with the line equation displayed on the plot. The Pearson correlation coefficient and p-value of correlation are displayed on the plot as well.

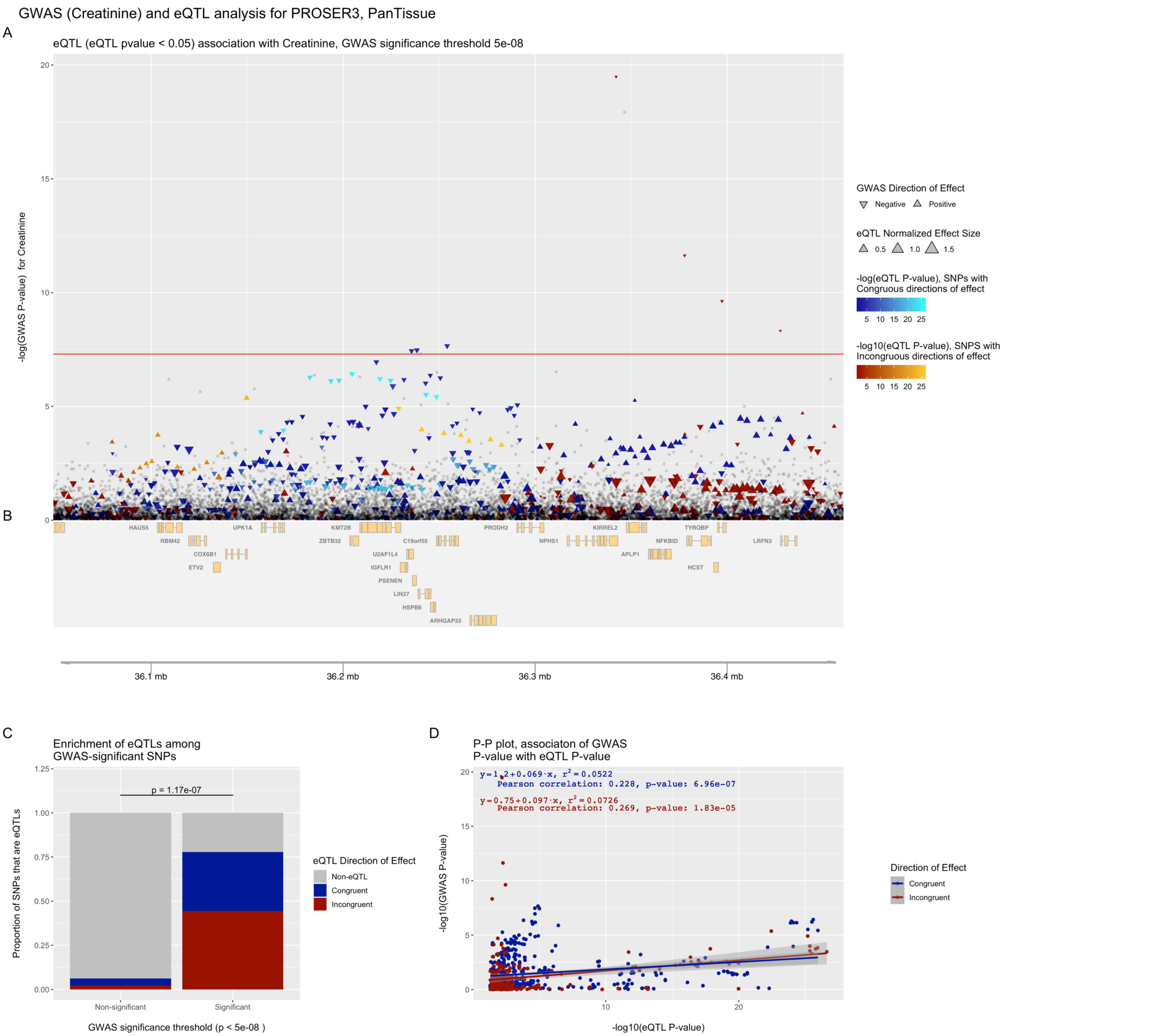


**Figure S23. eQTpLots analysis of the *PROSER3* locus and all significant associated phenotypes**

For each phenotype: (A) Plot illustrating any potential colocalization between phenotype-significant variants and eQTLs for the given gene (B) A depiction of the genomic region surrounding the locus of interest (C) Enrichment of eQTLs for the given gene among trait-significant variants. The p-value of enrichment was determined by Fisher's exact test (D) P-P plot illustrating correlation between $p_{eQTL}$ and $p_{trait}$ for the gene of interest. Correlation between the two probabilities is visualized by plotting a best-fit linear regression over the points, with the line equation displayed on the plot. The Pearson correlation coefficient and p-value of correlation are displayed on the plot as well.

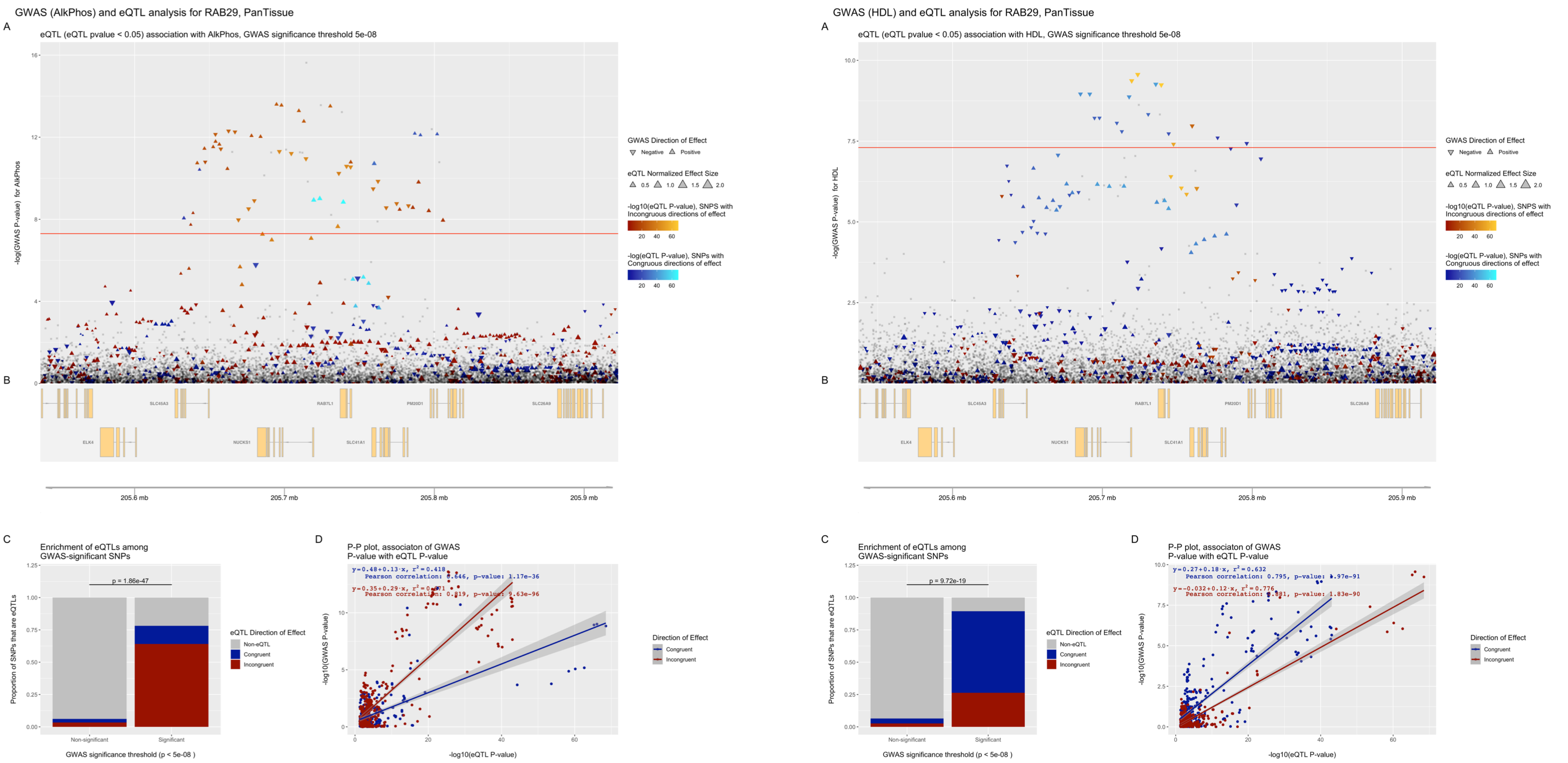


**Figure S24. eQTpLots analysis of the *RAB29* locus and all significant associated phenotypes**

For each phenotype: (A) Plot illustrating any potential colocalization between phenotype-significant variants and eQTLs for the given gene (B) A depiction of the genomic region surrounding the locus of interest (C) Enrichment of eQTLs for the given gene among trait-significant variants. The p-value of enrichment was determined by Fisher's exact test (D) P-P plot illustrating correlation between $p_{eQTL}$ and $p_{trait}$ for the gene of interest. Correlation between the two probabilities is visualized by plotting a best-fit linear regression over the points, with the line equation displayed on the plot. The Pearson correlation coefficient and p-value of correlation are displayed on the plot as well.

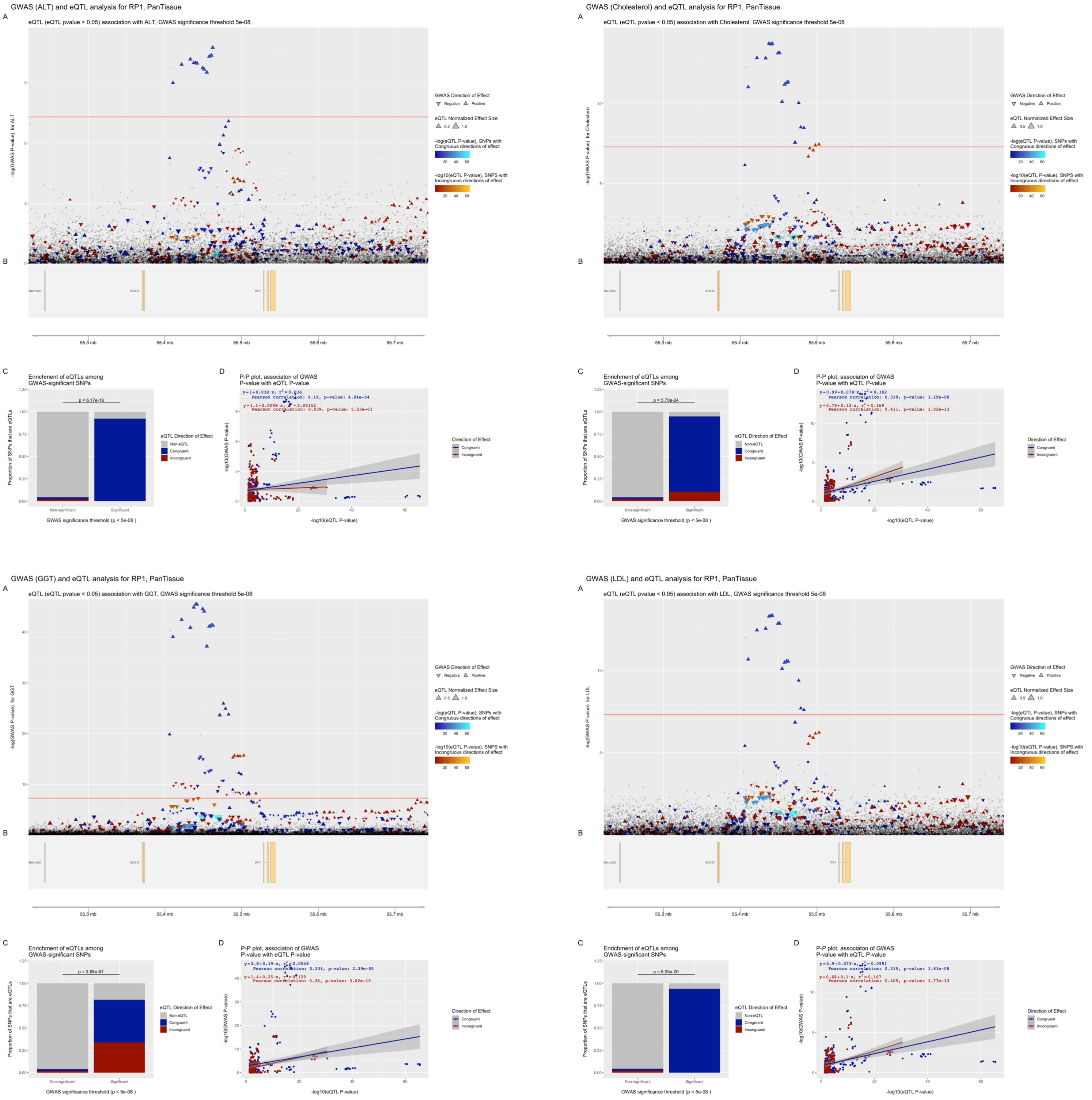


**Figure S25. eQTpLots analysis of the *RP1* locus and all significant associated phenotypes**

For each phenotype: (A) Plot illustrating any potential colocalization between phenotype-significant variants and eQTLs for the given gene (B) A depiction of the genomic region surrounding the locus of interest (C) Enrichment of eQTLs for the given gene among trait-significant variants. The p-value of enrichment was determined by Fisher’s exact test (D) P-P plot illustrating correlation between $p_{eQTL}$ and $p_{trait}$ for the gene of interest. Correlation between the two probabilities is visualized by plotting a best-fit linear regression over the points, with the line equation displayed on the plot. The Pearson correlation coefficient and p-value of correlation are displayed on the plot as well.

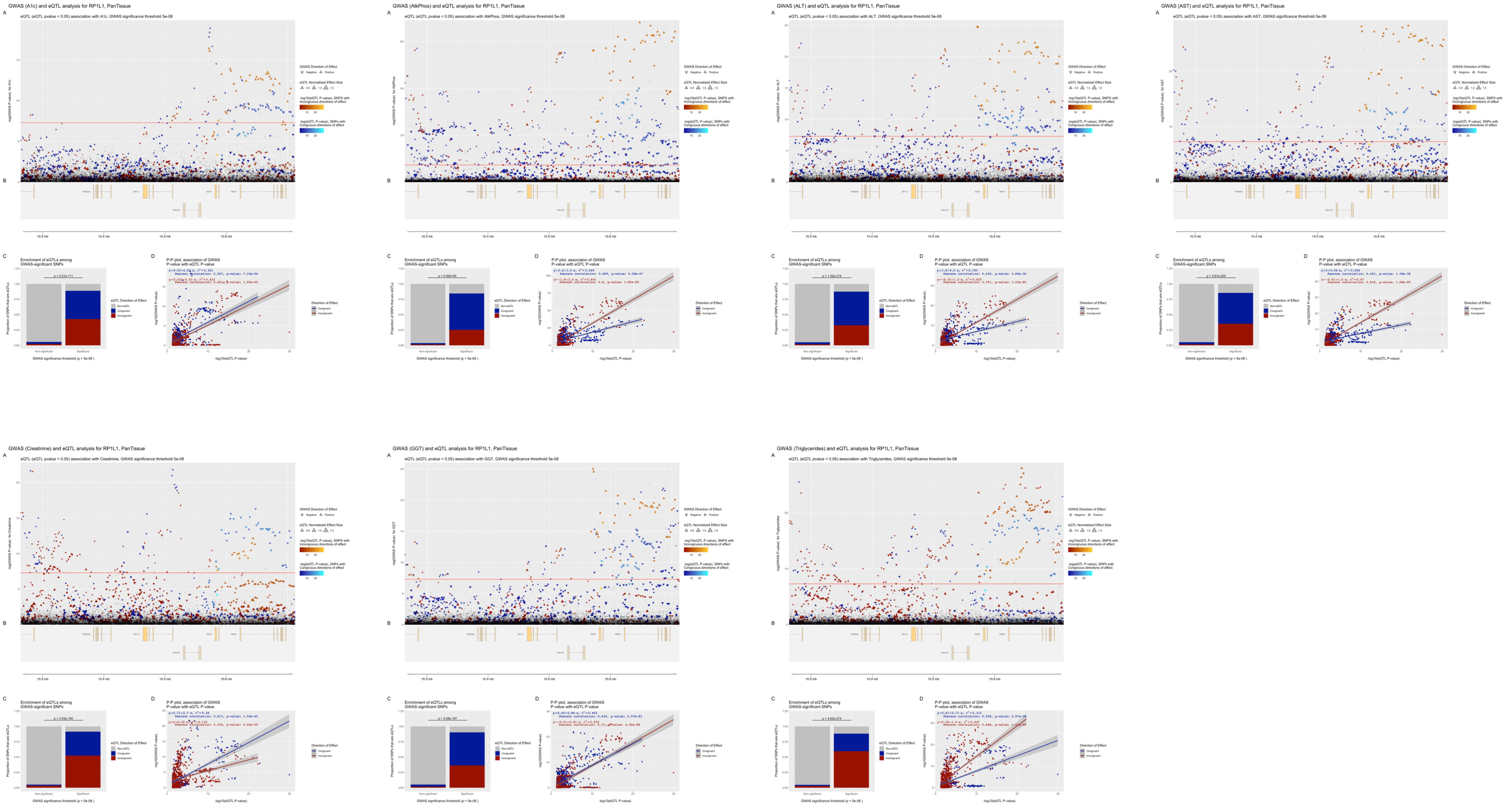


**Figure S26. eQTpLots analysis of the *RP1L1* locus and all significant associated phenotypes**

For each phenotype: (A) Plot illustrating any potential colocalization between phenotype-significant variants and eQTLs for the given gene (B) A depiction of the genomic region surrounding the locus of interest (C) Enrichment of eQTLs for the given gene among trait-significant variants. The p-value of enrichment was determined by Fisher's exact test (D) P-P plot illustrating correlation between $p_{eQTL}$ and $p_{trait}$ for the gene of interest. Correlation between the two probabilities is visualized by plotting a best-fit linear regression over the points, with the line equation displayed on the plot. The Pearson correlation coefficient and p-value of correlation are displayed on the plot as well.

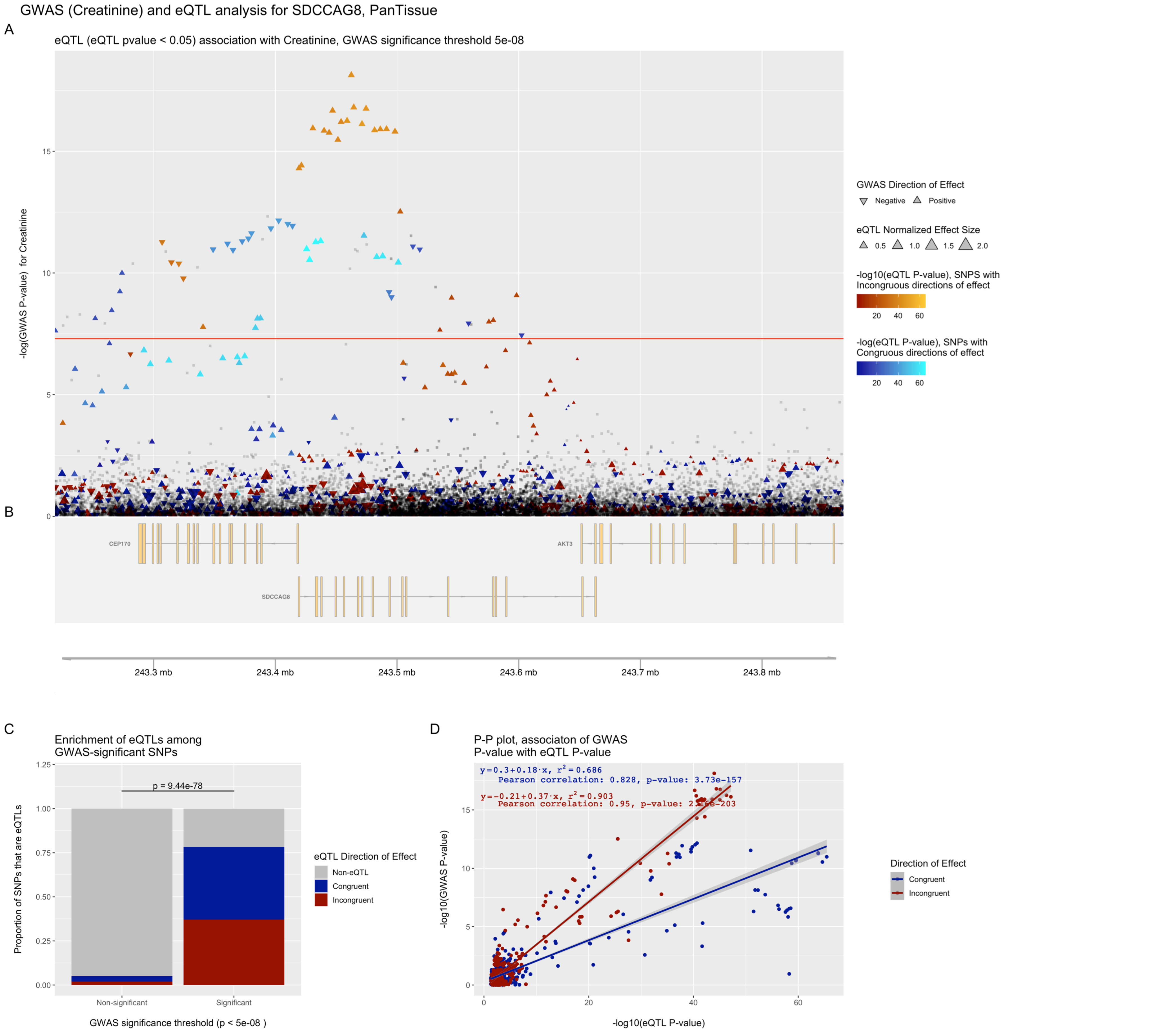


**Figure S27. eQTpLots analysis of the *SDCCAG8* locus and all significant associated phenotypes**

For each phenotype: (A) Plot illustrating any potential colocalization between phenotype-significant variants and eQTLs for the given gene (B) A depiction of the genomic region surrounding the locus of interest (C) Enrichment of eQTLs for the given gene among trait-significant variants. The p-value of enrichment was determined by Fisher's exact test (D) P-P plot illustrating correlation between $p_{eQTL}$ and $p_{trait}$ for the gene of interest. Correlation between the two probabilities is visualized by plotting a best-fit linear regression over the points, with the line equation displayed on the plot. The Pearson correlation coefficient and p-value of correlation are displayed on the plot as well.

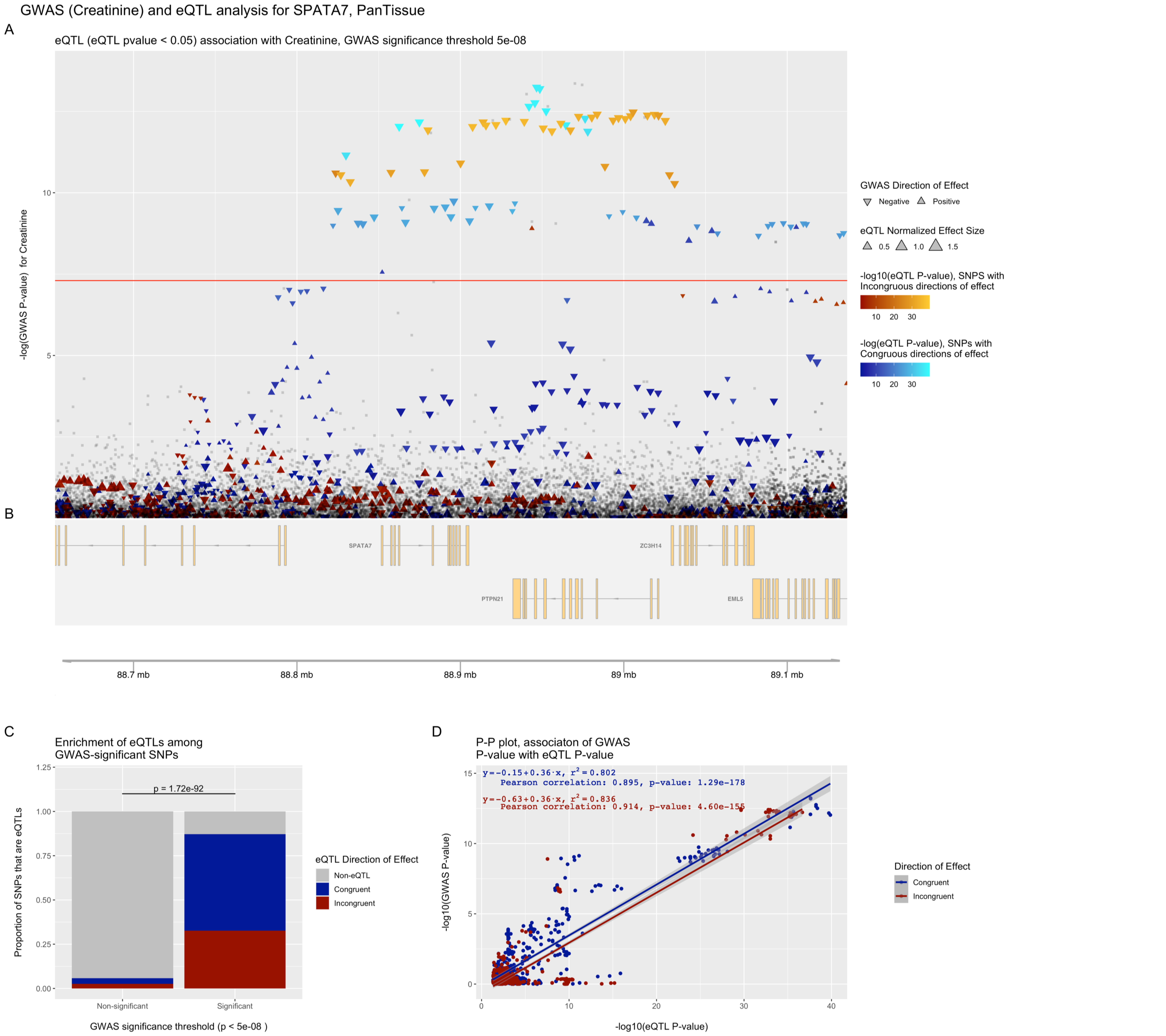


**Figure S28. eQTpLots analysis of the *SPATA7* locus and all significant associated phenotypes**

For each phenotype: (A) Plot illustrating any potential colocalization between phenotype-significant variants and eQTLs for the given gene (B) A depiction of the genomic region surrounding the locus of interest (C) Enrichment of eQTLs for the given gene among trait-significant variants. The p-value of enrichment was determined by Fisher's exact test (D) P-P plot illustrating correlation between $p_{eQTL}$ and $p_{trait}$ for the gene of interest. Correlation between the two probabilities is visualized by plotting a best-fit linear regression over the points, with the line equation displayed on the plot. The Pearson correlation coefficient and p-value of correlation are displayed on the plot as well.

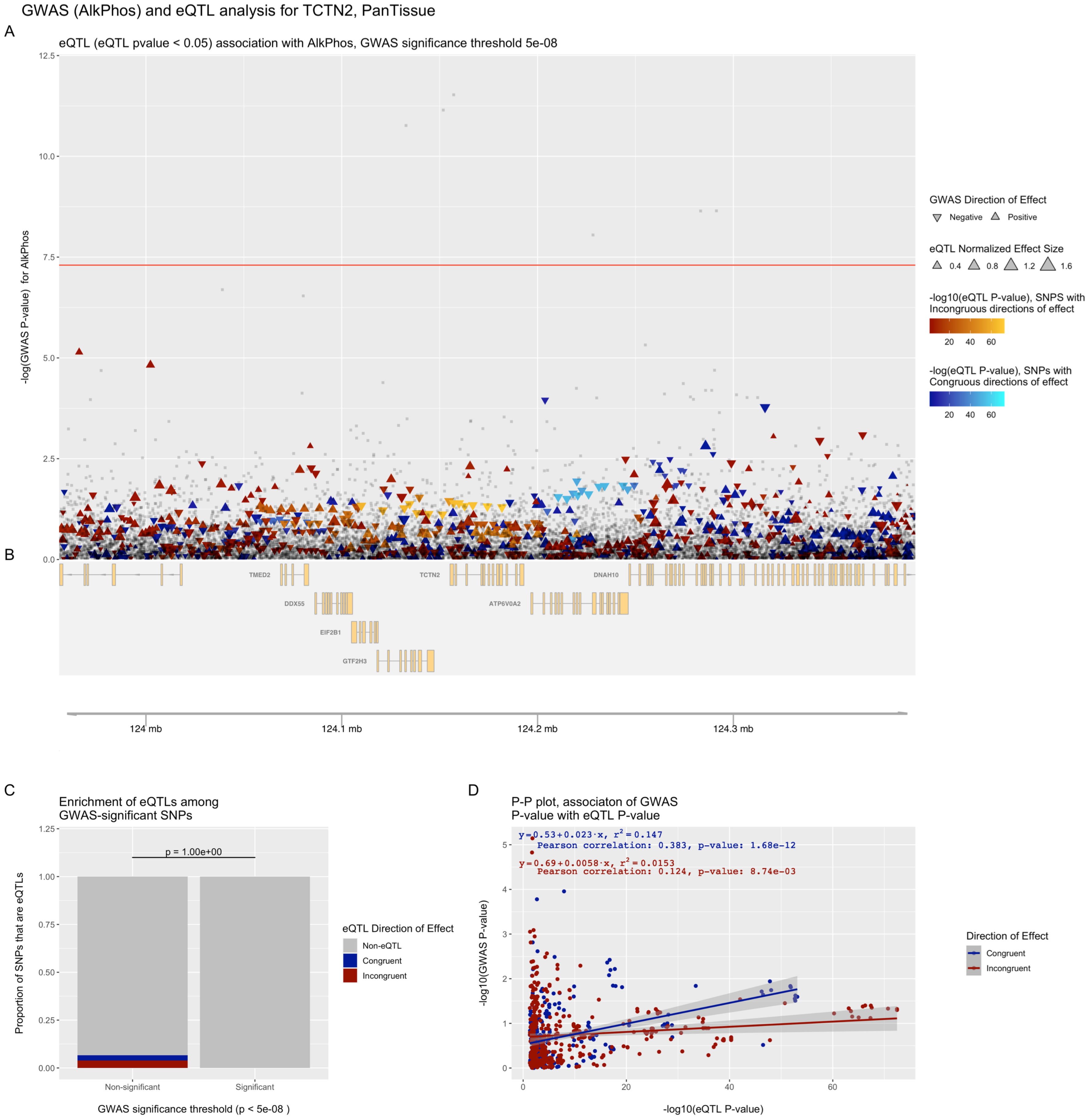


**Figure S29. eQTpLots analysis of the *TCTN2* locus and all significant associated phenotypes**

For each phenotype: (A) Plot illustrating any potential colocalization between phenotype-significant variants and eQTLs for the given gene (B) A depiction of the genomic region surrounding the locus of interest (C) Enrichment of eQTLs for the given gene among trait-significant variants. The p-value of enrichment was determined by Fisher's exact test (D) P-P plot illustrating correlation between $p_{eQTL}$ and $p_{trait}$ for the gene of interest. Correlation between the two probabilities is visualized by plotting a best-fit linear regression over the points, with the line equation displayed on the plot. The Pearson correlation coefficient and p-value of correlation are displayed on the plot as well.

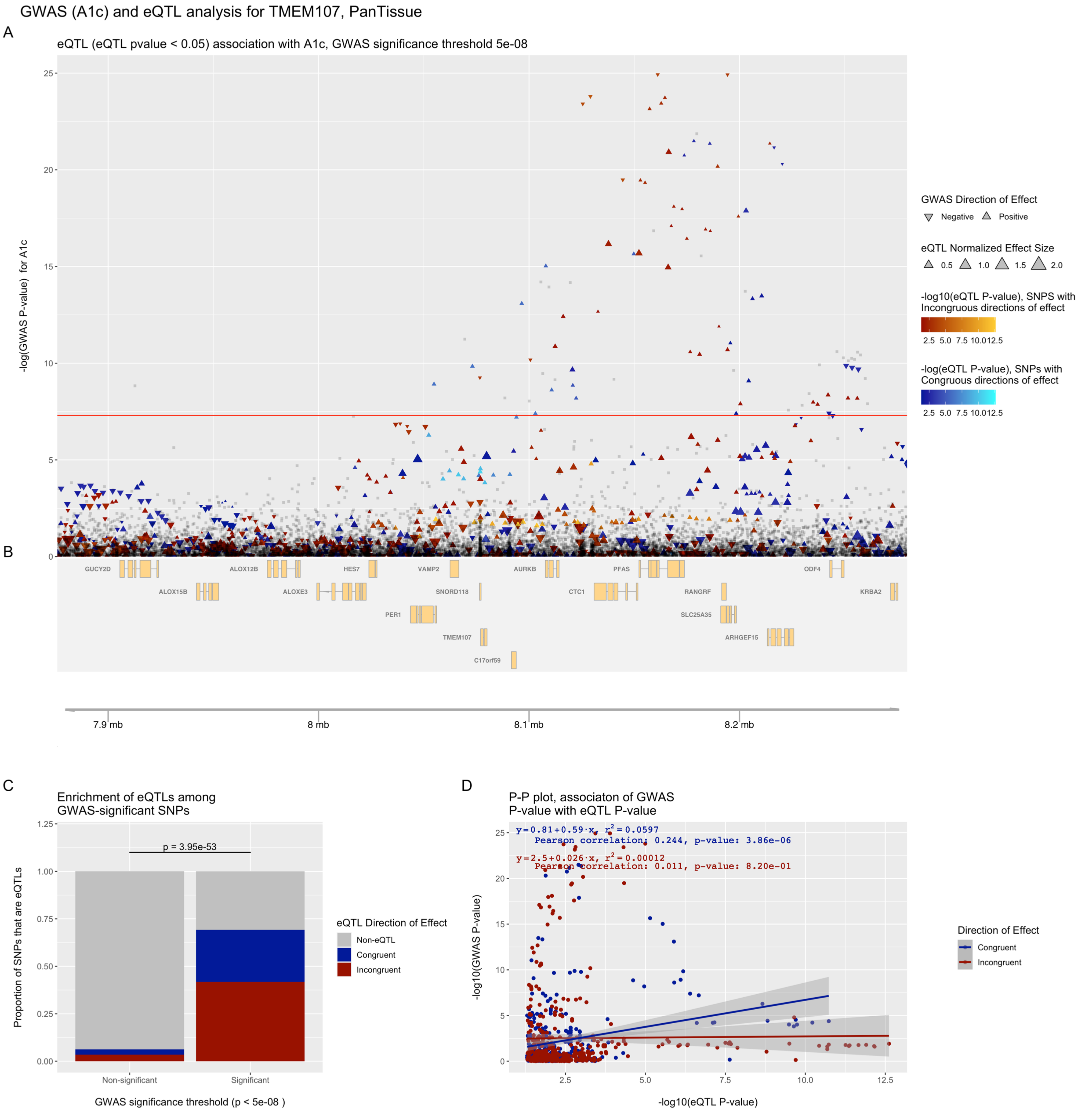


**Figure S30. eQTpLots analysis of the *TMEM107* locus and all significant associated phenotypes**

For each phenotype: (A) Plot illustrating any potential colocalization between phenotype-significant variants and eQTLs for the given gene (B) A depiction of the genomic region surrounding the locus of interest (C) Enrichment of eQTLs for the given gene among trait-significant variants. The p-value of enrichment was determined by Fisher's exact test (D) P-P plot illustrating correlation between $p_{eQTL}$ and $p_{trait}$ for the gene of interest. Correlation between the two probabilities is visualized by plotting a best-fit linear regression over the points, with the line equation displayed on the plot. The Pearson correlation coefficient and p-value of correlation are displayed on the plot as well.

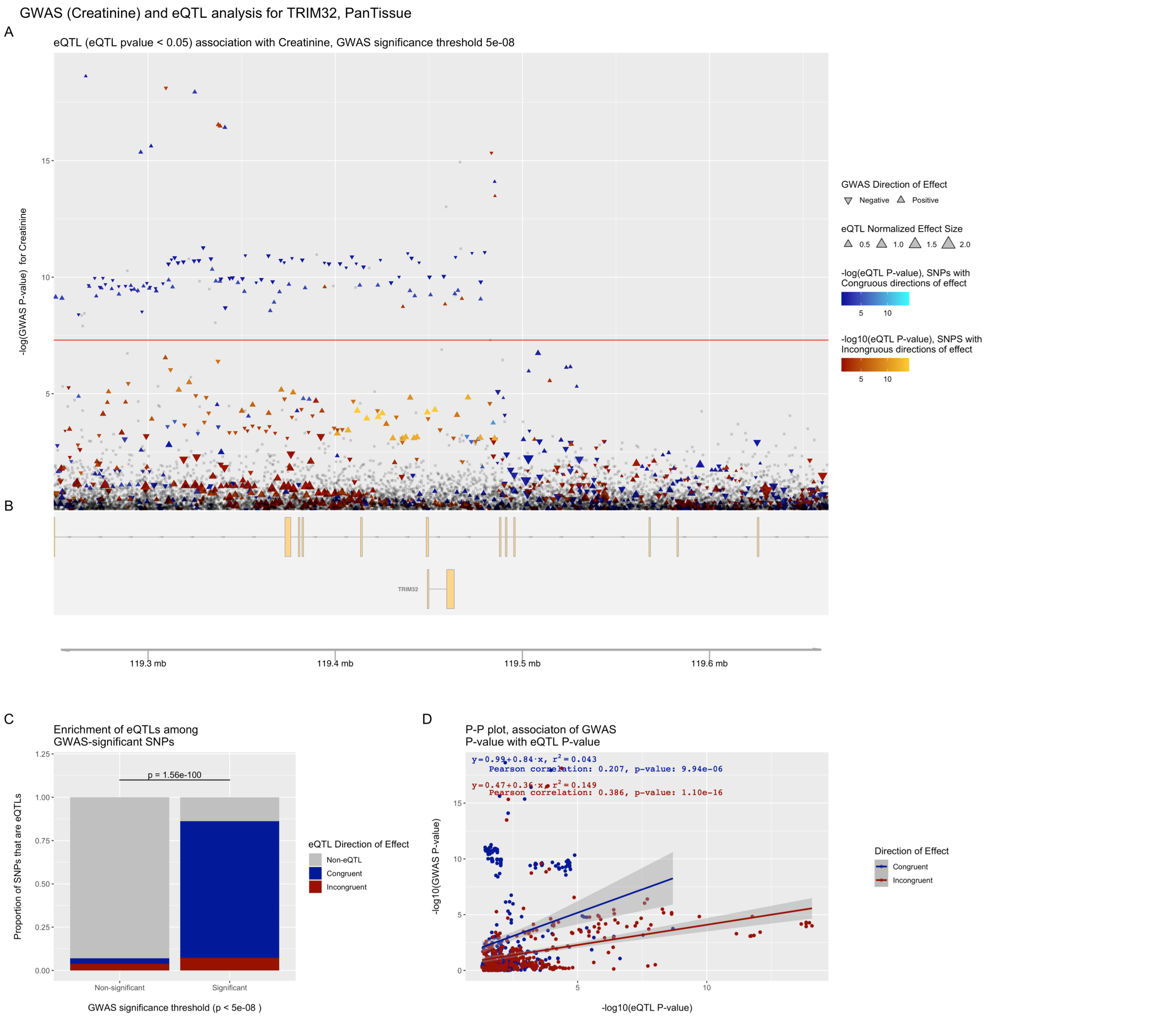


**Figure S31. eQTpLots analysis of the *TRIM32* locus and all significant associated phenotypes**

For each phenotype: (A) Plot illustrating any potential colocalization between phenotype-significant variants and eQTLs for the given gene (B) A depiction of the genomic region surrounding the locus of interest (C) Enrichment of eQTLs for the given gene among trait-significant variants. The p-value of enrichment was determined by Fisher's exact test (D) P-P plot illustrating correlation between $p_{eQTL}$ and $p_{trait}$ for the gene of interest. Correlation between the two probabilities is visualized by plotting a best-fit linear regression over the points, with the line equation displayed on the plot. The Pearson correlation coefficient and p-value of correlation are displayed on the plot as well.

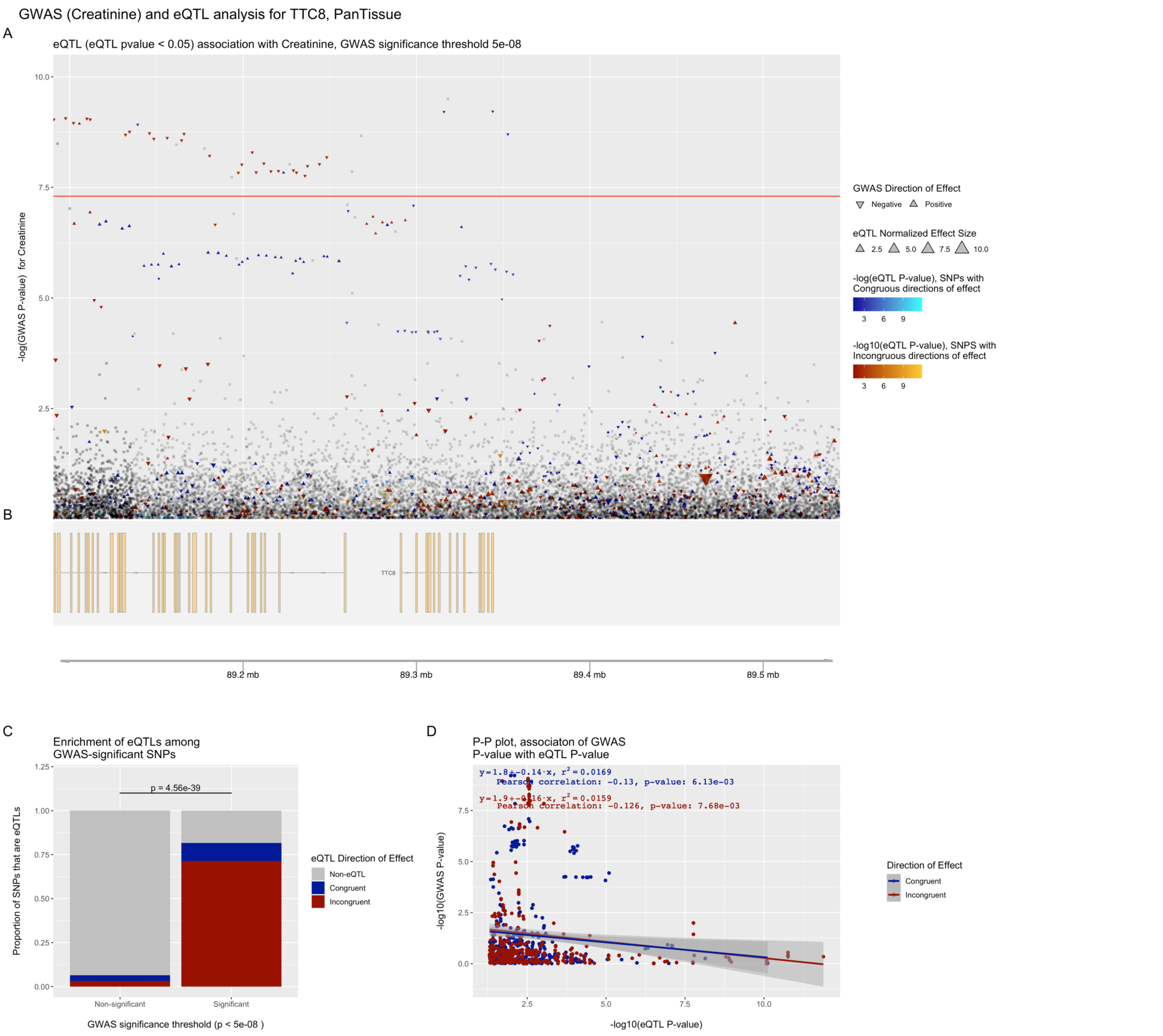


**Figure S32. eQTpLots analysis of the *TTC8* locus and all significant associated phenotypes**

For each phenotype: (A) Plot illustrating any potential colocalization between phenotype-significant variants and eQTLs for the given gene (B) A depiction of the genomic region surrounding the locus of interest (C) Enrichment of eQTLs for the given gene among trait-significant variants. The p-value of enrichment was determined by Fisher's exact test (D) P-P plot illustrating correlation between $p_{eQTL}$ and $p_{trait}$ for the gene of interest. Correlation between the two probabilities is visualized by plotting a best-fit linear regression over the points, with the line equation displayed on the plot. The Pearson correlation coefficient and p-value of correlation are displayed on the plot as well.

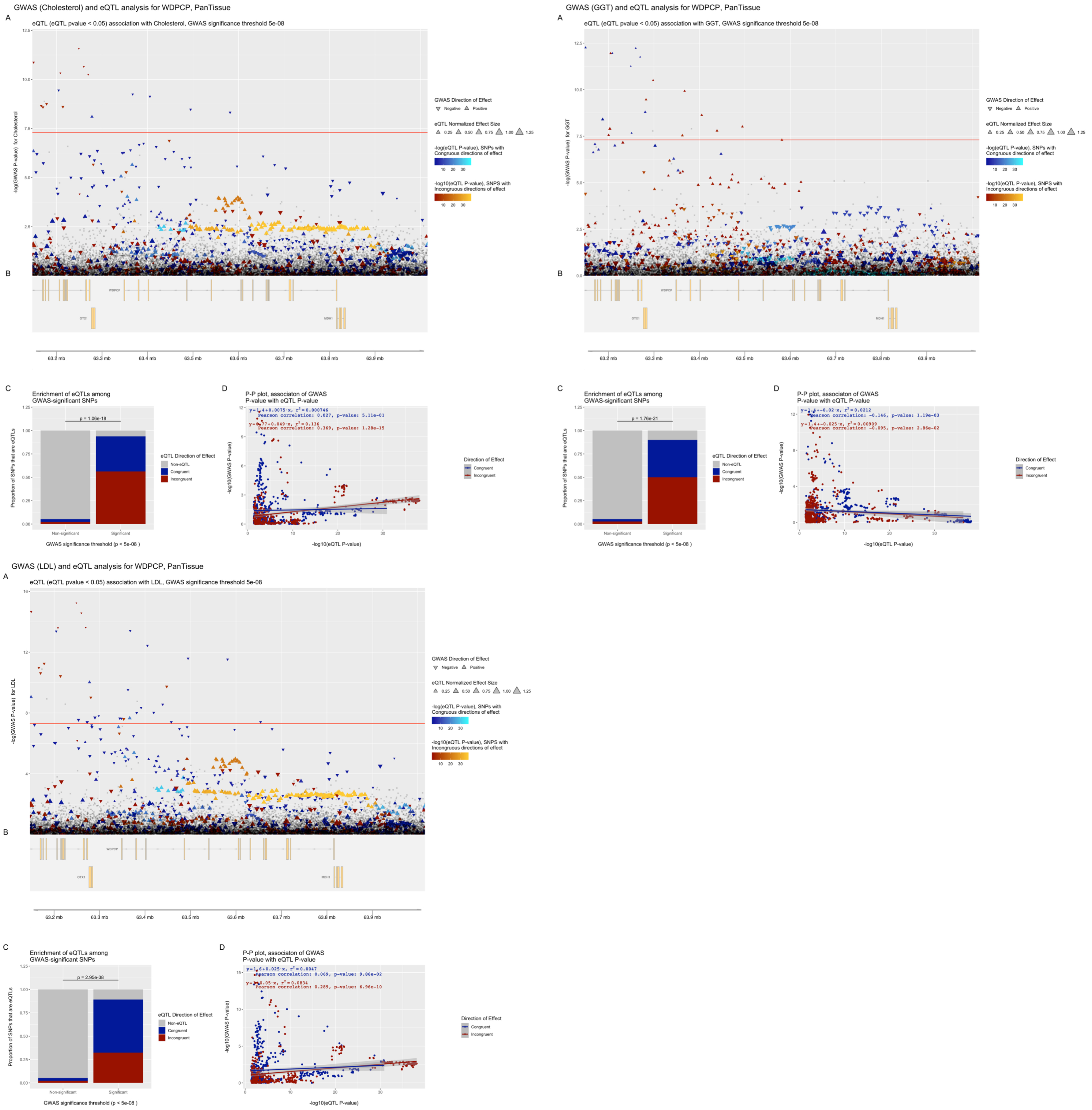


**Figure S33. eQTpLots analysis of the *WDPCP* locus and all significant associated phenotypes**

For each phenotype: (A) Plot illustrating any potential colocalization between phenotype-significant variants and eQTLs for the given gene (B) A depiction of the genomic region surrounding the locus of interest (C) Enrichment of eQTLs for the given gene among trait-significant variants. The p-value of enrichment was determined by Fisher's exact test (D) P-P plot illustrating correlation between $p_{eQTL}$ and $p_{trait}$ for the gene of interest. Correlation between the two probabilities is visualized by plotting a best-fit linear regression over the points, with the line equation displayed on the plot. The Pearson correlation coefficient and p-value of correlation are displayed on the plot as well.

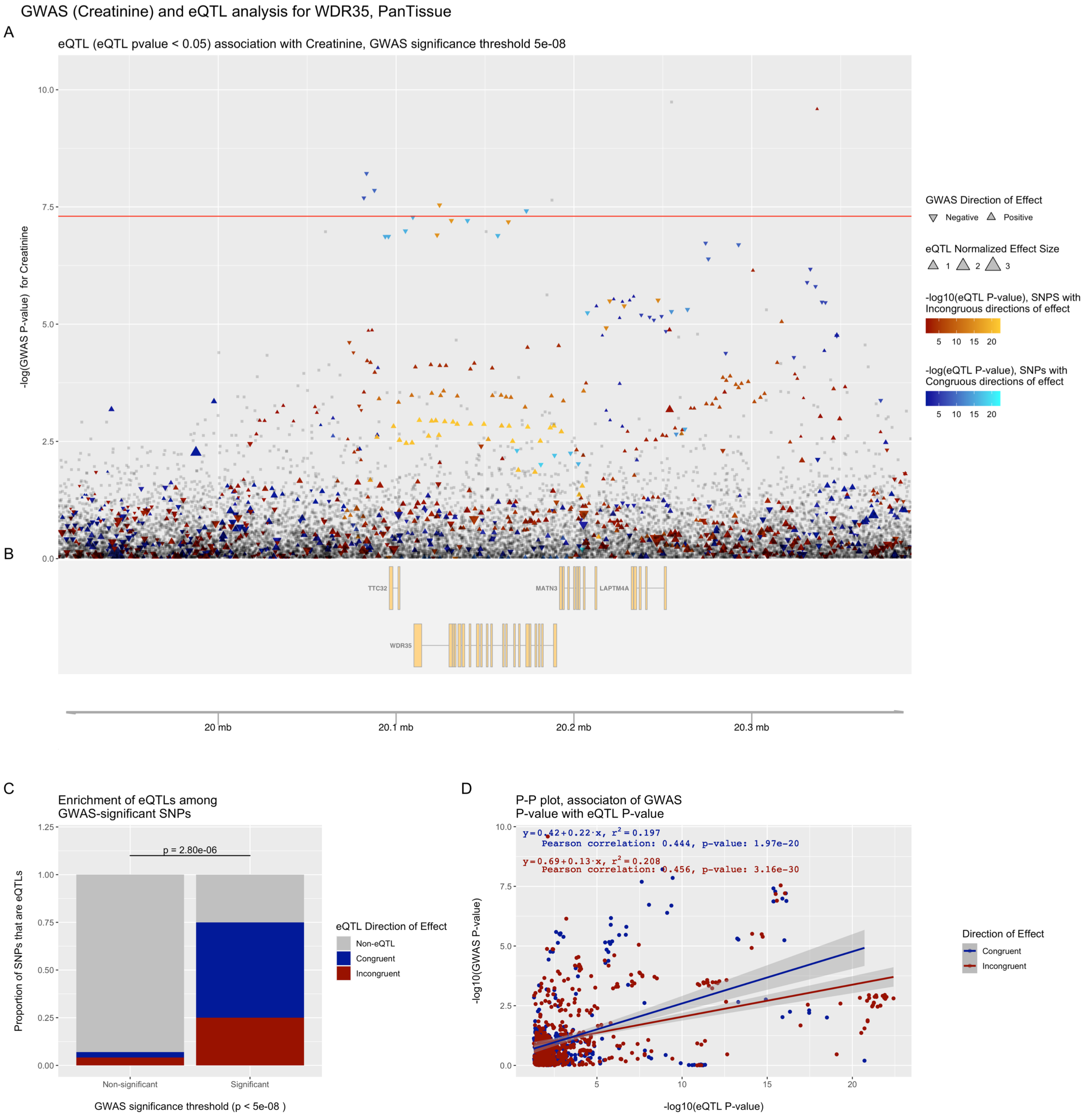


**Figure S34. eQTpLots analysis of the *WDR35* locus and all significant associated phenotypes**

For each phenotype: (A) Plot illustrating any potential colocalization between phenotype-significant variants and eQTLs for the given gene (B) A depiction of the genomic region surrounding the locus of interest (C) Enrichment of eQTLs for the given gene among trait-significant variants. The p-value of enrichment was determined by Fisher's exact test (D) P-P plot illustrating correlation between $p_{eQTL}$ and $p_{trait}$ for the gene of interest. Correlation between the two probabilities is visualized by plotting a best-fit linear regression over the points, with the line equation displayed on the plot. The Pearson correlation coefficient and p-value of correlation are displayed on the plot as well.

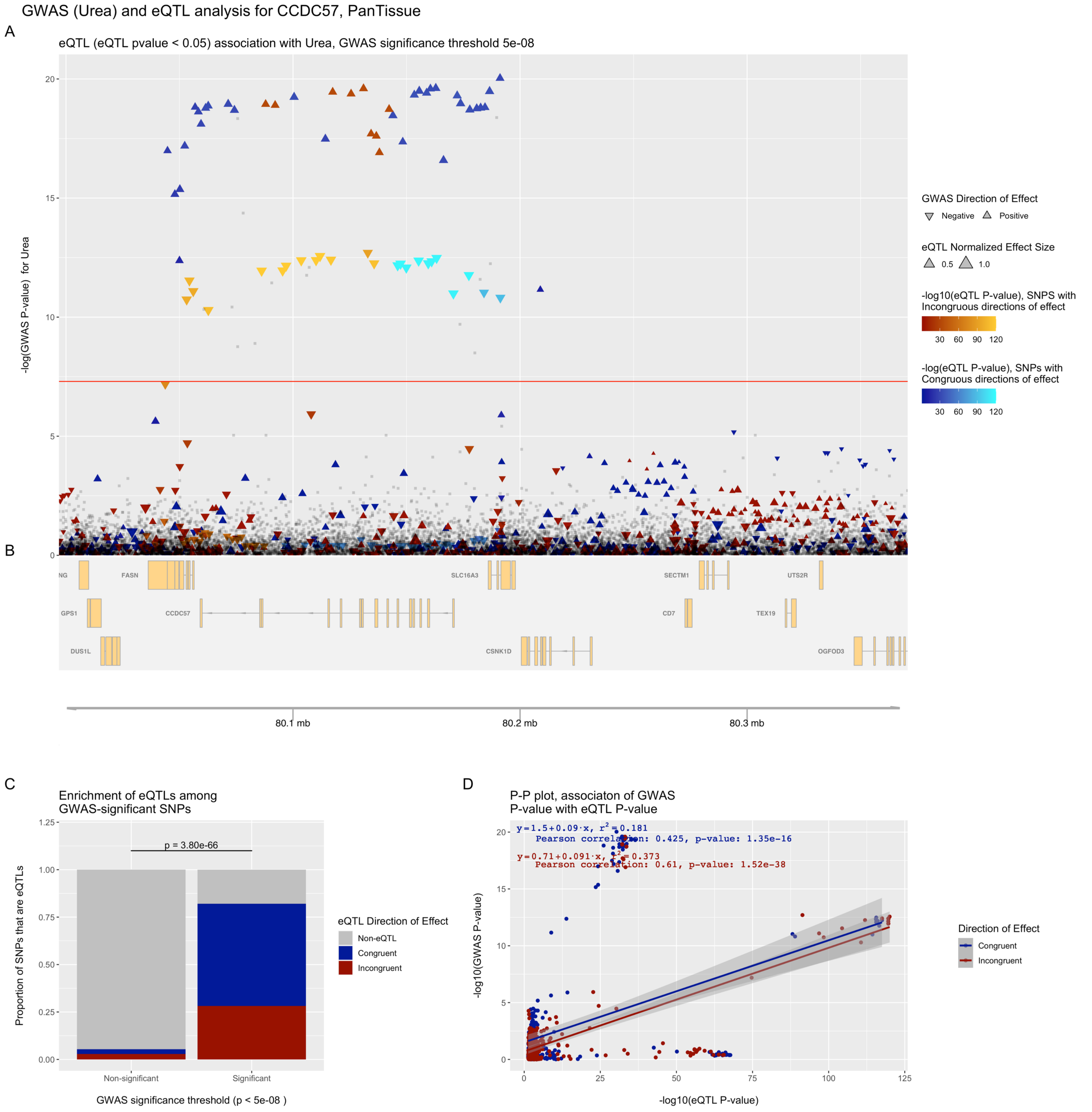


**Figure S35. eQTpLots analysis of the *CCDC57* locus and all significant associated phenotypes**

For each phenotype: (A) Plot illustrating any potential colocalization between phenotype-significant variants and eQTLs for the given gene (B) A depiction of the genomic region surrounding the locus of interest (C) Enrichment of eQTLs for the given gene among trait-significant variants. The p-value of enrichment was determined by Fisher's exact test (D) P-P plot illustrating correlation between $p_{eQTL}$ and $p_{trait}$ for the gene of interest. Correlation between the two probabilities is visualized by plotting a best-fit linear regression over the points, with the line equation displayed on the plot. The Pearson correlation coefficient and p-value of correlation are displayed on the plot as well.

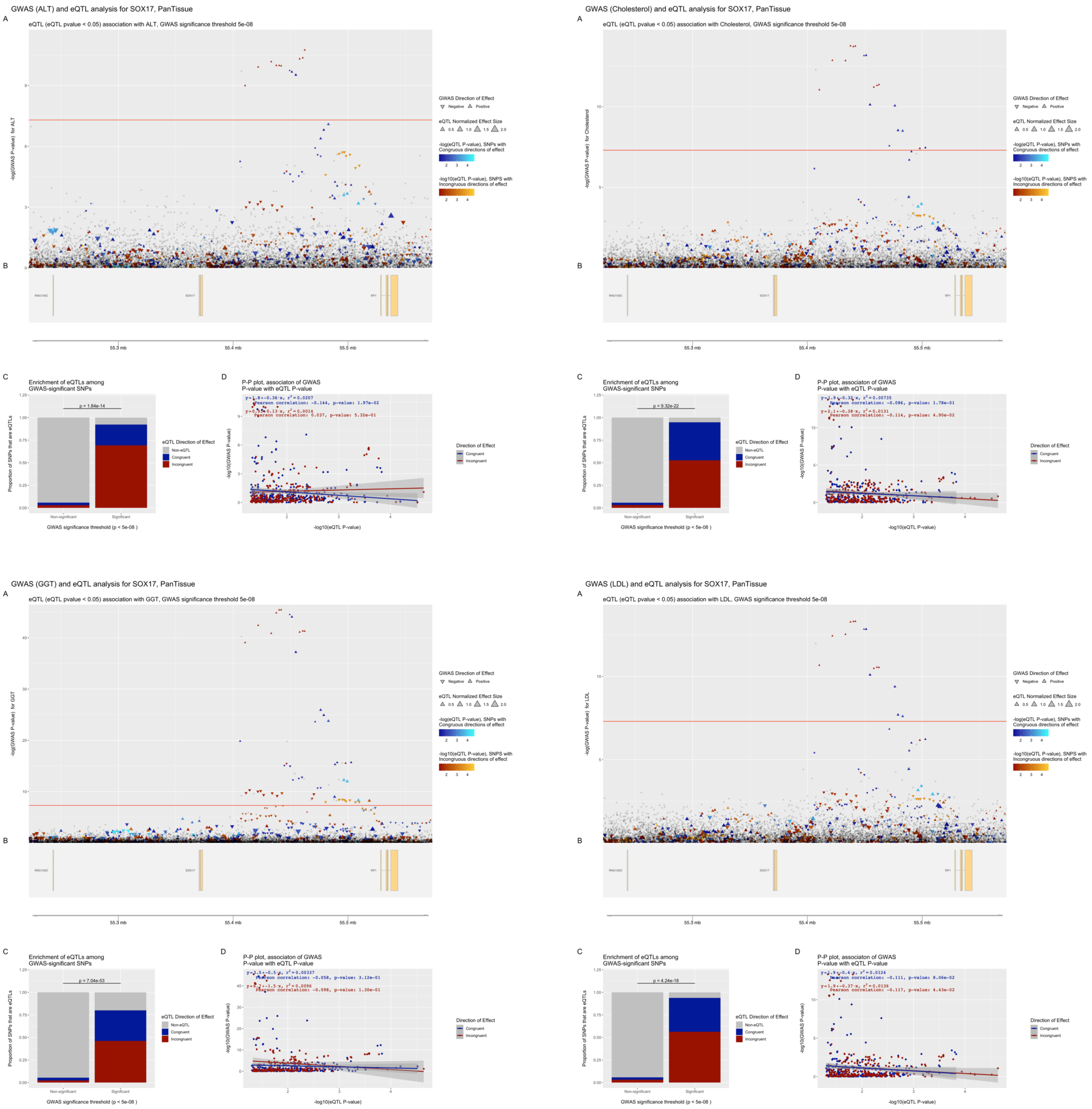


**Figure S36. eQTpLots analysis of the *SOX17* locus and all significant associated phenotypes**

For each phenotype: (A) Plot illustrating any potential colocalization between phenotype-significant variants and eQTLs for the given gene (B) A depiction of the genomic region surrounding the locus of interest (C) Enrichment of eQTLs for the given gene among trait-significant variants. The p-value of enrichment was determined by Fisher's exact test (D) P-P plot illustrating correlation between $p_{eQTL}$ and $p_{trait}$ for the gene of interest. Correlation between the two probabilities is visualized by plotting a best-fit linear regression over the points, with the line equation displayed on the plot. The Pearson correlation coefficient and p-value of correlation are displayed on the plot as well.

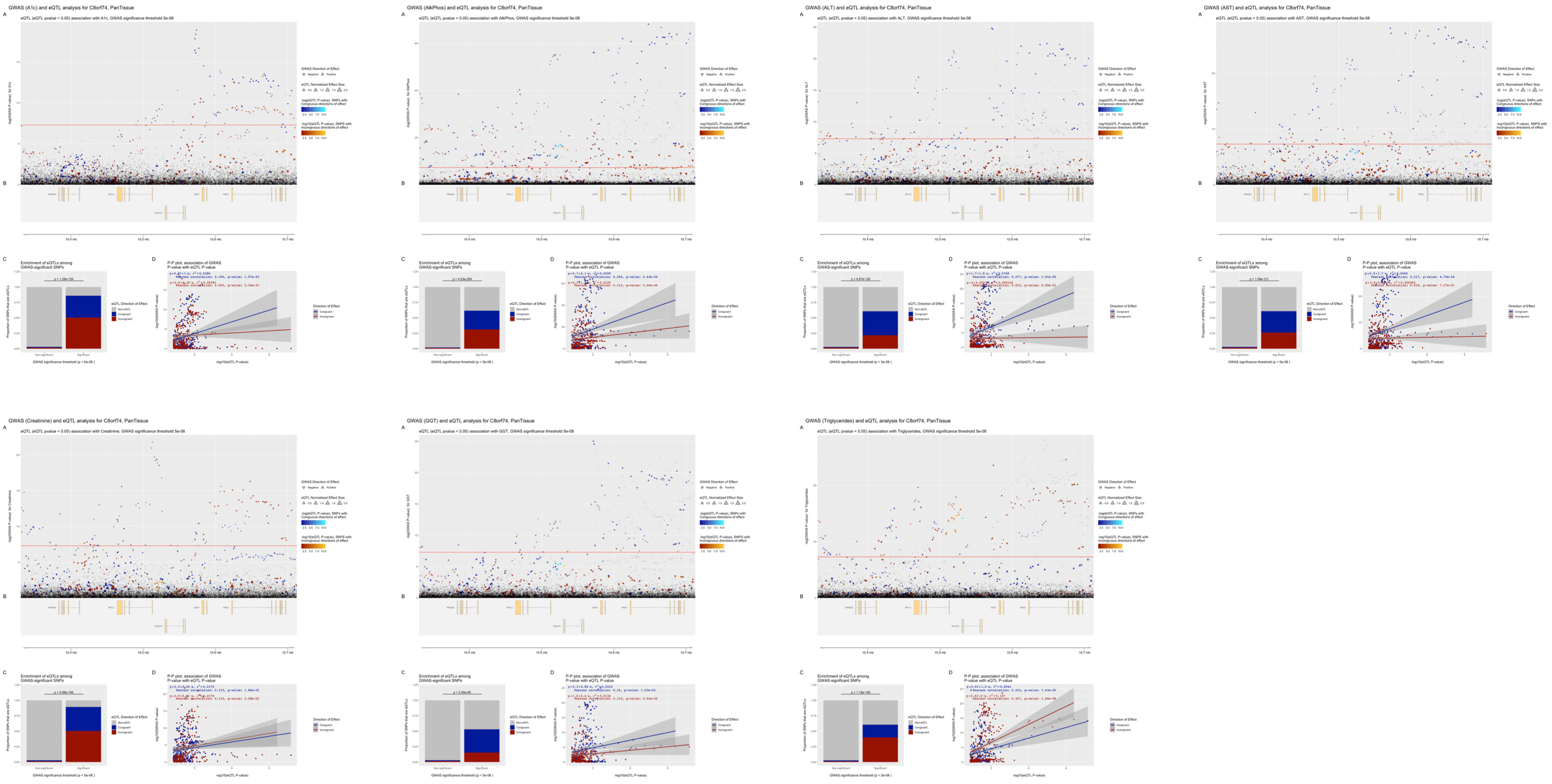


**Figure S37. eQTpLots analysis of the *C8orf74* locus and all significant associated phenotypes**

For each phenotype: (A) Plot illustrating any potential colocalization between phenotype-significant variants and eQTLs for the given gene (B) A depiction of the genomic region surrounding the locus of interest (C) Enrichment of eQTLs for the given gene among trait-significant variants. The p-value of enrichment was determined by Fisher's exact test (D) P-P plot illustrating correlation between $p_{eQTL}$ and $p_{trait}$ for the gene of interest. Correlation between the two probabilities is visualized by plotting a best-fit linear regression over the points, with the line equation displayed on the plot. The Pearson correlation coefficient and p-value of correlation are displayed on the plot as well.

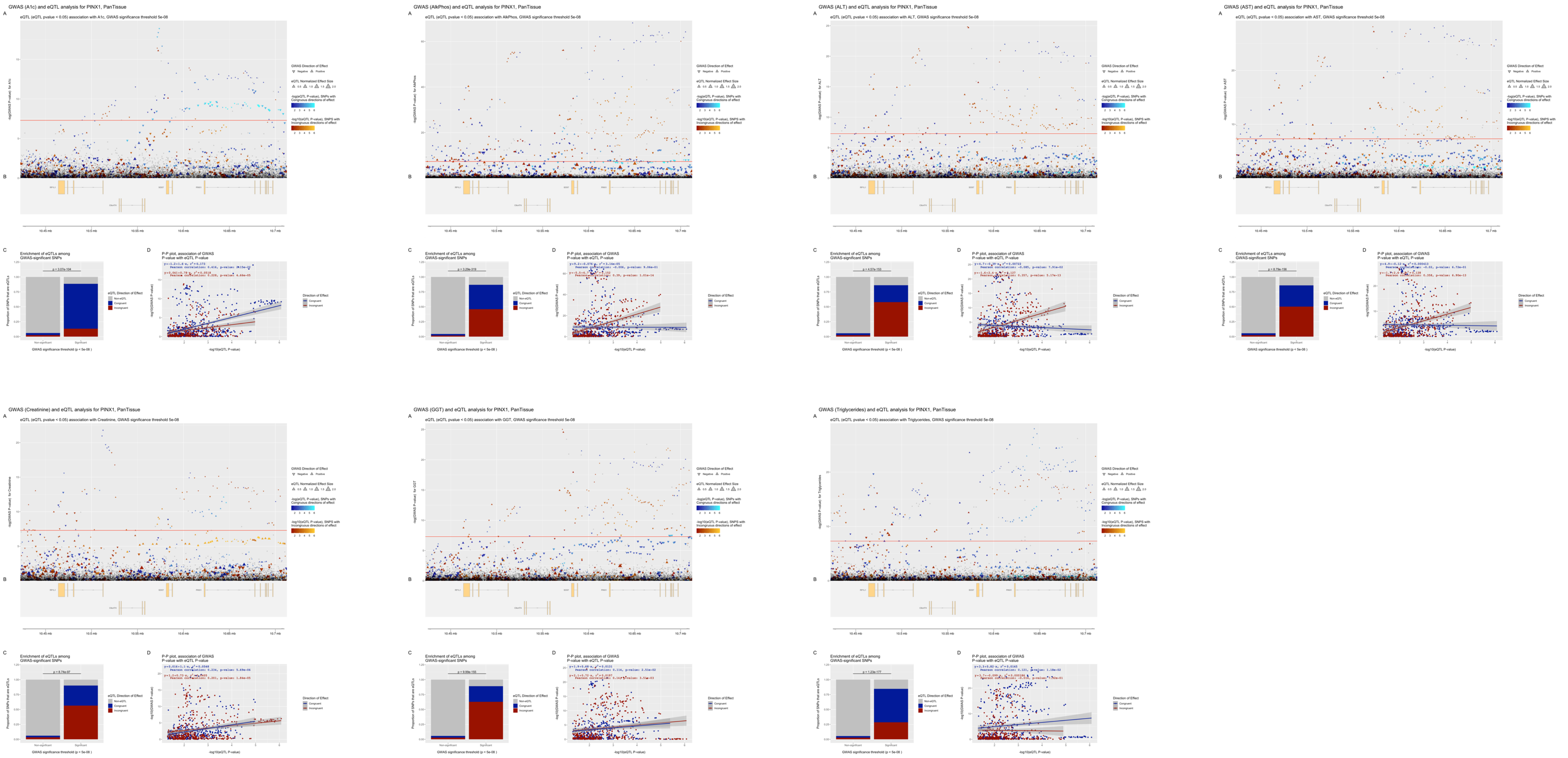


**Figure S38. eQTpLots analysis of the *PINX1* locus and all significant associated phenotypes**

For each phenotype: (A) Plot illustrating any potential colocalization between phenotype-significant variants and eQTLs for the given gene (B) A depiction of the genomic region surrounding the locus of interest (C) Enrichment of eQTLs for the given gene among trait-significant variants. The p-value of enrichment was determined by Fisher's exact test (D) P-P plot illustrating correlation between $p_{eQTL}$ and $p_{trait}$ for the gene of interest. Correlation between the two probabilities is visualized by plotting a best-fit linear regression over the points, with the line equation displayed on the plot. The Pearson correlation coefficient and p-value of correlation are displayed on the plot as well.

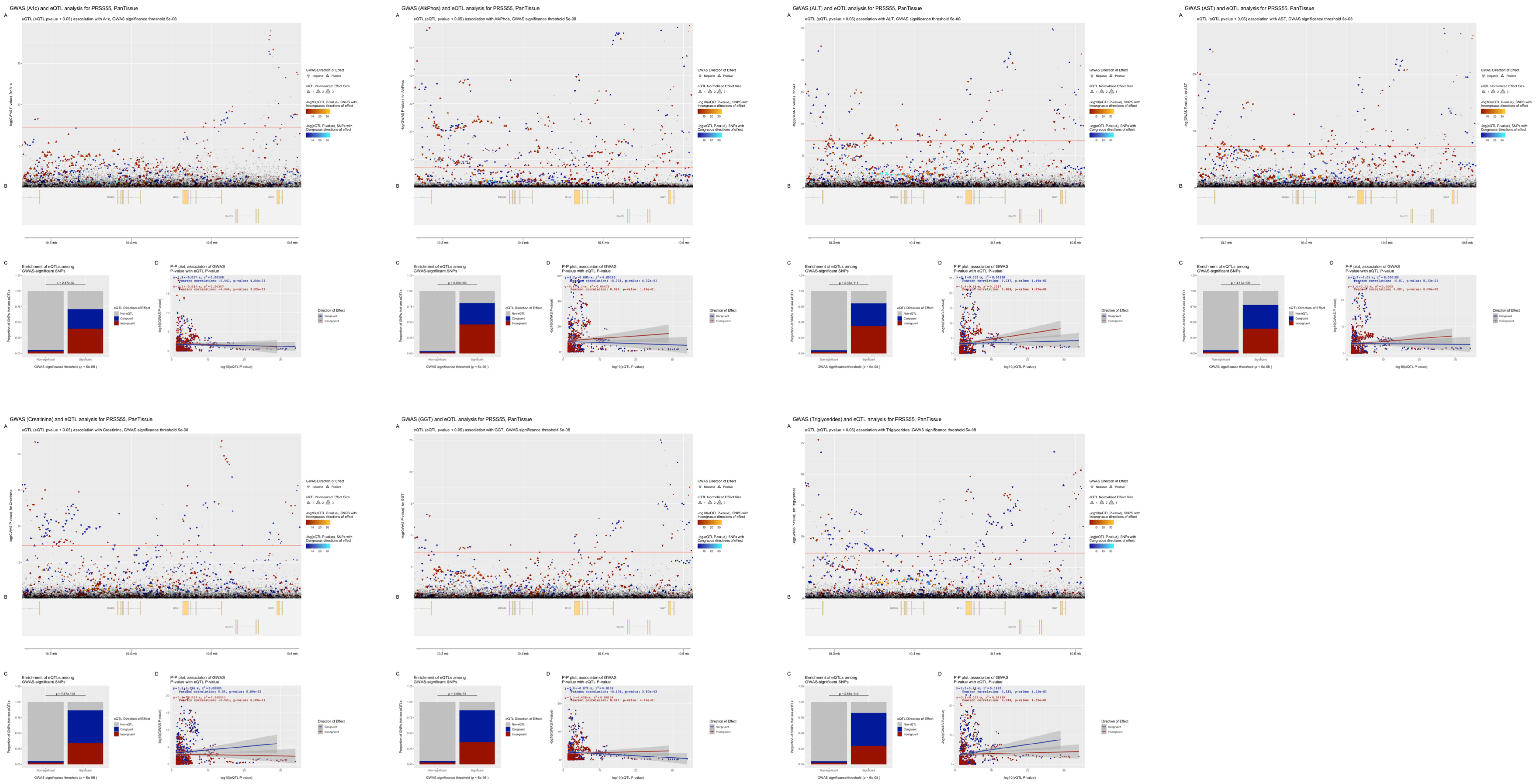


**Figure S39. eQTpLots analysis of the *PRSS55* locus and all significant associated phenotypes**
For each phenotype: (A) Plot illustrating any potential colocalization between phenotype-significant variants and eQTLs for the given gene (B) A depiction of the genomic region surrounding the locus of interest (C) Enrichment of eQTLs for the given gene among trait-significant variants. The p-value of enrichment was determined by Fisher’s exact test (D) P-P plot illustrating correlation between $p_{eQTL}$ and $p_{trait}$ for the gene of interest. Correlation between the two probabilities is visualized by plotting a best-fit linear regression over the points, with the line equation displayed on the plot. The Pearson correlation coefficient and p-value of correlation are displayed on the plot as well.

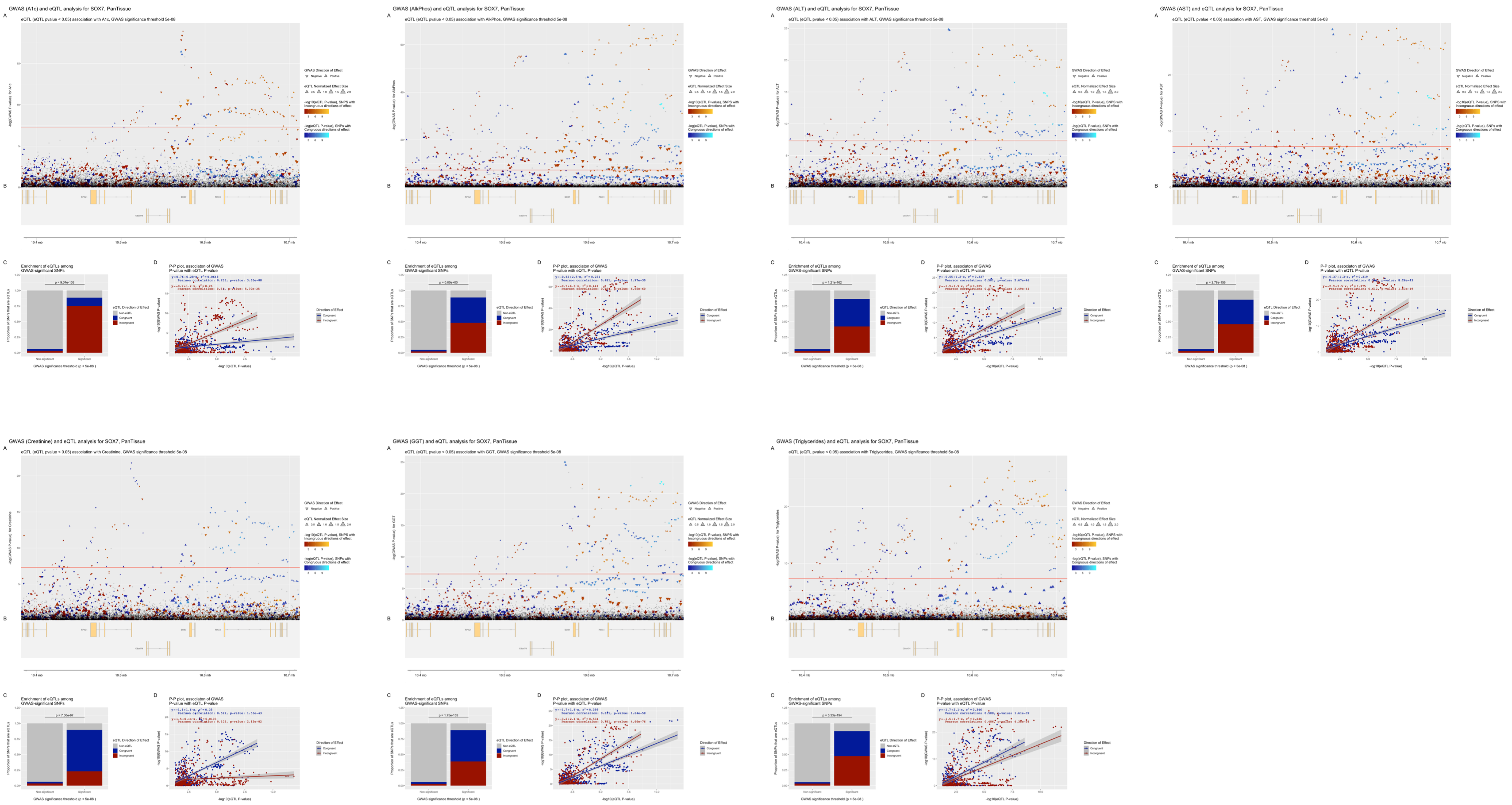


**Figure S40. eQTpLots analysis of the *SOX7* locus and all significant associated phenotypes**

For each phenotype: (A) Plot illustrating any potential colocalization between phenotype-significant variants and eQTLs for the given gene (B) A depiction of the genomic region surrounding the locus of interest (C) Enrichment of eQTLs for the given gene among trait-significant variants. The p-value of enrichment was determined by Fisher's exact test (D) P-P plot illustrating correlation between $p_{eQTL}$ and $p_{trait}$ for the gene of interest. Correlation between the two probabilities is visualized by plotting a best-fit linear regression over the points, with the line equation displayed on the plot. The Pearson correlation coefficient and p-value of correlation are displayed on the plot as well.

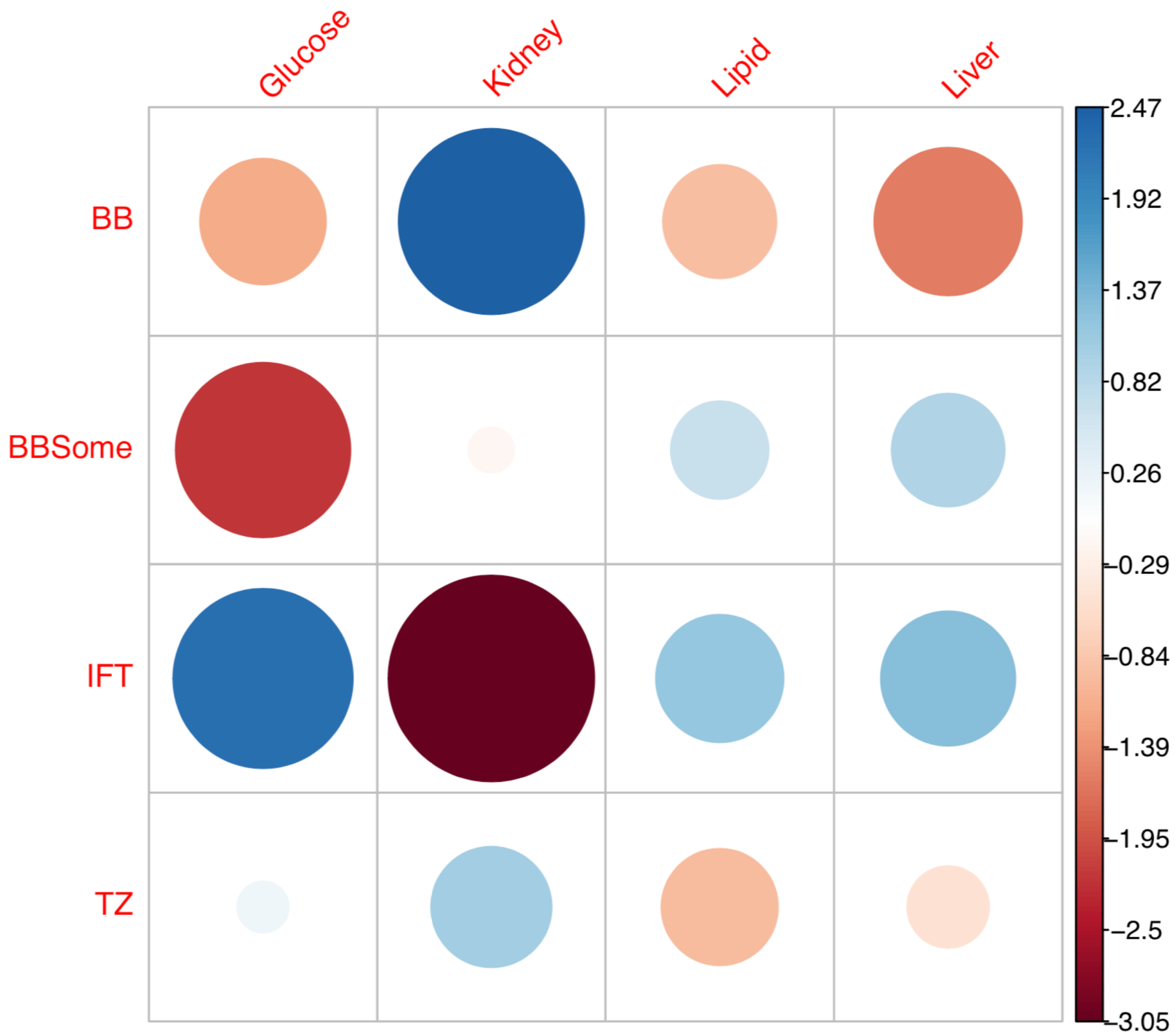


**Figure S41. Visualization of Pearson residuals for chi square test of DiCE Contingency table**
Residuals of the chi square test of the DiCE Contingency table shown in Table S16 are displayed. The size of each bubble reflects the magnitude of the Pearson residual for chi-square analysis for each cell. Positive residuals are in blue, negative residuals are in red, with a color gradient corresponding to the absolute magnitude of the residuals.

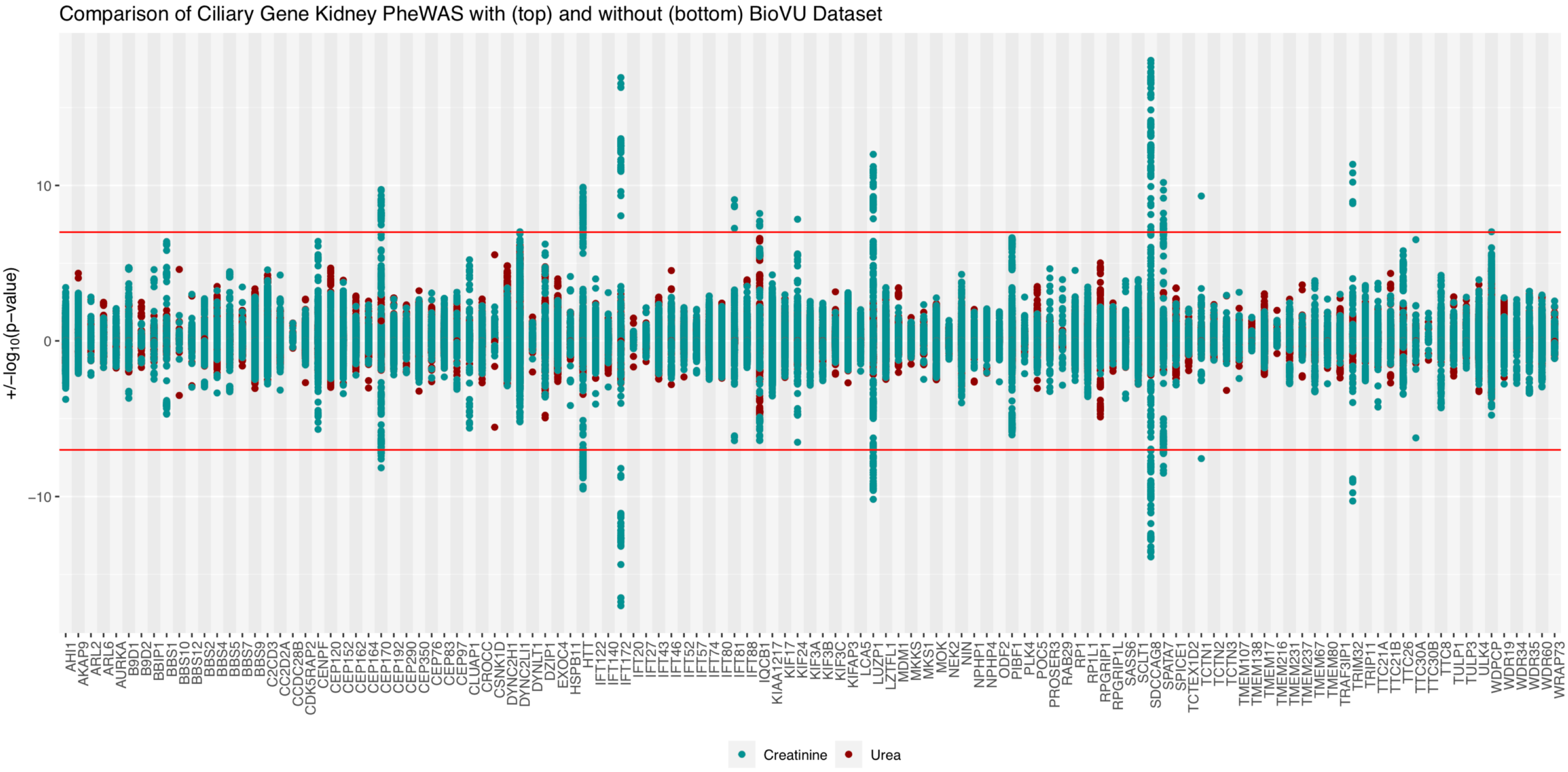


**Figure S42. Comparison of meta-analysis results for kidney-related traits with and without overlapping BioVU datasets**
In our meta-analysis of kidney-related traits, there was an overlap of 6,492 individuals between the large meta-analysis of eGFR and Urea traits (n = 567,460) conducted by Wuttke et al. (which contained samples from an earlier release of the BioVu dataset), and a GWAS performed on the current BioVU dataset (urea n = 33,322, creatinine n = 33,493). To illustrate differences in meta-analysis results including versus excluding this overlapping data, we generated a Hudson plot illustrating the results of the meta-analysis of kidney-related traits including overlapping BioVU samples (top) and without the overlapping BioVU samples (bottom). Data for creatinine is in teal, and data for urea is in red. The genome-wide significance threshold ($p<5e-8$) is shown as a red line. Each analyzed gene is given equal space along the horizontal axis, with all tested genetic variants for a given gene plotted at the midline of the gene block.
